# A Dual Homeostatic Regulation of Dry Mass and Volume Defines a Target Density in Proliferating Mammalian Cells

**DOI:** 10.1101/2025.04.24.650395

**Authors:** Nishit Srivastava, Ludovico Calabrese, Camille N. Plancke, Romain Rollin, Larisa Venkova, Kristina Havas, Marco Cosentino Lagomarsino, Matthieu Piel

**Author notes:** Co-first authors (Nishit Srivastava, Ludovico Calabrese). These authors contributed equally, although differently, to the work (Marco Cosentino Lagomarsino, Matthieu Piel).

## Abstract

The concentration of macromolecules, especially proteins, is vital for cellular function and is influenced not only by synthesis and degradation but also by the total cell volume. While we understand various growth regulation mechanisms, the coupling of dry mass and volume in growing mammalian cells remains unclear. Here we show that two independent mechanisms acting in single cells —one regulating volume through biophysical modulation and the other controlling protein biosynthesis—work together to maintain macromolecular dry mass density and restore it following perturbations. These mechanisms ensure that proliferating cells remain within a specific range around a target density, providing density homeostasis at the population level. Although the target density appears consistent across different cell types, it is disrupted around cell division, upon perturbations of growth pathways and in senescent cells. It may represent an optimal value for cellular processes, ensuring the efficiency of essential intracellular functions.

## Introduction

Cellular concentrations of key macromolecules, such as proteins, are essential for a range of processes, including protein-catalyzed reactions, complex assembly, biosynthesis, phase separation, and polymerization. It is well-established that macromolecular concentrations are tightly regulated within a narrow range, indicating a sophisticated control mechanism beyond the regulation of individual components (*1–10*). While recent research has illuminated various homeostatic feedbacks and buffering mechanisms (*3*, *5*, *10*, *11*), the coupling of cell volume growth with macromolecular content production—and thereby macromolecular concentration—remains poorly understood.

The long-standing pump-leak model, supported by extensive experimental evidence (*12–14*), describes cell volume regulation based on the so-called Donnan equilibrium. According to this model, cell water content is determined by the balance of hydrostatic and osmotic pressures, with the cell maintaining global electroneutrality and balanced ion fluxes. This model provides a formula for cell volume as a function of the number trapped molecules, where small metabolites and their counterions—such as negatively charged amino acids like glutamate paired with potassium—, with a total amount approaching 100 mM in cultured mammalian cells, predominate in influencing cellular volume, by being most abundant (*13*). In contrast, proteins, which account for the majority of macromolecular dry mass, are present at millimolar concentrations. While protein crowding affects the diffusion of large particles (*15*), it has no impact on the entropy of small molecules contributing to the cell’s wet volume. Thus, despite their effect on water potential (*5*), proteins have a minor direct effect on wet volume compared to small osmolytes, as evidenced by the general linearity of Ponder plots for mammalian cells (*16*). This leads to a critical question: how are cell volume growth and macromolecular biosynthesis related, in single cells, given that they depend on different compounds—proteins and small osmolytes, and thus on different intracellular processes?

The regulation of biosynthesis has been extensively studied, revealing various mechanisms of concentration control (*17–23*). High macromolecular density and large complexes, such as ribosomes, are known to influence protein translation and degradation (*3*, *10*), generating feedback mechanisms that regulate protein concentrations. Recent findings suggest that feedbacks may also occur at the transcriptional level (*24*, *25*), through condensate formation affecting enzymatic activity and ion channels (*5*, *26–29*), as well as regulation of dry mass synthesis, potentially involving a modulation of nuclear import/export rates (*11*). These feedback mechanisms, acting over various timescales, maintain tight control over protein concentrations. The physiological importance of this control is highlighted by evidence linking cytoplasmic dilution to ageing, senescence and other diseases (*2*, *30–35*). However, the regulation of volume growth—or water content—remains relatively unexplored, especially in the context of cell growth, leaving the coordination of volume and macromolecular mass an elusive puzzle.

To tackle this “fundamental enigma” in cell physiology, it is crucial to draw a picture that integrates biosynthesis regulation mechanisms with the biophysics of volume control, at the single-cell level. In this study, we investigate how total volume and macromolecular dry mass are dynamically coupled in single cultured proliferating mammalian cells. By combining quantitative methods and mathematical modeling, we show that two homeostatic mechanisms together define a target macromolecular dry mass density, and keep the cell population within a narrow density range, despite noise due to cell division and fluctuating growth.

## Results

We first produced a large and carefully curated dataset of measured macromolecular dry mass, volume and projected area in cultured proliferating cells. To disentangle the effect of cell adhesion and spreading on cell growth, we used two different types of HeLa cells, one growing on adhesive substrates (that we call HeLa adherent) and the other in suspension (HeLa S3), as well as a leukemia derived hematopoietic cell line, also growing in suspension (HL 60). Concomitant measures of volume and macromolecular dry mass were recorded optically as previously described (*12*, *36*), using a combination of fluorescence exclusion (*37*, *38*), quantitative phase imaging (*39*, *40*), and an optimized analysis routine (Fig. 1A and Fig. S1 A-D). We define here dry mass density as the ratio of the two independent measures of macromolecular dry mass and total cell volume. We first plotted the (conditional) population averages along the cell-cycle (Fig. 1 B,C and Fig. S1 E-H). We verified that single cells in all the cell lines were best described as following robust exponential growth in both macromolecular dry mass and volume (Fig. S1 I,J for HeLa adherent) with mean specific growth rates of about 0.03 h^-1^ (Fig. 1D,E for HeLa adherent), that were on average equal for macromolecular dry mass and in volume and corresponded to a doubling time of about 20 hours. Figure 1D shows conditional averages across the cell population of the mass and volume growth rates along the cell cycle. As expected from previous studies, due to the slowdown of dry mass/protein production during mitosis (*36*, *41*) and mitotic swelling, mass and volume growth rates differ in mitosis, leading to cell dilution (*36*, *42*) (Fig. 1F, G, S1 K-O). At mitotic exit, cell volume rapidly decreased as adherent daughter cells were respreading, before resuming steady growth at the end of the spreading phase (EOS, Fig. 1F, S1 L,M), that last in average about 2.5 hours after birth (Fig. S1 P). Surprisingly, at mitotic exit, mass production immediately resumed at a much higher rate than volume, leading to a rapid increase in dry mass density (Fig. 1F left part, S1 K). Within a few hours after mitosis, both dry mass and volume growth rates converged to the same average value and dry mass density stabilized to a value similar to its pre mitotic value (Fig. 1F, S1 N). Overall, this analysis showed that cultured human cells grow exponentially in dry mass and volume at similar rates during the bulk of interphase, and maintain their dry mass density distribution despite large variations in mitosis.

**Figure 1.**
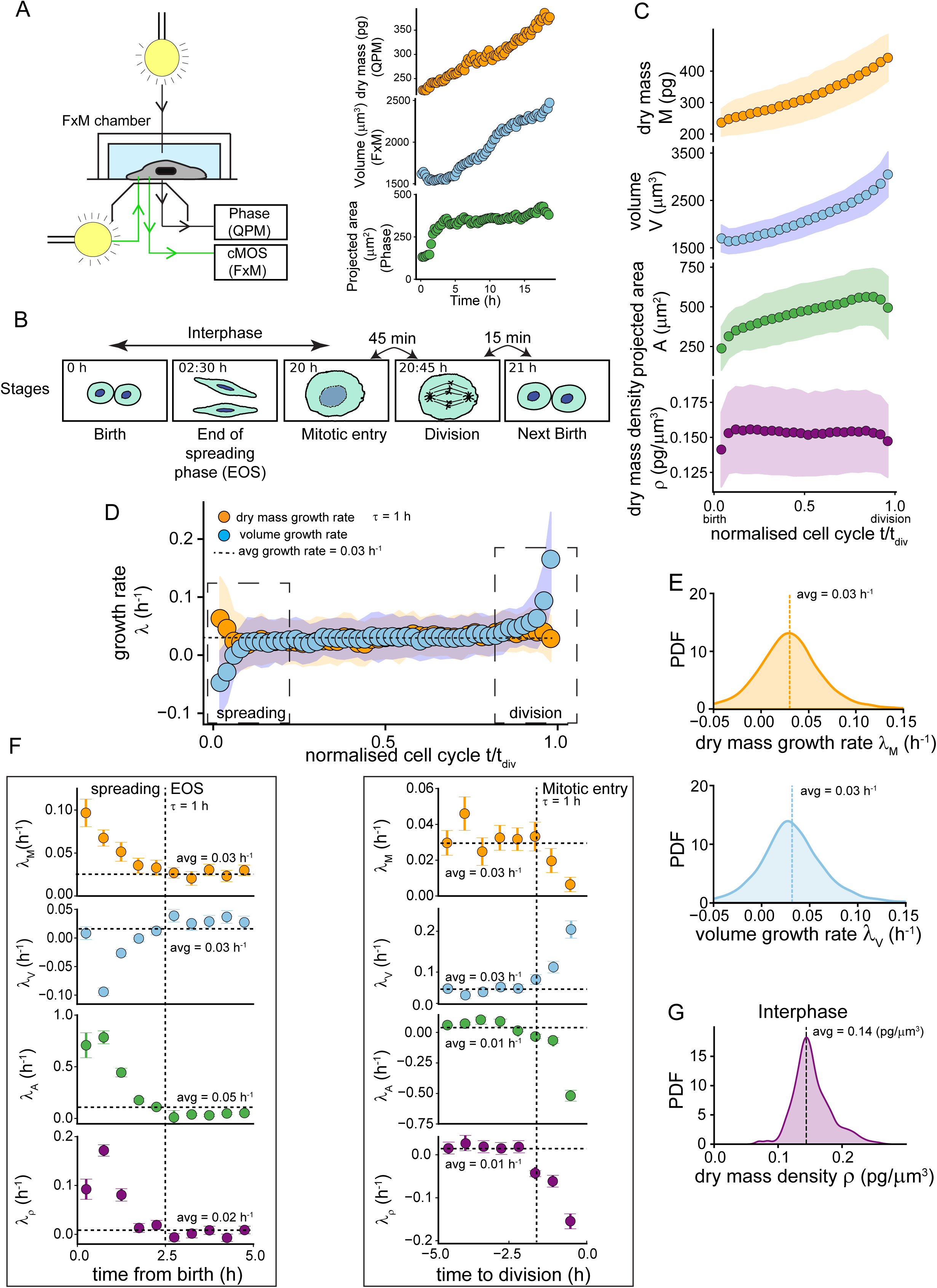
Single cells dry mass and volume trajectories from birth to division. **(A)** Schematic of the acquisition setup (left). Example of a single live cell trajectory in dry mass (Quantitative Phase Microscopy, QPM, top right), volume (Fluorescence Exclusion Microscopy, FXm, middle right) and projected area (phase contrast); t=0 is birth **(B)** Schematic defining the cell cycle stages used as reference points in our data analysis (see methods for details); **(C)** From top to bottom: dry mass (M, orange), volume (V, blue), projected area (A, green) and dry mass density (ρ=M/V, purple) as a function of the normalized cell cycle time (t/t_div_, t_div_ time of division), with t/t_div_ = 0 for birth and t/t_div_ =1 for division **(D)** Conditional averages of growth rate in dry mass (*λ*_M_= 1/M dM/dT, orange) and volume (*λ*_V_= 1/V dV/dT, blue) calculated over 1 hour windows (see methods), against normalized cell cycle time; **(E)** Probability density function (PDF) of dry mass growth rate (left, orange) and volume growth rate (right, blue) **(F)** Zoom on the data shown in D for the first 5 hours after birth (left) and the 5 hours before division (right), for dry mass growth rate (orange), volume growth rate (volume), area growth rate (*λ*_A_, green) and density growth rate (*λ*_ρ_, purple); **(G)** PDF of the dry mass density of all the cells across all the timepoints during interphase (birth to mitotic entry). The dataset used in the entire figure is for HeLa adherent cells (n=133 cells, N=3 experiments). C,D,F: binned averages +/− SEM

To understand how growth rates stabilized in post mitotic cells, we analyzed the behavior of dry mass density time course in single cells, focusing first on HeLa adherent cells (Fig. 2A). We found that, at mitotic exit and during the spreading phase, an initial loss of volume while dry mass growth has already restarted leads, in a large majority of cells, to an increase in the dry mass density (2B, S1 K-N). Single cells then reach different density values, below or above the mean dry mass density they had prior to mitosis (Fig. 2B, S2 A) suggesting that dry-mass density is greatly perturbed around cell division. Consequently, the dispersion of dry mass density (measured by the coefficient of variation, CV) is larger in the cell population at the end of spreading (EOS) compared to before mitosis (mitotic entry) and birth, suggesting that while the post-mitotic phase perturbs density, this perturbation gets progressively corrected (Fig. 2C). The different post-mitotic trends of dry mass and volume during the spreading phase and during mitosis suggest that dry mass and volume depend on different processes (Fig. 1F, S1 K-N). To quantify their coupling, we compared the CV of the dry mass density (which is the ratio of dry mass and volume) and the sum of the CVs in dry mass and in volume, defining a “coupling index” as the difference between the two (Fig. 2D). We found that, as expected for parameters controlled by different processes, the coupling was almost zero when considering cells between birth and EOS. On the other hand, when considering post-EOS cells, the coupling was significantly higher for all cell types, and stronger in cells growing in suspension (Fig. 2D). To assess the coupling at the single-cell level, we plotted the two growth rates against each other (measured in single cells by specific time derivatives over 2 hours), and performed a conditional average to extract the trend. The time-conditional average of the mass growth rate was on average equal to the volume growth rate (Fig. 1D) for post-EOS cells, while it was not the case for mitotic cells (Fig. S2B). Nevertheless, when considering the deviation of growth rates from their mean value, for single cells at a given time, it appears that the two growth rates can take very different values (Fig. 2E), even if they are on average equal, at the population level and on long enough time scales (Fig. 1D, Fig. 2E). Together, these analyses suggest the existence of homeostasis mechanisms that ensure the coupling of dry mass and total volume growth rates after EOS, ensuring equal average growth in mass and volume despite largely independent fluctuations.

**Figure 2.**
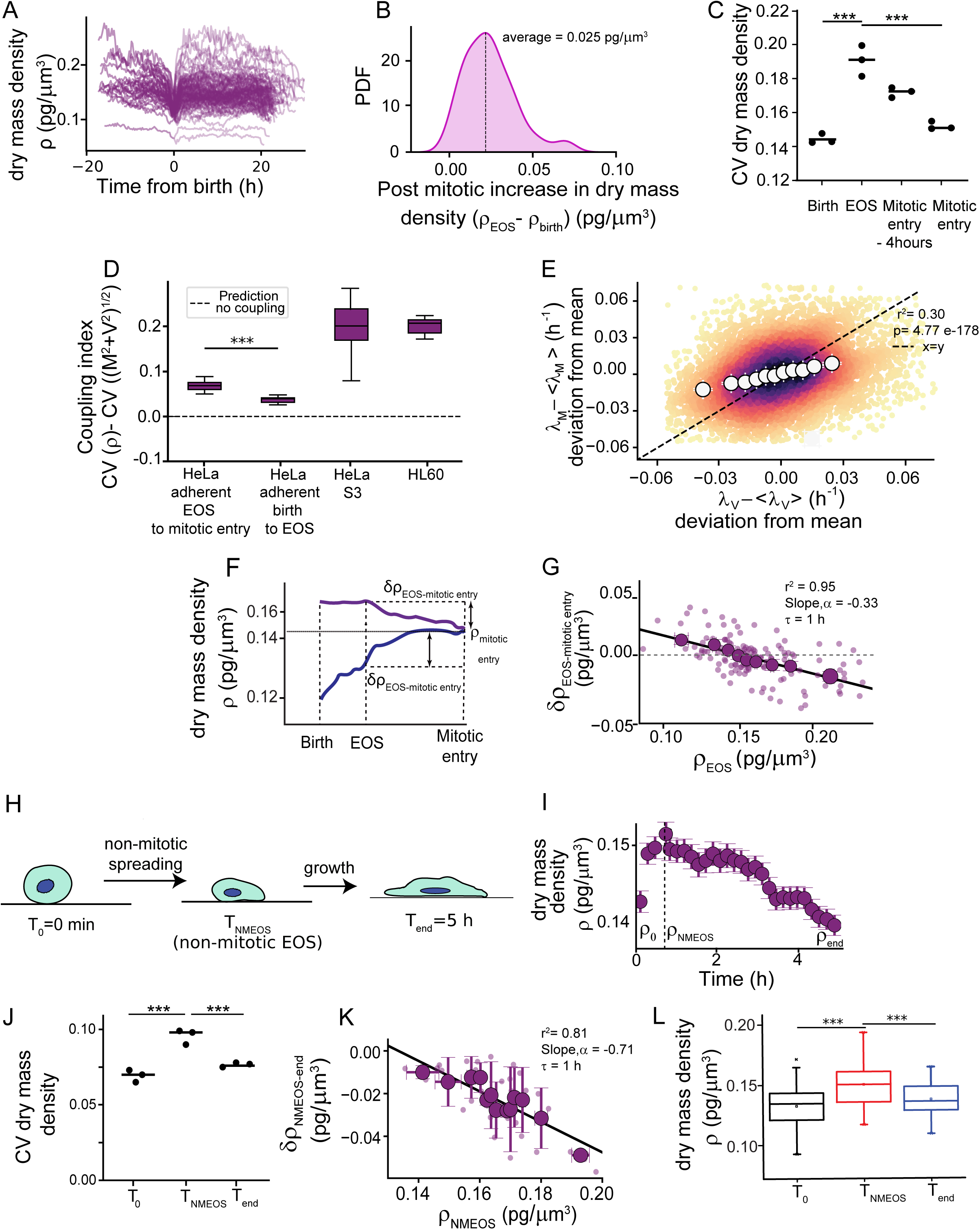
Evidence for dry mass homeostasis with hours timescale correction of large perturbations. **(A)** Dry mass density trajectories for individual cells centered around birth (t=0,birth), t<0 is for the mother cell and t>0 for the two daughters **(B)** Distribution of the magnitude of change in dry mass densities between birth (ρ_birth_) and EOS (ρ_EOS_) in single cells **(C)** Coefficient of variation (CV) of dry mass density at birth, EOS, 4 hours before mitotic entry and mitotic entry (each point corresponds to an independent experiment,*** p<0.005) **(D)** Dry mass and volume coupling index (see methods for details; briefly, it is defined for each single cell, has a positive value, not different from zero when the two variables are not coupled and increases for stronger coupling). Coupling strength of HeLa adherent cells between birth and EOS is not different from zero (p= 0.8), while it is for EOS to next mitotic entry (*** p<0.005) **(E)** Heatmap scatter plot of deviations in mass growth rate versus volume growth measured as deviation of growth rates from their single-cell mean values calculated during the bulk phase of the cell cycle (darker colors indicate higher density of data, discrete derivatives measured over 2 hours). Binned averages are shown as white circles. **(F)** Schematic explaining, on two single cells dry mass density trajectories, the measure of dry mass density changes from EOS to mitotic entry (ρ_EOS-mitotic entry_). Cell1 (purple) is above average at EOS and decreases (ρ_EOS-mitotic entry_<0), while Cell 2 (blue) is lower at EOS and increases (ρ_EOS-mitotic entry_>0) **(G)** Dry mass density correction (8ρ) from EOS to mitotic entry versus dry mass density at EOS. **(H)** Schematic of the dynamic spreading experiment (see methods; in brief, cells display rapid spreading for about 30 minutes, a timepoint termed NMEOS for Non-Mitotic End of Spreading, calculated for each cell based on the spreading area trajectory) **(I)** Dry mass density plotted as binned average ± SEM from T_0_ to T_end_. Dashed line shows average NMEOS time **(J)** CV of dry mass density at T_0_,T_NMEOS_, T_end_ (*** p<0.005) **(K)** Dry mass density correction (8ρ) between NMEOS and T_end_ against dry mass density at NMEOS; **(L)** Box plots of the dry mass density distribution at the main timepoints of the spreading experiment (*** p<0.005). G and K: small circles are single cell data and large ones are binned average ± SEM; solid line is a linear fit on the binned averages (slope and goodness of fit, r^2^ are shown); for all the plots in this figure, solid circles are binned mean values calculated on equally weighted bins weighted on the number of observations in each bin. A-C and E-H: HeLa adherent cells (n=133 cells, N=3 independent experiments). Panel D is based on the full dataset for the 3 cell lines (HeLa adherent: n=133, N=3; HeLa S3: n=140, N=3; HL 60: n=115, N=2); I-M: HeLa adherent cells (n=54 cells, N=3 independent experiments).

To test whether the coupling of growth in dry mass and total volume is strong enough to ensure a global dry mass density homeostasis at the cell cycle scale, we performed an analysis on single-cell tracks, comparing the change in dry mass density between EOS and mitotic entry with the initial dry mass density at EOS (Fig. 2F,G). This analysis, which is impervious to possible individual dry mass density setpoints in different cells, showed a clear homeostatic trend, more diluted cells increased their dry mass density while denser cells decreased it, meaning that cells corrected both the positive and the negative perturbations of dry mass density that they faced following mitosis. This trend was absent during mitosis (mitotic entry to division) and between birth and EOS (Fig. S2 C,D). Dry mass density correction could also be observed between the start and end of single cells growth trajectories for all three cell lines we studied (Fig. S2 E-G), when considering windows of 4 to 10 hours outside the region of the cell cycle where density is perturbed. Previously, we and others identified two main sources of dry mass density perturbation in proliferating cultured cells: mitotic swelling (*36*, *42*) and cell spreading (*12*), both phenomena occurring one after the other in dividing cells. To further test the robustness of the density homeostasis mechanism experimentally, we induced a non-mitotic density perturbation (Fig. 2H). Based on previous work (*12*, *43*), we know that spreading cells loose volume, while they keep growing in mass (Fig. S2 H-J). Similar to post mitotic spreading, non-mitotic spreading induced an increase in dry mass density (Fig. 2I). During the post spreading steady-state growth regime, dry mass density decreased at a slow pace until it eventually reached its pre-spreading level, after several hours of growth (Fig. 2I), indicative of a correction mechanism similarly to what was observed during post-mitotic steady-state growth. Because cells spreading at different speeds lost different amounts of volume, as reported before (*12*), the cells at the end of the fast-spreading phase displayed a larger CV of dry mass density than before spreading (Fig. 2J). After spreading stopped, cells resumed steady growth. Comparing single cell changes in density after 5 hours of growth showed that cells that were denser at the end of the spreading phase showed a higher decrease in density (Fig. 2K). This demonstrates that density homeostasis is constantly working in single cells to correct perturbations, independently of mitosis. Therefore, 5 hours after spreading stopped, not only the CV of dry mass density, but also the average dry mass density of the cell population, were back to their pre-spreading values (Fig. 2J,L). Together, these experiments confirm that dry mass density is corrected upon perturbations, at the hour timescale, suggesting corrections acting via modulation of growth rates. In conclusion, dry mass and volume growth rates are distinct but coupled quantities, which leads to a homeostatic behavior of dry mass density acting throughout the cell cycle.

A simple explanation for the coupling between dry mass and volume growth could be that volume increase is a direct consequence of production of dry mass, while the two quantities show independent fluctuations on short timescales due to noise in their regulatory pathways. If this were the case, stopping growth in dry mass should rapidly lead to an arrest of volume growth. We tested this hypothesis by acutely treating cells with cycloheximide (CHX) at a concentration that stops dry mass increase (Fig. S3 A), and recorded mass and volume in single cells (Fig. 3A). About 30 minutes after the treatment, while mass growth had completely stopped, volume kept increasing at an even higher rate than before the treatment (Fig. 3B,C). Volume growth rate remained at a high value for several hours (Fig. 3B), leading to a large dilution of the CHX treated cells (Fig. 3D), before starting to decrease and finally stopping more than 4 hours after the growth arrest in dry mass (Fig. 3E). A similar result was obtained when decreasing mass growth rate with inhibitors of the mTORC pathway, which were also used at a concentration that arrested growth in dry mass (Torin and Rapamycin, Fig. S3 B-E), ruling out some indirect effect of CHX on cell volume. Because water fluxes are known to equilibrate osmotic pressure across the cell surface in the minute timescale (*13*), these results rule out a simple direct coupling between volume and dry mass and demonstrate experimentally that, as previously proposed from a theoretical point of view (*14*), volume and macromolecular dry-mass growth, although coupled, are driven by largely distinct mechanisms.

**Figure 3.**
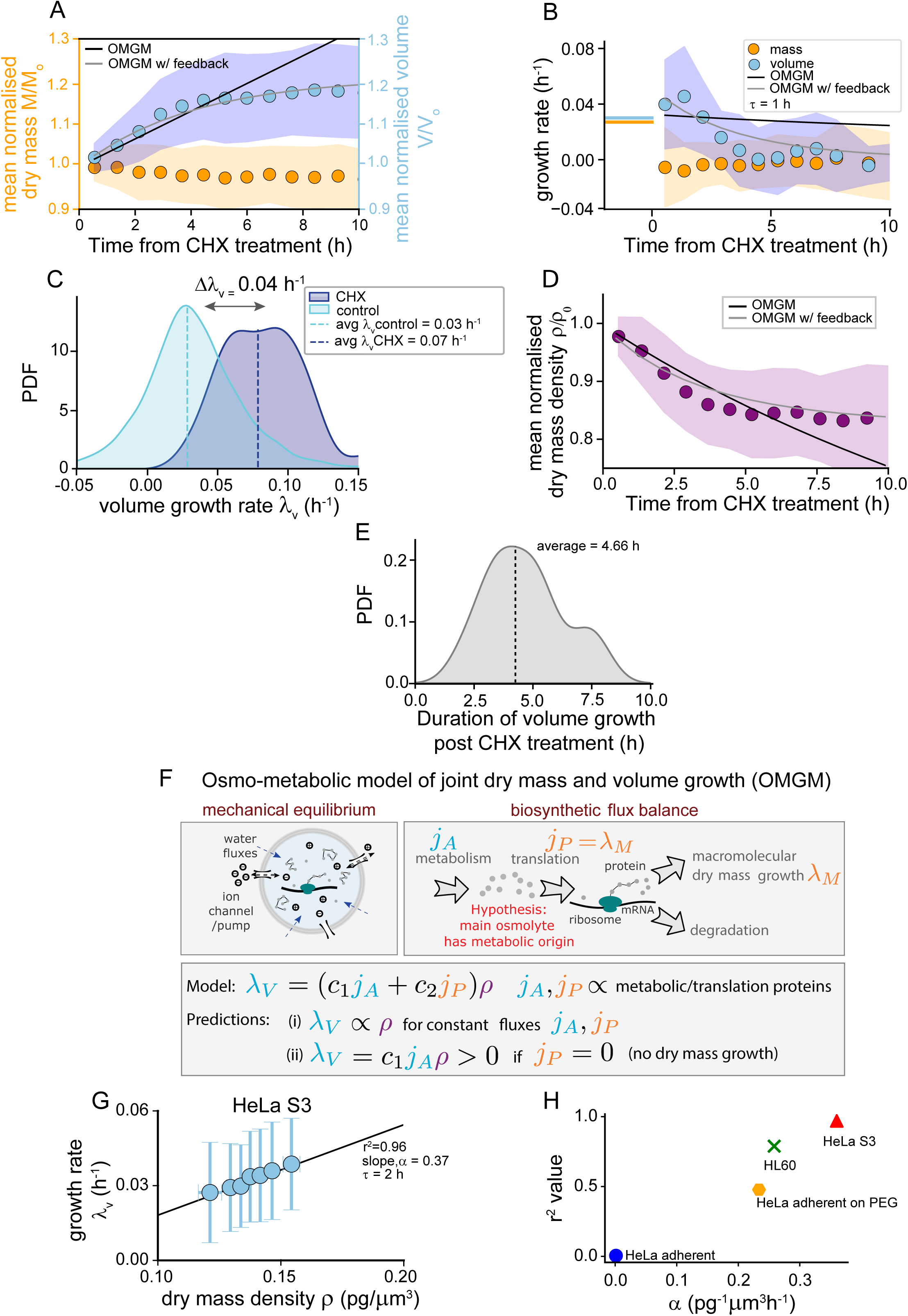
An osmo-metabolic model explains volume growth as a function of dry mass density. **(A-E)** HeLa adherent cells treated with 1μm CHX at time 0; **A:** experimental data (circles) and model fits (black and grey lines) for volume (blue) and dry mass (orange); **B:** dry mass and volume growth rates (discrete time derivatives over 1=1 hour); **C:** Distribution of maximum volume growth rate after CHX treatment (dark blue) compared to that of control cells without CHX treatment (light blue). Δλv is the difference between the mean of the two distributions; **D:** Dry mass density temporal evolution after dry mass growth arrest; **E:** Distribution of time delay between growth arrest in dry mass and in volume; **(F)** Schematic describing the osmo-metabolic growth model (OMGM, see Supplementary information file for details for the two versions of the model) where trapped osmolytes (e.g. metabolites such as free amino acids) could contribute to growth in volume as well as in dry mass (e.g. proteins) thereby acting as coupling agents; j_A_ is the rate of accumulation of trapped metabolites, j_P_ is the rate of protein accumulation; **(G)** Average volume growth rate (𝜆_v_) conditioned to dry mass density (π), discrete derivatives over 2 hours. **(H)** Goodness of fit (r^2^) against slope of linear regressions (α) on binned averages of volume growth rate against dry mass density for all cell lines and culture conditions. A-E: HeLa adherent cells treated with CHX (n=56, N=3), except C light blue, control steady growing HeLa adherent cells shown in Figures 1 and 2; A, B, D: black line are model prediction while grey line are incorporating a feedback to account for volume growth arrest dynamics (see Supplementary Information file for details on the model); G: HeLa S3 cells steady state growth, as in Fig. 2 (n=115, N=3, linear regression p = 0.0015); H: all steady state growth datasets for the various cell lines and conditions, as in Fig. 2. For all graphs, circles show binned averages of equal weighted bins; A, B, D: colored regions are ± SEM, G: binned average ± SD; solid line is a linear fit on the binned averages (slope and goodness of fit, r^2^ are shown).

To understand how the two distinct processes of growth in dry mass and volume could be coupled to ensure dry mass density homeostasis, we turned to mathematical modelling. We figured that the observed decoupling must be due to the processes setting the osmotically active component of the cell at fixed macromolecular dry mass. To describe this situation, we chose an approach that combines the Donnan equilibrium (*13*, *14*) for the cell volume with a model of biosynthetic fluxes (Fig. 3F). Rollin and coworkers (*13*) recently proposed such a theoretical framework including a biosynthesis module for protein production and a balance equation that relates the accumulation of trapped osmolytes (which, like Rollin et al., we assume to be mostly metabolites and their counter ions) to the number of enzymes that produce or import them (*44*). This model is able to explain the recent observation that mass biosynthesis plateaus while volume keeps increasing in G1 arrested cells (*2*, *13*), which resembles the decoupling that we observed upon CHX treatment. We adapted this model to the appropriate level of description for the purpose of this study (see the SI Text sec. 1 and 2). Using analytical calculations, the model provides a simple equation relating the volume growth rate (Fig. 3F) to the macromolecular dry mass density.

Because the parameters of this model have simple and explicit physical meaning, we then used them to calculate values that can be also estimated from an independent experimental data, such as the dataset on untreated steady growing cells, in order to test the consistency of our modelling approach. First, we obtained an estimate for the fraction of osmotically active volume in a cell (the portion of a cell’s total volume that is capable of participating in osmotic processes), resulting in about 70%, which corresponds to previous estimates (*12*). Specifically (see SI Text sec. 1 and 2 for more details) this quantity can be derived from the offset of so-called Ponder plots, which report the volume changes upon osmotic shocks of different entities (*12*, *16*). Subsequently, using the model to predict the change in volume growth rate upon an arrest of dry mass accumulation, we obtained a quantitative prediction of the magnitude of the increase, corresponding to a an increase of 11_v_ of 0.07 h^-1^, which is compatible in sign with the experimentally observed jump, and within the correct order of magnitude, although it overestimates the observed value (Fig. 3C, see SI Text sec. 2D for the estimation of this value from the model; in short, the model relates the increase to the values of the growth rates and the dry mass density of unperturbed cells, which are obtained from the measurements before the treatment). Besides yielding consistent physical parameters, this model reproduces quantitatively the experimental data for the growth arrest experiment (see fits in Fig. 3A-D). It also provides a simple mechanistic explanation for our surprising observation of an increase in volume growth rate when protein production is stopped: although protein production is stopped, shortly after the treatment, the cell still contains the same quantity of enzymes (and transporters) producing trapped osmolytes. Hence, the rate of accumulation of trapped osmolytes will be maintained until other regulatory processes reduce the enzymatic activity, which our observations suggest to occur over several hours (Fig. 3E). The maintenance of the rate of osmolyte accumulation in a context of arrest of protein production (and thus a major reduction in the consumption of the metabolites, the main trapped osmolytes), leads according to the model to an increase in the rate of accumulation of osmolytes and thus to the immediate increase in the volume growth rate, in line with what we observed experimentally. Our osmo-metabolic model and observations thus support a first ‘hard wired’ mechanism by which volume and dry mass growth rates are coupled. Osmotic equilibrium on seconds time scales paralleled by biosynthetic fluxes on hours timescales explain how these two variables are on average equal, while remaining largely independent on short timescales, volume growth rate being indirectly determined by the quantity of proteins via protein control over the rate of accumulation of trapped osmolytes such as metabolites.

Since the osmo-metabolic model explains well the effect of an arrest in macromolecular dry-mass accumulation, we asked whether the same mechanism is theoretically sufficient to ensure dry mass density homeostasis. The model predicts, for a regime of steady state growth, a positive coupling of the volume growth rate to the dry mass density, under the assumption that the accumulation of trapped osmolytes depend on a metabolic flux that should scale with the number of metabolic enzymes, hence with the total dry mass (Fig. 3F). Cells that are denser would thus grow faster in volume (they have a higher overall flux, hence more capacity to produce trapped osmolytes, see SI text for a detailed mathematical description), leading to homeostatic feedback. This prediction is verified in our dataset for both suspended cell types (Fig. 3G,H). However, this trend is absent for HeLa cells grown on an adherent substrate (Fig. 3H). We hypothesized that this could be due to the effect of adhesion on cell volume, via the previously described mechano-osmotic coupling, producing a high level of fluctuations in volume growth rate at short time scales (*12*, *45*). This effect could prevent the experimental observation of the coupling, even if it existed. Consistently, the coupling could be restored by growing adherent HeLa cells on a non-adhesive substrate (Fig. 3H). In conclusion, the osmo-metabolic coupling described by our model provides a potential mechanism for a homeostatic feedback between volume growth rate and dry mass density. It also explains the behavior of volume and mass growth rate in response to an arrest in protein production.

Previous studies suggested that protein production itself, and thus mass growth rate, could depend on protein concentration, and thus on dry mass density (*3*, *10*, *15*, *24*). We asked whether we could observe this coupling. Confirming these previous hypotheses, we found that, in our dynamic single-cell dataset, mass growth rate decreases with increasing dry mass density, providing a second homeostatic coupling (Fig. 4A). This relationship holds robustly in the three cell types (Fig. 4B and S4 A). In order to assay the timescale at which this mechanism operates, we induced an abrupt decrease in cell dry mass density using a hypo-osmotic shock by adding 30% of water in the cell culture medium. As expected (*12*), within less than a minute, cells had swollen by 25 to 30%, followed by a volume regulatory decrease, on the timescale of several tens of minutes (Fig. 4C). The adaptation was only partial, resulting in a final cell dilution of about 10% (Fig. 4C and S4 B). Cells kept increasing their dry mass steadily during all the phases of volume changes (Fig. 4C). Strikingly, dry mass growth rate showed a rapid increase, approximately doubling in about 10 minutes after the hypotonic shock, and remained high for about one hour before slowly decreasing (Fig. 4D,E and S4 C). This induction of mass growth rate upon osmotic dilution is reminiscent of the over-shoot in mass growth rate observed at mitotic exit (Fig. 1D,F and S1 K-N), when cells restart dry mass accumulation whilst their dry mass density is still low. Together, these results suggest that dry mass growth is coupled to density with a minute timescale adjustment of dry mass growth rate that operates at all time in growing cells.

**Figure 4.**
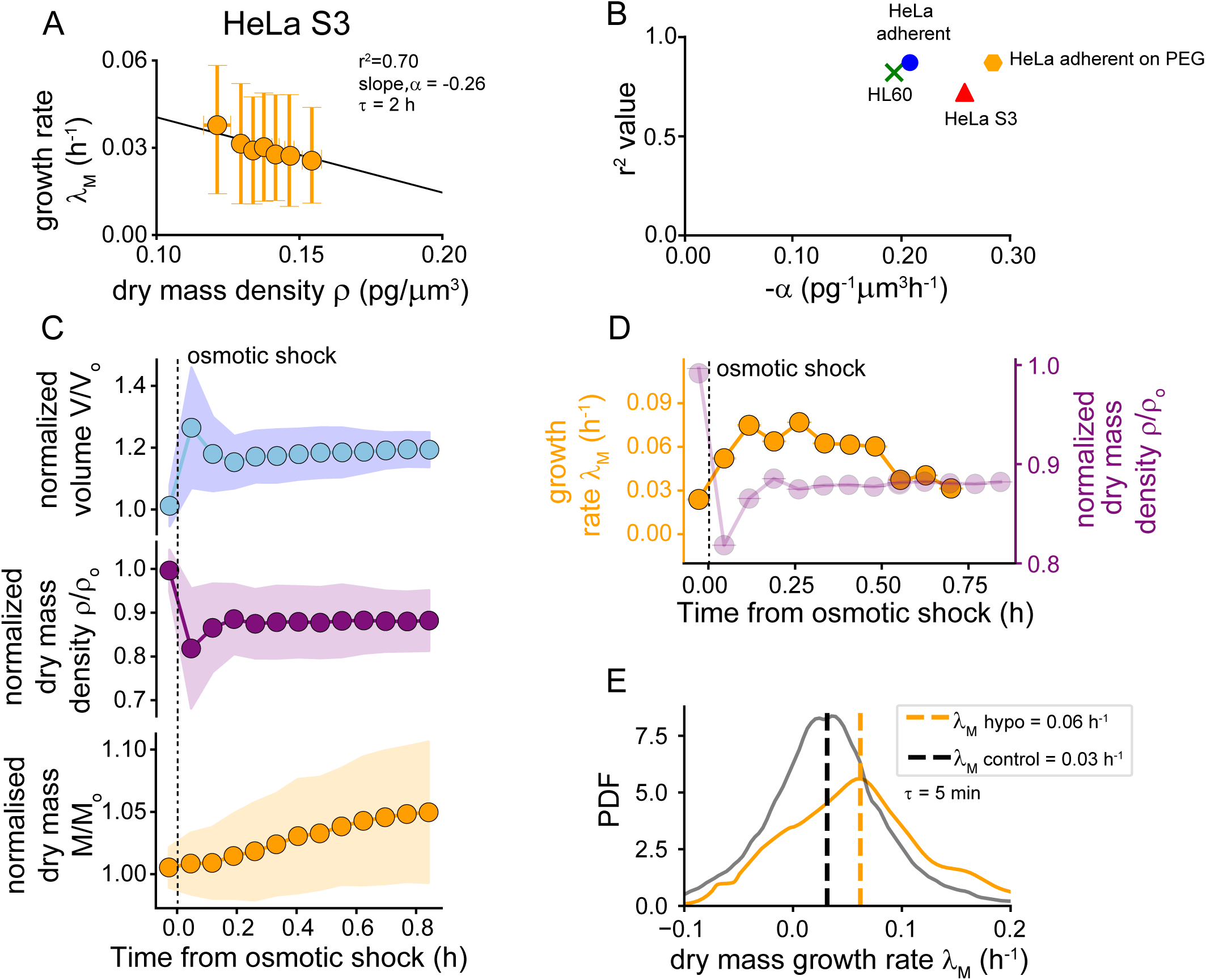
Homeostatic modulation of dry mass growth rate acts at the minute timescale. **(A)** Conditional average of mass growth rate (measured over a τ=2 hours window) at fixed dry mass cell density (linear fit, solid line, p <0.005). **(B)** Goodness of the linear fit (r^2^) against the slope (α) for mass growth rate versus dry mass density plots (see Fig. S4A), for each cell type and conditions **(C)** Average volume, dry mass density and dry mass following a 30% hypoosmotic shock, normalized to their value before the shock (shock is applied at time 0, dotted line). **(D)** Conditional average of mass growth rate and dry mass density plotted against time from the application of the hypoosmotic shock. **(E)** Distribution of single cell mass growth rates (𝜆_𝑀_) following an hypoosmotic shock compared to control steady growing cells. A: HeLa S3 cells (n=140 cells, N=3); B: all cell types and conditions as in Figure 2D; C-E: HeLa adherent cells with osmotic shock (n=52, N=3); Data are represented as binned mean values (solid circles), A, C: ± SD; D: ± SEM, with equal weighted bins.

The observation of a coupling of volume (new) and mass (consistent with previous studies (*3*, *10*)) growth rate to dry mass density opens the question of whether the two couplings can jointly explain the observed density homeostasis. This observation prompted us to propose a second mathematical model with two key phenomenological ingredients, to explain dry mass density homeostasis (Fig. 5A and SI Text, sec. 3 and 4 for details). In this new model, the mass growth rate is not autonomous, but is itself coupled linearly to macromolecular dry mass density on a short timescale (since here we consider timescales larger than the minutes adaptation of mass growth rate observed above). The two combined homeostatic mechanisms in our model lead to the emergence of a specific target density (Fig. 5B, π*), meaning a dry mass density towards which the cells tend to converge, because cells with a higher dry mass density would decrease their density while cells with a lower dry mass density would increase it. This target density corresponds to the dry mass density at which the two growth rates are equal. The time derivative of the dry mass density should thus decrease with increasing density, be positive at low densities and negative at high density, and cross zero at the target density value defined by the crossing of the dry mass and volume growth rate curves. Plotting the time-derivative (on 2-hours steps) of the dry mass density in steady growing single cells, as a function of dry mass density validates this prediction (Fig. 5C,D). This plot shows a strong negative dependency in all cell lines (Fig. 5D-F), except during mitosis, when biosynthesis shuts down and cells swell (Fig. S5 A). This phenomenon was observed robustly for various time intervals ranging from minutes to hours timescales (Fig. S5 B), demonstrating that homeostatic mechanisms are constantly at work, even on short timescales. The target density values and the slope of the time derivative of the dry mass density correspond to the values predicted by the model based on the experimental slopes of the dry mass and volume growth rates as a function of dry mass density, for each cell line, showing the consistency of our model (Fig. 5G, H, see also SI Text sec. 4AB). Surprisingly, the value of the target density was very similar for the different cell lines we assayed (Fig. 5G) and did not depend on the timescale of the derivative (Fig. S5 C). Hence, the existence of a density-sensing mechanism acting on mass growth rate, in addition to the osmo-metabolic mechanism coupling volume growth to density, coherently explains the existence of the target density on a quantitative level. This suggests that these two couplings might be sufficient to recapitulate the robust dry mass density homeostasis observed in our data, i.e., the maintenance of a dry mass density distribution centered around a target density, in a population of growing cells, despite large variations in density following each mitosis.

**Figure 5.**
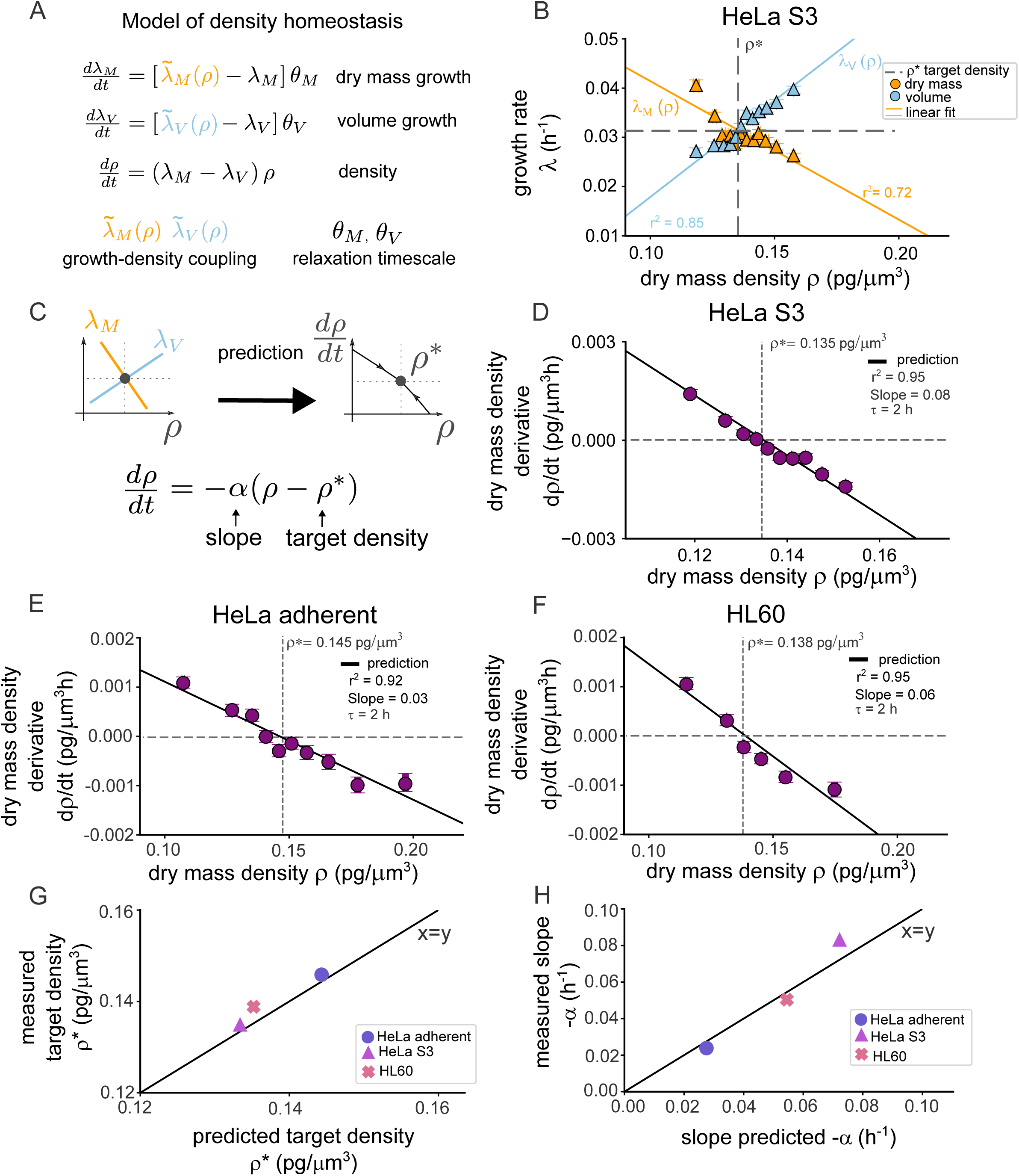
Density-coupled mass- and volume growth rates enforces density correction towards a target density. **(A)** Principle of the mathematical model of density dynamics from dry mass and volume growth (see Supplementary Information file for more details). In the model, dry mass and volume grow exponentially with growth rates 𝜆_𝑀_and 𝜆_𝑉_. The growth rates depend on the dry mass density on average. The functions 𝜆_𝑀_(π) and 𝜆_𝑉_(π), which we define as homeostatic modules, encode the dependence of dry mass and volume growth rates on density (linear fit, p = 6.73 e^-47^ for 𝜆_𝑀_(π), p = 1.66 e^-104^ for 𝜆_𝑉_(π)) with relaxation timescales Σ_M_ and Σ_V_, thus controlling how dry mass density evolves. **(B)** Mean growth rates in dry mass (orange) and volume (blue) plotted against dry mass density, defining the target density π* where the two growth rates are equal. **(C)** Schematics and equation showing how the model can predict the target density π* from the data shown in panel B, a value that can also be directly extracted independently by measuring the time derivative of density as a function of density. **(D-F)** Conditional average of the dry mass density derivative with respect to the dry mass density at the start of the time window used for the derivative for (D) HeLa S3, (E) HeLa adherent and (F) HL60 cells, discrete derivatives measured over 1 = 2 hours. Points are binned mean ± SEM with equal weighted bins. Solid black lines are model predictions for the change of dry mass density derivative with dry mass density of cells and dashed line indicates the target density (π*). p-value (p = 2 e^-4^ for (D) HeLa S3, 1.13 e^-4^ for (E) HeLa adherent, and 2 e^-4^ for (F) HL60). π*, slope (α) and r^2^ are given for each plot. **(G)** Experimentally measured target density for different cell types plotted against model prediction for the target density. **(H)** Slope of the linear fit on the experimental data shown in panels D-F plotted against model prediction. The experimental dataset used is the complete dataset for steady growing cells, used in other figures (e.g. Fig. 2D).

However, growth rates are noisy at the single-cell level, and density is perturbed at each cell division cycle (during mitosis and respreading). To understand how the distribution of dry mass density is maintained through cycles of growth and division, we need to understand the balance between correction mechanisms and sources of fluctuations. We thus expanded our dry mass density homeostasis model by adding two sources of noise (Fig. 6A and SI Text sec. 5): (i) continuous-time noise in the growth rates of single cells, (ii) discrete-time noise gained after each division event due to mitotic swelling and post-mitotic spreading, where we know the variability of dry mass density increases (based on experiments shown in Fig. 2B). The model parameters, such as the level of noise, as well as the strength of homeostasis, can be extracted from the experimental dataset (see SI text sec. 5 and 6 for details). In the model, we assume that without any source of noise, the growth rate of single cells at fixed dry mass density takes the value determined from the experimental data (growth rate versus dry mass density plots shown in Fig. 3 for volume and in Fig. 4 for dry mass). We also assume again that the mean dry mass and volume growth rates are coupled in a homeostatic manner to the dry mass density, with different coupling levels (corresponding to the slopes of the coupling in Fig. 5A). These elements define the target density in the bulk phase (Fig. 5B). The perturbation felt across a division can be estimated by the observed jumps of density from mitotic entry to end of spreading (see SI notes and Fig. S6 A,B). From the experimental data, we can follow lineages for up to two consecutive division cycles. Plotting the CV of dry mass density along these two cycles shows that the CV is high at the beginning of each cycle and lower at the end (colored circles in Fig. 6B) as expected from our previous analyses demonstrating density homeostasis during one division cycle (Fig. 2). However, due to the limited observation time, we cannot observe directly how the CV would evolve on longer timescales experimentally. By contrast, the model allows us to assess the effects of noise and of the homeostatic coupling over many cycles of growth and division (Fig. 6BC, SI notes and Fig. S6 C-H). To extract the predictions for long-term relaxation, we simulated dry mass density tracks over several cell division cycles. The CV of the simulated tracks followed the same trend as the experimental data on the first two cycles but could be extended for more cycles (white triangles in Fig. 6B, Fig. S6F and SI Text, sec. 5 and 6). This allowed us to test the longer-term outcome, for the dry mass density distribution, under various hypothesis for the balance between correction mechanisms and sources of fluctuations. In the absence of noise but in the presence of a homeostatic mechanism, the CV goes to zero, that is, all cells reach the target density within a few division cycles (Fig. 6C, green triangles). In the presence of noise, but without any homeostatic mechanisms, the dry mass density range diverges and the population slowly diffuses away from the target density (Fig. 6C, red triangles). When both the homeostatic mechanisms (with the level of restoring force extracted from the experimental dataset) and the noise (also at the level measured in the experimental dataset) are present, the dry mass density distribution always reaches a steady variance, independent of the number of cycles simulated (Fig. 6C, blue triangles). This shows that the balance between the level of homeostatic coupling and the level of noise present in the cultured cells defines a stable rage of dry mass density across a large number of division cycles, and that, without this coupling, this level of noise would induce a dispersion in the distribution of dry mass density.

**Figure 6.**
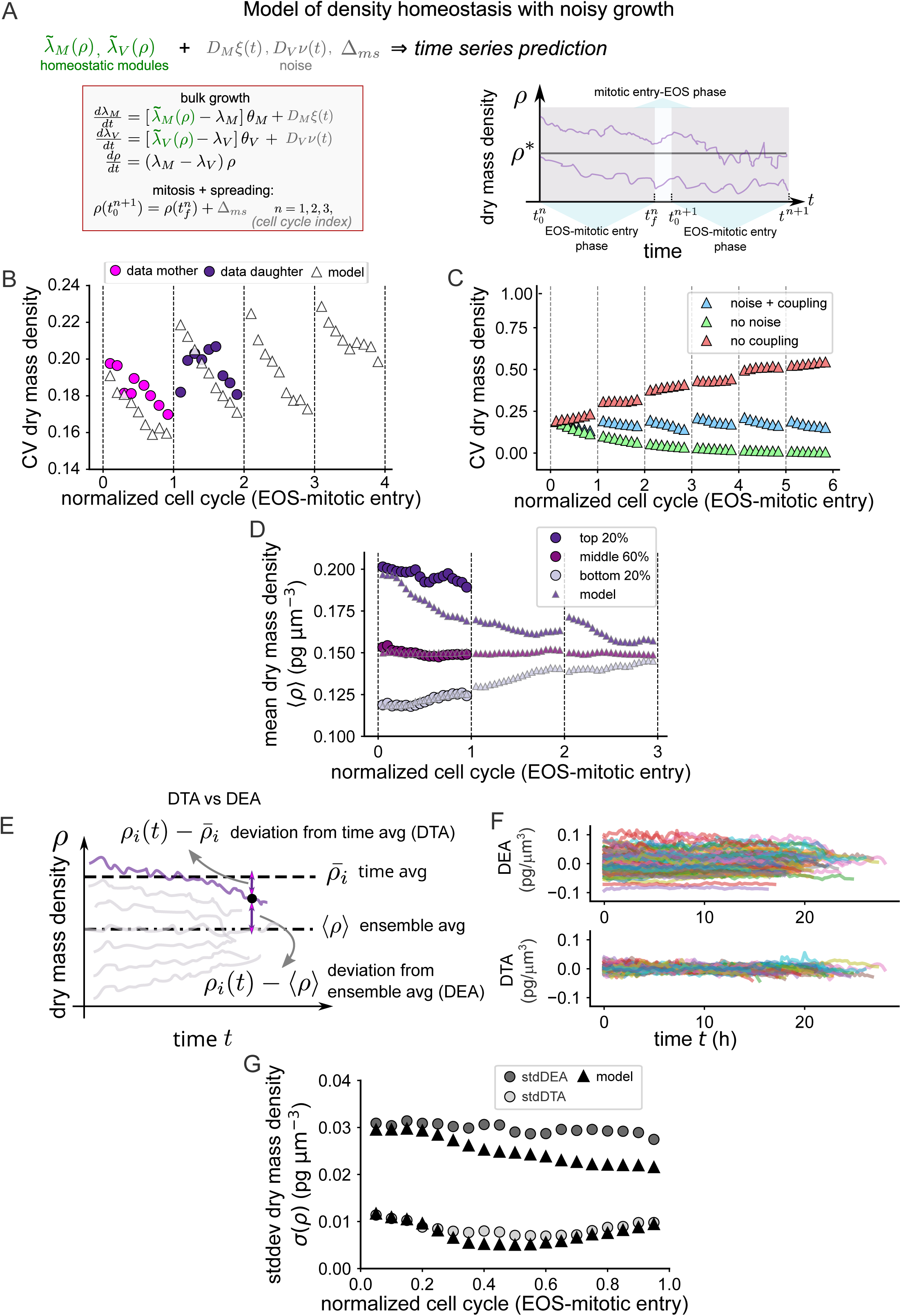
A mathematical model of noisy growth predicts the distribution of dry-mass density in a population of proliferating cells across generations. **(A)** *Left*: Sketch of the mathematical model. The model includes the homeostatic relationship between growth and dry mass density (see Fig. 5A) but with noisy growth that comes both from instantaneous noise during the cell cycle (D_M_*ξ(t) and D_V_*ϖ(t)) and noise due to mitosis and spreading (Δ_ms_). *Right*: The model simulates explicitly dry mass density tracks across the bulk of the cell cycle (between EOS and mitotic entry), including the effect of mitosis and spreading as an increase in the noise as seen in data (see SI file for more details). **(B)** CV (coefficient of variation) of dry mass density across several cell division cycles. Magenta and dark purple points are experimental data on two cell division cycles with mothers and daughters of the same lineage. Triangles show the prediction of the model calibrated with experimental data (see SI). **(C)** CV of dry mass density from simulated single cell trajectories, in the presence of noise and without homeostatic coupling (red triangles), or with coupling (blue triangles), and with coupling by without noise (green triangles). **(D)** Evolution of the average dry mass density in different quantiles of the density distribution, during the cell division cycle. Dark purple points: cells initially in the top 20% of the density distribution at EOS. Grey points: cells initially in the bottom 20% of the density distribution at EOS. Purple points: all the other cells (60% of the cells, around the mean density). Circle and triangle points indicated experimental data and model respectively. Experimental data is represented as binned mean ± SEM. Experimental data are shown only in the first generation, while the model is run for several generations. **(E)** Schematics explaining the calculation of deviation from time average (DTA) and deviation from ensemble average (DEA), on single cell dry mass trajectories between EOS and mitotic entry. **(F)** DEA (top) and DTA (bottom) for single cell dry mass density trajectories for experimental data. **(G)** Temporal evolution of the standard deviation of the single cells DEA (stdDEA) and DTA (std DTA) between EOS and mitotic entry (std is calculated on single cell data binned by normalized cell cycle time). Dark grey circles: std DEA; light grey circles: std DEA, for experimental data. Black triangle: corresponding model predictions. Normalized cell cycle where 0 is at EOS and 1 at mitotic entry, is plotted on x-axis. The experimental dataset used is the complete dataset for steady growing adherent HeLa cells, used in other figures (n=133 cells, N=3 independent experiments).

Due to the density perturbations introduced at each cell division, within a division cycle not all the cells will reach the target density. We thus looked in more detail into what determines the lifetime of the fluctuations, or equivalently what sets the timescale of dry mass density relaxation towards the target density within a cell cycle. Without noise, the model states that the relaxation timescale τ is roughly equal to the inverse of the difference between the mass and volume growth rates (see Eq. S86 in SI Text, sec. 6), which generally depend on the dry mass density ρ. Therefore, the exact value of τ depends on how the growth rates vary with the dry mass density. For instance, if the volume growth rate did not change with the dry mass density, while the dry mass growth could vary 2-fold, the fastest relaxation rate would take a value of 0.03 h^-1^, since the typical mass and volume growth rate are around 0.03 h^-1^. Therefore, the key parameter is the range of growth rates that can be achieved by changing dry mass density, and it is given by the slopes of the growth rate versus density curves. In the case of adherent HeLa cells, the dry mass density relaxes with a timescale of roughly 10-30 hours in the fastest scenario, which is comparable to a cell cycle time and consistent with the relaxation time scales observed in the non-mitotic spreading experiment (see Fig. 2I-M). In addition, since the difference between the dry mass and volume growth rates increases when cells are further away from the target density, they will relax faster than cells that are close to the target density, while, in the presence of noise, the lifetime of the fluctuations can be extended. To investigate the lifetime of the fluctuations, we focused on the behavior of dry mass density time course of single cells and we plotted the average dry mass density behavior of cells grouped by different initial dry mass densities (Fig. 6D). As expected from the model (triangles in Fig. 6D), the plot shows that groups of cells at the extremes of the dry mass density distribution (far from the target density) relax to the average dry mass density (experimental data: dots in Fig. 6D), but fail to fully reach it within the observed time horizon (the next division). We interpret this phenomenon as a consequence of noisy growth and a slow relaxation timescale. Using a model-based extrapolation, we plotted the predicted mean dry mass density if cells would continue to grow for an extended duration without dividing (triangles in Fig. 6D). The plot shows that the inferred relaxation would bring back the two extremes of the dry mass density distribution only after several cycles. Therefore, although a convergence can be observed in the experimental data, for cells far from the target density, it is too slow to allow a perfect correction within a single cell cycle, explaining the width of the distribution of dry mass densities in the population of proliferating cells.

Finally, to bridge the gap between single cells dry mass density tracks and the distribution of densities at the population level, we looked at the difference between the dry mass density of a single cell and the ensemble average 〈ρ〉 (the average dry mass density across all cells in the population). This difference is called the deviation from the ensemble average (DEA) (Fig. 6E). In addition, we also looked at the difference between the dry mass density of a single cell and its own time average ρ_i_ (the average dry mass density across its own full track) (Fig. 6E). This difference is called the deviation from the time average (DTA). We note that the typical behavior of the cell can be different from the average behavior in two ways: (i) the DEA can be large but of the same order as the DTA (DEA ≈ DTA) (ii) the DEA may be small but still significantly larger than the DTA (DEA ≫ DTA). In the first case, fluctuations are large but all cells fluctuate around the population average (the target density). In the second case, even if fluctuations are small, cells do not fluctuate around the population average but around their own time average, different for each cell. This second kind of behavior is called non-self-averaging (a well-known notion in the field of statistical physics (*46–48*), as taking the average of dry mass density timepoints in a single cell track does not correspond to taking the average of dry mass density timepoints across cell tracks. In our dataset, we found that the DEA was far bigger than the DTA (Fig. 6F,G grey dots and Fig. S6I,J), which means that each cell mostly hovered around its own time average. Crucial to this behavior is the fact that in between cell cycles (from mitosis to end of spreading) the dry mass density shows independent jumps, hence bearing some memory of the previous cycle (Fig. S6 A,B). For tracks simulated using our model, despite the built-in homeostatic mechanism, the DEA (dark grey points) and the DTA (light grey points) also exhibits a non-self-averaging behavior (that is, DEA > DTA, Fig. 6G triangles and Fig. S6, J) just like the experimental data. This is again due to the slow relaxation timescale, which implies that within our observation time window (one cell cycle) single cells maintain a strong individuality in terms of dry mass density. The model predicts that this individuality needs a few cell cycles to be lost (Fig. S6 K-M). In conclusion, our model and simulation reproduce both the non-self-averaging behavior of single dry mass density trajectories (Fig. 6F,G) and the population level dry mass density homeostatic behavior around a target density (Fig. 6B,C and S6 N). This demonstrates that, within the range of parameters displayed by the real cells, the homeostatic mechanism we identified is able, on the timescale of a cell division cycle, to sufficiently correct both the initial noise produced by the division process and the dynamical noise due to noisy growth rates, to maintain the cell population within a controlled stable range of dry mass density centered around the target density.

Our results highlight density homeostasis in a population of proliferating cells as a process able to slowly correct periodic and continuous perturbations that drive individual cells away from their target density. According to simulations, the homeostatic modules we identified are essential to prevent the distribution of dry mass density in a population of proliferating cells to slowly deviate from the target density (Fig. 6C red triangles and Fig. S6F-H). In order to test this aspect experimentally, we aimed at perturbing the homeostatic coupling of growth rates. From our analyses, the coupling of volume growth rate to density should be hard to perturb biologically since it is not due to a specific pathway, but it may be perturbed by introducing an independent external noise on the volume. Consistent with this hypothesis, adherent cells, which have a weaker volume growth to dry mass density coupling, display a larger CV of dry mass density than suspended cells (Fig. S7 A). We reasoned that, because of this, disrupting the coupling of dry mass growth rate to dry mass density in adherent cells should disrupt density homeostasis. The mass growth rate homeostatic coupling relies on a rapid response of mass growth rate to changes in dry mass density (see Fig. 4). On such a short timescale (minutes), the dry mass fraction of ribosomal complexes must remain constant, suggesting that the increase in mass growth rate could correspond to an increase in ribosomal activity or a decrease in protein degradation rate. To assess whether mTORC1, a master regulator of ribosomal activity, is involved in the coupling of dry mass growth rate to dry mass density, we measured phosphorylated p70 S6kinase levels, a readout of mTORC1 activity. We found that, when diluting the cells with a hypotonic shock, phosphorylated S6 kinase level increased (Fig. 7A). This suggests that the dry mass growth rate increase observed immediately following the hypotonic shock could be due to an activation of the mTORC pathway. We thus reasoned that preventing this activation of mTORC could prevent the rapid adaptation of the mass growth rate upon dry mass density changes. To test this hypothesis, we used the mTORC inhibitor Rapamycin at a low concentration (100nM), which partially decreased the mass growth rate (see the dose response in Fig. S3 B,C). We found that this partial inhibition of mTORC was sufficient to fully abolish the increase in mass growth rate upon osmotic cell dilution (Fig. 7B and S7 B). We then treated steady-state interphase growing cells with the same low dose of Rapamycin. We found that, for denser cells, the dry mass growth rate was similar both in control and in treated cells, but that treated cells with a lower dry mass density displayed an almost three-fold lower growth rate than control cells, resulting in a flat relation between the dry mass growth rate and the dry mass density (Fig. 7C and S7C). Directly modulating the rate of protein synthesis with a low dose of CHX (see Fig. S3A for the does response) had the same effect (Fig. 7D and S7D). This indicates that the homeostatic module relying on the modulation of dry mass production depends on translation activity. Therefore, we asked about the effect of perturbing this homeostatic module on the recovery from dry mass density perturbation following division and spreading. As predicted from the model, using the experimental slopes for the volume and dry mass homeostatic modules (Fig. 7E), the time derivative of dry mass density did not depend anymore on dry mass density (Fig. 7F), suggesting that the convergence towards a target density was lost in these cells. Consistently, dry mass density homeostasis across the cell division cycle was lost (Fig. 7G) and the CV of dry mass density was not restored during interphase (Fig. 7H), corresponding to the simulations for a population of cells growing in the presence of noise but without homeostatic coupling (red triangles in Fig. 6C). In conclusion, these perturbation experiments demonstrate that the dual coupling between dry mass and volume growth is not only sufficient in theory, but also required for the correction of dry mass density fluctuations and long-term maintenance of dry mass density homeostasis in a population of proliferating mammalian cells.

**Figure 7.**
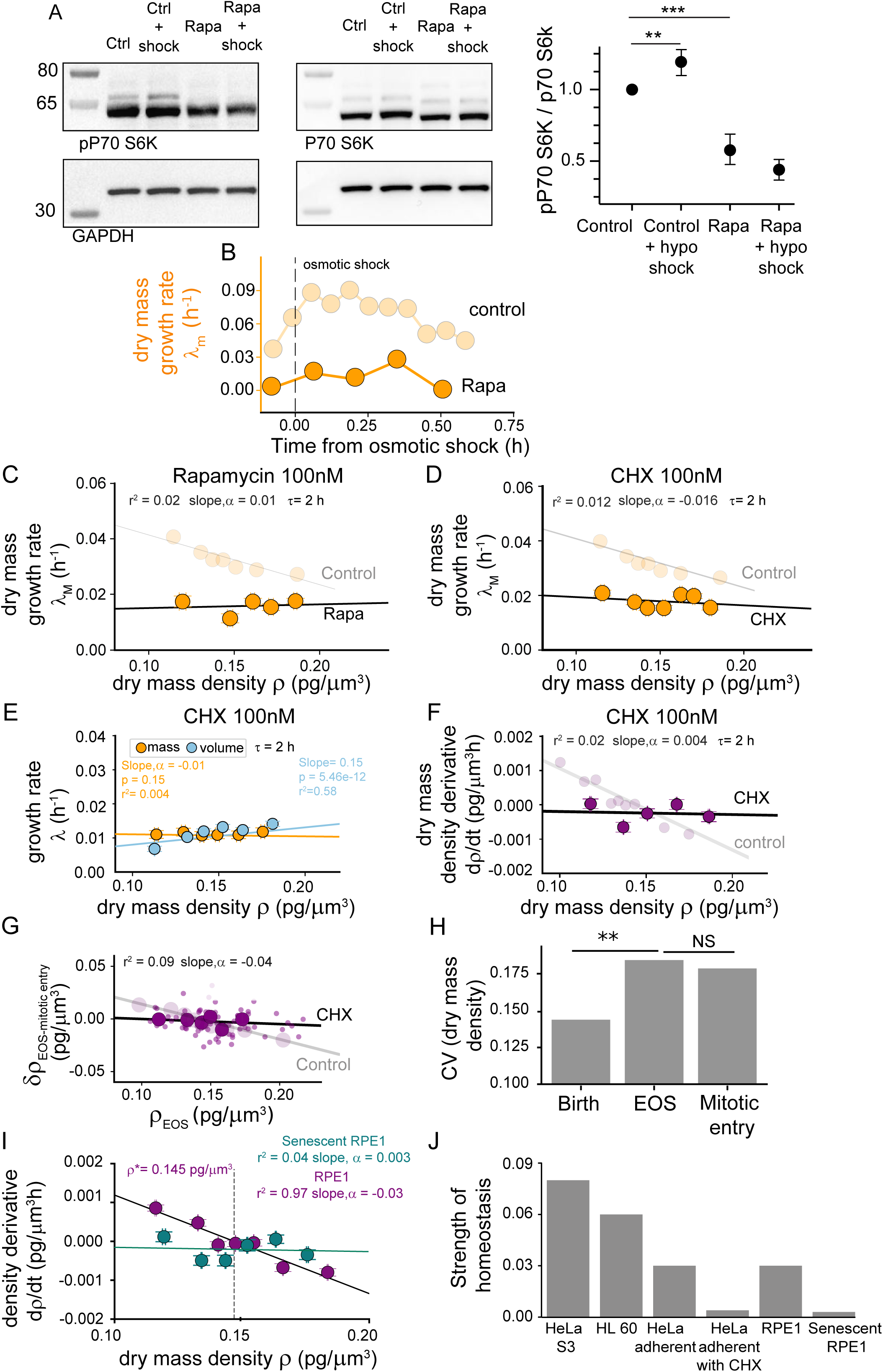
Dry mass density homeostasis is lost upon perturbations of the mTORC pathway, reduced protein translation, and induction of senescence. **(A)** Left: Western blot for phosphorylated p70 S6kinase (pP70 S6K, top left image) along with total P70 S6kinase (P70 S6K, top right image) and GAPDH loading controls (bottom images), in control HeLa adherent cells (Ctrl), upon 30% hypoosmotic shock (Ctrl+shock), Rapamycin treatment (Rapa), and combined treatments (Rapa+shock). (Right) Quantification of pP70 S6K levels: normalized ratio of pP70 S6K over p70 S6K (N=3 independent experiments; ** p<0.05, *** p<0.005). **(B)** Conditional average of dry mass growth rate for 100nM Rapamycin treated cells (dark orange) and control cells (same dataset as in Figure 2D), after a 30% hypoosmotic shock (applied at time 0). **(C-D)** Conditional average of dry mass growth rate against dry mass density for cells growing in the presence of: (C) Rapamycin 100 nM and (D) CHX 100nM. Data for control cells is shown in the background for comparison (light orange, same data as Fig. S4A). Solid line: linear fit (slope and r^2^ values are shown on the plots). **(E)** Conditional average of growth rates in dry mass (orange) and volume (blue) against dry mass density for cells dividing in presence of 100nM CHX. Solid line: linear fit (p-value and r^2^ shown on the plot). **(F)** Target density plot: conditional average of the dry mass density derivative against the dry mass density for cells dividing in presence of 100nM CHX. Solid black line: model prediction. Control cells shown in the background (light purple, same data as in Fig. 5E). **(G)** Dry mass density correction between EOS and mitotic entry against dry mass density at EOS for cells dividing in presence of 100nM CHX. Small dots are single cells and large circles indicate binned mean ± SEM. Solid line: linear fit on the binned averages (slope and goodness of fit, on the plot, p <0.005). **(H)** CV of dry mass density at birth, EOS and mitosis for cells dividing in the presence of 100nM CHX (** p<0.05). **(I)** Target density plot: conditional average of the dry mass density derivative against dry mass density, for RPE1 control cells (purple), and senescent RPE1 cells (green). Solid line: linear fit (slope and r^2^ values shown on the plot). **(J)** Strength of homeostasis measured as the slope of target density plots is plotted for all the cell lines and conditions shown in the article. B-G, I: binned mean ± SEM with equal weighted bins; B: HeLa adherent cells hypotonic shock treated with 100nM Rapa (n=49, N=3); C: HeLa adherent cells growing in the presence of 100nM Rapa (n=120, N=3); D-H: HeLa adherent cells growing in presence of 100nM CHX (n=127, N=3); I: RPE1 control cells (n=68, N=2) and RPE1 senescent cells (n=70, N=2); B-G, I: time derivatives are taken over a time window of 2 hours (1).

These results suggests that cells with an altered metabolism and protein translation could rapidly loose dry mass density homeostasis and show an increasing dispersion as well as a change in the mean of their dry mass density distribution. This is reminiscent of the state of senescent cells, which are known to have reduced metabolic activity. Because this state was recently associated with a reduction in dry mass density (2), we asked whether senescent cells could also show signs of a loss of dry mass density homeostasis. To address this question, we chose to use RPE1 cells, which are immortalized but not of cancer origin, contrary to the other cell types used in the rest of the study, and can be induced to senesce using well established protocols (here we used Palbociclib treatment for 9 days, see methods). We found that senescent RPE1 cells have a higher dry mass and a higher volume, but, as expected from prior work (*2*, *24*), a much lower dry mass density, mostly because the volume of these cells was drastically increased (by a factor of more than 4, while the dry mass less than doubled, Fig. S8 A-C). Plotting dry mass and volume growth rates against dry mass density showed that RPE1 cells behave very similarly to adherent HeLa cells, with a strong negative dependency of dry mass growth rate on dry mass density, but almost no dependency of volume growth rate, likely due to the elevated mechano-osmotic driven noise in volume growth rates in adherent cells (Fig. S8 D, E). In this Pablociclib-induced senescent state, RPE1 cells keep growing robustly in both dry mass and volume, although at a lower rate than control cells. But, similar to HeLa cells treated with low doses of CHX or Rapamycin, senescent RPE1 cells show almost no dependency of growth rates on dry mass density (Fig. S8 F,G). Consistently, while control RPE1 cells have a clear target density, with a value similar to the other cell lines in this study, senescent cells showed no sign of dry mass density correction towards a target density (Fig. 7I,J). In conclusion, these experiments show that non-cancer RPE1 cells behave similarly to the three cancer cell lines, with a very similar target density, while senescent RPE1 cells behave like cells treated with drugs affecting metabolic pathways and show a loss of dry mass density homeostasis, explaining how they drift away from their initial target cell density, to reach a more diluted state. This suggests that dry-mass density homeostasis, although robust in all the cell lines studied, can be lost in the context of specific cellular states, such as senescence, associated with changes in metabolic regulation. The mechanisms that maintain dry mass density homeostasis are thus crucial in ensuring a proper physiological state and disrupting these mechanisms may lead to cell states associated with ageing or diseased conditions.

## Discussion

In this study, we employed mathematical modeling with explicit and interpretable parameters, and provided a comprehensive dataset of single-cell measurements of dry mass and volume throughout the cell division cycle. Our findings reveal that growing mammalian cells possess a mechanism for macromolecular dry mass density homeostasis, enabling them to maintain a stable global concentration of proteins within a targeted range. This capacity stems from two independent processes that adjust the growth rates of mass and volume as a function of dry mass density.

Our results lead us to conclude that the first process is biophysically determined by osmotic equilibrium, primarily relying on small osmolytes of metabolic origin, likely including amino acids, as proposed by Rollin and colleagues (*13*). The excess volume growth following a halt in mass growth supports this model, which – once we cast it in mathematical form - predicts the observed increase in volume growth rate, as well as the observed proportionality between volume growth rate and dry-mass density.

The second process regulates protein biosynthesis (and thus macromolecular dry mass) through concentration-dependent modulation of the mTORC pathway. Together, these mechanisms create a robust alignment of mass and volume growth rates, defining a target density that equips growing cells with the ability to manage dry mass density fluctuations throughout the cell cycle. Finally, our further phenomenological model of density homeostasis connects volume and mass correction to overall density control, showing that the first two processes quantitatively explain the latter.

We identified two primary sources of dry mass density perturbations that act “locally” in the cell cycle, before and after cell division, although additional factors may exist in living tissues. The first source is inherent to the cell division process, during which volume increases sharply (a phenomenon known as mitotic swelling observed in all mammalian cells,(*36*, *42*) while dry mass growth halts, likely under the regulation of mitotic kinases (*41*). Upon mitotic exit, as these kinases deactivate, protein production resumes rapidly in still-diluted cells, explaining the initial spike in mass growth.

The second source of dry mass density perturbation arises from mechano-osmotic coupling in spreading cells, a phenomenon we and others previously described (*11*, *12*), due to the modulation of ion fluxes by cell surface tension. This coupling contributes to post-mitotic dry mass density dispersion and generates fluctuations throughout the cell cycle, potentially accounting for the greater coefficient of variation in dry mass density observed in adherent cells compared to suspended cells. We hypothesize that, in densely packed tissues and during cell migration (*12*), mechano-osmotic coupling could induce dry mass density fluctuations, making the regulation of dry mass growth rate essential for maintaining dry mass density homeostasis in multicellular organisms.

Together, these two perturbations around cell division counteract a “bulk” phase of the cell cycle that attempts to restore a target macromolecular dry mass density by the two homeostatic mechanisms we identified (osmo-metabolic response and dry mass density coupled regulation of biosynthesis). The resulting equilibrium entails considerable cell-to-cell diversity in dry mass density as well as stochastic oscillations (where, for example, the CV of density is a periodic function with the period of the cell division cycle). A similar phenomenon is being proposed by a parallel study for growth-rate fluctuations in bacteria (*49*). Our further stochastic homeostasis mathematical model captures this phenomenology and shows how the homeostatic control is strictly necessary, given the perturbations, to maintain a steady (albeit wide and showing considerable lineage individuality) density distribution across cells.

While our work establishes the phenomena of dry mass density control and identifies a target density, it does not elucidate the detailed molecular mechanisms involved. Future studies should focus on three unresolved aspects: the mechanism underlying dry mass growth rate modulation, the nature of the small metabolites driving volume growth, and the definition and significance of the target density and the homeostatic range of dry mass density around it. We discuss these aspects below.

Recent advances suggest that various processes could influence dry mass growth rate modulation. Physical phenomena affecting compound diffusion may induce dry mass density sensitive changes in protein production. For instance, crowding effects alter the diffusion of molecules based on their size (*10*), while viscosity effects are less specific (*3*). These phenomena may impact protein translation and degradation rates, linking net dry mass production to dry mass density (*3*). These physical effects act directly on ribosomal activity through diffusion-limited processes, while our findings indicate that the mTORC pathway is involved, suggesting a more specific mechanism for dry mass density sensing. A parallel population-level study focused on recovery from osmotic shocks, also finds modulation of dry mass and protein synthesis rates as a function of dry mass density (termed “cell mass density”), and explore several molecular mechanisms, including a role for the Na^+^/H^+^ transporter (NHE) and the mTORC pathway (*11*). The same study also proposes a role for nuclear import/export, although, similarly to transcriptional feedback, this could only play a role on longer timescales than the mechanisms we identified here in single cells. The slow global protein degradation rates reported in the literature (in the hours timescale, (*50–52*)) make it unlikely that they account for the rapid increases in dry mass growth rate observed upon cell dilution; this hypothesis was experimentally ruled out in the case of long-term adaptation of dry mass density following osmotic shocks (*11*).

Physico-chemical phenomena, such as the formation of protein condensates, may serve as sensors of water potential by modulating enzymatic activities, especially in contexts of volume regulation (*27*). Condensates formed by WNK1 or other mTORC regulators could thus act as dry mass density sensing mechanisms. Additionally, integrated stress response pathways (*28*, *53*) might respond to changes in global protein concentrations and induce translation upon dilution. Determining whether the homeostatic modulation of dry mass growth rate results from an active density-sensing pathway or from dry mass density-dependent physical phenomena will require thorough investigation into its physical and molecular foundations.

An intriguing question arising from our work is the identity of the osmotically dominant cellular compounds. While our model describes intracellular ions in a simplified way, as slaved to the electroneutrality condition and mainly acting as counterions (*12*), we acknowledge that ions are very abundant and may suffice to tilt cell volume in certain conditions (*54*, *55*). However, we believe that the effects observed in this study, where no large perturbations of ion pumps and ion equilibria are expected, reasonably come from small neutral osmolytes of metabolic origin, as evidenced by the volume rate increase observed upon arrest of protein synthesis. Is there a single metabolite primarily responsible for cellular growth, or does volume growth depend on a multitude of species? In unicellular organisms, metabolite concentration analyses have been extensively conducted. For instance, in *E. coli*, free glutamate and free potassium are present in nearly equal amounts, ensuring both osmotic equilibrium and electroneutrality (*56*). In yeast, glutamate significantly dominates other amino acids, reaching tens of millimolar concentrations (*56*). In mammalian cells, the exact quantity of amino acids and small metabolites is less well-defined. Recent studies on cancer cell cultures indicate glutamate concentrations comparable to those in yeast and bacteria, approaching 100 mM (*57*, *58*), significantly outpacing other metabolites. The specific abundance of glutamate could make it, at least in cancer cells, a major driver of cell growth in volume, increasing the ‘homeostatic pressure’ of cells that accumulate it at a higher rate (*59*). This would contribute to its central role in defining ‘winner’ cells, providing competitive advantage to cancer cells and making its synthesis pathway a successful target for cancer therapy (*60–62*). Conversely, normal tissue analyses report lower values, where glutamate still predominates but to a lesser degree (*63*), leaving the question unresolved.

A notable finding of our study is the identification of a target density that characterizes the mean dry mass density distribution in proliferating cells. Our model suggests that this target value is influenced by various parameters, allowing for control by external signals and adaptation in different cell types or biological contexts such as cell differentiation and aging (*30*). The existence of a target density aligns with earlier proposals that cells may operate optimally at certain dry mass density values (*64*) and with measurements of narrow buoyant mass ranges across diverse cells and organisms (*65*, *66*), as well as recent report using the same definition we use here, i.e. a ratio of dry mass measured by quantitative phase over volume measured by FxM. This report also finds a surprisingly conserved value for dry mass density across cultured cell lines, with values between 0.15 and 0.2 mg/ml, slightly higher than ours (close to 0.14 mg/ml for the four cell lines studied), potentially due to a different calibration of the quantitative phase imaging (*11*). The value of 0.14 mg/ml can also be derived from protein mass fraction measurements yielding similar values in E. coli (*67*, *68*) (; see SI Text sec. 2 for estimates based on our osmo-metabolic model).

It is particularly striking that calculations of dry mass density from water content data across numerous studies employing various methods on different cell types yield excluded volume measurements between 0.29 and 0.31 mg/ml, consistent with our findings (*69–72*); see also SI Text sec. 2). This suggests a relatively narrow range of optimal operating densities, with the convergence mechanisms we identified potentially representing a phenomenon by which cells maintain optimal densities, deviating only under specific circumstances (*1*, *28*, *30*, *31*, *73–75*).

As shown by a growing number of reports, dry mass density appears to be much more tightly regulated than other parameters like cell volume or dry mass alone (*1*, *28*, *30*, *66*). Our study uncovers how single mammalian cells maintain their intracellular concentrations within a constant range as they grow and divide in culture conditions. If cells optimally function only in a narrow range of protein concentrations, then global deviations in protein concentrations may carry important functional and physiological consequences. Although a change in concentration of a specific protein might not significantly affect cellular physiology, a shift from the global target density due to failing homeostatic pathways would affect proper biosynthesis, as is seen in large senescent cells, placing cells in a state from which they cannot recover. Equally, the robust diluted state and activation of pathways resembling responses to hypotonic shocks observed in aneuploid yeast and mammalian cells (*75*, *76*) could correspond to a disruption of dry mass density homeostasis in these cells. This view would support simple unifying views of many physiologically aberrant states, also related to changes in metabolic or other pathways regulating protein synthesis. The question of whether these changes correspond to protective mechanisms or rather reveal fragilities in cellular homeostatic mechanisms (suggesting they might be involved in a variety of diseased states), opens exciting perspectives for future studies.

## Materials and methods

### Cell culture

Human cervical adenocarcinoma cells HeLa EMBL (kyoto) stably expressing hgem-mcherry (HeLa adherent) was established using the lentiviral vector mcherry-hGeminin(1/60)/pCSII-EF. HeLa S3 and HL60 was bought from ATCC. All these cell lines were maintained in DMEM Glutamax media (Gibco) supplemented with 10% FBS (Biowest and panbiotech) and 1% penicillin-streptomycin (Thermofischer scientific) at 37°C and 5% CO_2_. Retinal pigment epithelial (RPE-1) cells were cultured in DMEM/F12 (Gibco) supplemented with 10% FBS (Biowest/Panbiotech) and 1% penicillin-streptomycin (Thermofischer scientific) at 37°C and 5% CO_2_.

### Drug Treatments

The following pharmacological inhibitors and chemical compounds were used: Cycloheximide (CHX) (Sigma Aldrich) at concentrations 100nM, 1mM and 2mM, Rapamycin (Tocris) at concentrations 100nM, 250nM and 1mM, Torin (Tocris) at concentrations 100nM, 500nM and 1mM and Palbociclib (Sigma) at 1mM. RPE-1 cells were treated with 1mM Palbociclib for 9 days to arrest cells at G1 stage leading to induction of senescence as has been shown earlier.

### Volume measurement with fluorescence exclusion microscopy (FXm)

The volume measurement using FXm has been described earlier (*37*). Briefly, PDMS chambers were prepared a day before the experiments. PDMS (1:10) was poured in the molds to replicate the chambers. To prevent the dextran from leaking into the media, two rectangular PDMS slabs were bonded to the inlet and outlet of the chambers by plasma treatment. The inlet and outlet was created by punching 2 mm holes by a biopsy puncher. The chambers were then cleaned with a scotch tape and then irreversibly bonded to a 35mm glass bottom dish (fluorodish, World precision instruments WPI). These chambers were cured for 4 hours at 70°C in an oven. Afterwards, the chambers were put in vacuum for 2 minutes and then treated with 50mgmL^-1^ fibronectin (Sigma) in PBS for 1 hours for adherent cells. Afterwards, the chamber was washed and incubated overnight with cell culture media. In case of suspended cells, no fibronectin was used and chamber was incubated with cell culture media directly. Cells were then detached with Versene (Gibco) and resuspended in fresh media and then injected into the inlet of the chamber. For adherent cells, 4 hours after cell injection, media was replaced with an equilibrated media containing 1mg/mL FITC dextran in the inlet channel and side reservoirs. For suspended cells, FITC dextran was directly added to the cell suspension before injection into the inlet channel of the PDMS chamber. Images were acquired every 15 minutes for 40 hours for cell cycle experiments, every 30 seconds for 5 hours for spreading experiments and every 2 seconds for an hour for osmotic experiments. Additionally, for osmotic experiments, cells were injected into the chambers coated with 0.01% PLL (Sigma) to maintain a round cell shape during the experiment. Cells were incubated in the chamber for 2 hours before the cell culture media was exchanged with a media of known osmolarity along with FITC dextran, as mentioned earlier. Few images of the cells were taken before the media was replaced to understand the effect of osmolarity change on individual cell trajectories during the experiments. The exchange of media was usually rapid (<1sec) within the chamber. Hypoosmotic solutions were made by water addition to the culture medium. Osmolarity of the working solutions was measured by osmometer Type 15M (Loser Messtechnik).

### Mass measurement with quantitative phase microscopy (QPM)

Cell dry mass measurement was performed using Phasics camera as described in (*78*). Kohler illumination was performed to set up imaging. Care was taken to avoid any condensation in the dishes or around PDMS chips. The reference images were generated by trypsinizing cells in the chamber and removing them. Thereafter, 32 empty fields were acquired on these PDMS chips and a median image was constructed for further image analysis.

### Live-cell imaging

Live-cell imaging was performed on a Ti-Eclipse (Nikon) inverted microscope at 37°C and 5% CO_2_ using a 20x objective (NA 0.45). Images were acquired using MetaMorph (Molecular devices). The excitation source was a LED for FXm experiments to ensure homogeneity of the field of illumination. For combined cell volume and dry mass measurement, a script was used in Metamorph to switch between a CMOS camera (Hamamatsu) and Phase camera (Phasics).

### Image analysis

For cell volume measurement, image analysis was performed as explained in (*38*). Briefly, a MATLAB script, which was written in collaboration with a company QuantaCell, was used for the analysis of the images. The fluorescent signal was calibrated for every time point and individual stage positions, using the intensity of the pillar and around the cell of interest. Once the pillars were identified, background normalization was done using a gridfit method. Subsequently, fluorescence intensity was measured in a polygon around the cell to obtain the cell volume.

For cell dry mass measurement, image analysis was performed using a MATLAB script which was written in collaboration with a company QuantaCell. Each image from the phasics camera was associated with a corresponding reference image to calculate the change in the refractive index and was processed through phasics SDK to generate an intensity and a phase image. The phase image was further analyzed to calculate dry mass of individual cells by performing background normalization, segmenting the cells and finally, calculating the change in the intensity around each cell to calculate its dry mass.

### Choice of key timepoints for cell volume and dry mass analysis

For the analysis of relationship between dry mass density at birth and mitotic entry/division and total cell cycle duration, cells were tracked through the cell cycle and key events were recorded. The birth was defined as the first frame (15minutes) after the cell division (Fig 1 and S1). Mitosis was defined as the point occurring three frames (45 minutes) before the cell division.

### Experiments with translation and mTOR inhibitors

For CHX, Rapamycin and Torin experiments, cells were treated with the drugs at mentioned concentrations and immediately injected in the chambers. For short term experiments, cells were imaged for about 10 hours after treatment with drugs. For experiments with CHX where cell cycle trajectories were recorded, cells that divided in the chambers in first 5-6 hours of the beginning of experiments were tracked for the next 20 hours. For Western Blotting, cells were treated with specific drugs or osmotic shocks, and were scraped from the dish and resuspended in Laemmli buffer. Proteins were separated using sodium dodecyl sulphate polyacrylamide gel electrophoresis (SDS-PAGE) and transferred to PVDF membranes. After incubation with primary antibodies (p70 S6 Kinase antibody, Cell Signalling #9202 and phospho-P70 S6 Kinase antibody, Cell Signalling #9234) and secondary antibodies (IRDye and VRDye (LI-COR)] antibodies, the membrane was visualized using Odyssey CLx Imaging system (LI-COR).

### Data filtering and analysis

All the data analysis was done using custom built Python scripts. For cell cycle experiments, cell trajectories that didn’t have at least one mitosis and birth or that didn’t grow after few hours of experiments was discarded. For all the plots, solid circles are binned mean values calculated on equally weighted bins weighted on the number of observations in each bin. Growth rates were calculated by performing linear fit over 4 data points. For the plots where a linear relationship was tested, linear regression on the median bins (details in the analysis code) weighted by the number of observed cell counts in each bin was performed. The result of the fit with its slope coefficient and coefficient of determination (r^2^) is always indicated on the plots. Unless stated otherwise, statistical significance was determined by two-tailed unpaired or paired Student’s t test after confirming that the data met the appropriate assumptions of normality, independent sampling and homogenous variation). Statistical data is presented either as mean ± SEM or SD. Sample size (n) and number of independent experiments (N) is mentioned in the figure legends of each figure panel corresponding to the data. Individual cells were collected across all the fields (n>25) in a single FXm chip.

## Supporting information

Supplementary Model file

---

**Supplementary Figure 1 (related to Figure 1).**
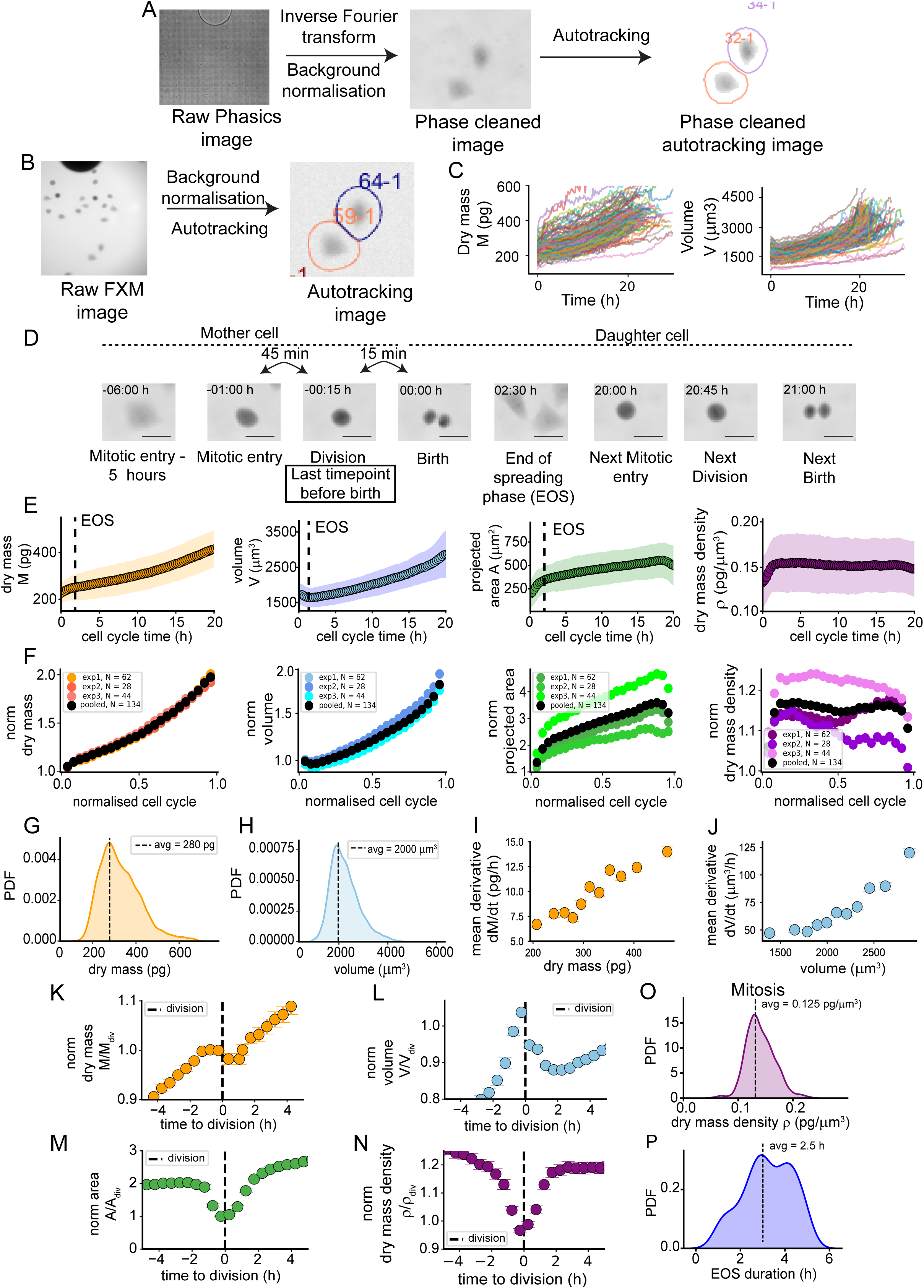
Dry mass and volume growth of single HeLa adherent cells. **(A)** Representative images and analysis pipeline for QPM (see Methods for details). **(B)** Representative FXm images and analysis pipeline for FXm (see Methods for details) **(C)** All single cells trajectories analysed in this dataset in dry mass (left) and volume (right) as a function of time from birth (time 0) **(D)** Representative images used to determine the phases of the cell division cycle. The reference time, called birth, is the first image showing a clear furrowing. Division is the timepoint preceding it (15 minutes before) and mitotic entry is the timepoint one hour before. EOS is determined from the measure of the spreading area, at the end of the fast spreading phase, for each single cell (see methods for details) **(E)** Dry mass (orange), volume (blue), projected area (green) and dry mass density (purple), against time after birth (t=0 h is birth) **(F)** Normalised (relative to value at time 0) values for the three independent experiments (exp1,2 and 3) and pooled values (indicated by black circles) of, left to right: dry mass, volume, projected area and dry mass density, plotted against normalized cell cycle time (t/t_div_) with t/t_div_ = 0 for birth and t/t_div_ =1 for division. PDF of **(G)** dry mass and **(H)** volume of all the cells measured during the whole cell cycle from birth to division. Test for exponential growth of cells in their **(I)** dry mass and **(J)** volume by plotting time derivatives against mean. Mass and volume derivatives were calculated over 1 hour. Binned averages normalized (relative to value at division) for **(K)** dry mass, **(L)** volume, **(M)** projected area and **(N)** dry mass density, plotted against time from division (−5 hours to +5 hours with t=0 at division). **(O)** PDF of the dry mass density of all the cells across mitosis (mitotic entry to division). **(P)** PDF of the duration of the end of spreading (EOS) phase for all the single cells. E,F,I,J,K-N: binned averages ± SEM calculated on equally weighted bins weighted on the number of observations in each bin. All the data are from HeLa adherent cells (n=133 cells, N=3 independent experiments).

**Supplementary Figure 2 (related to Figure 2).**
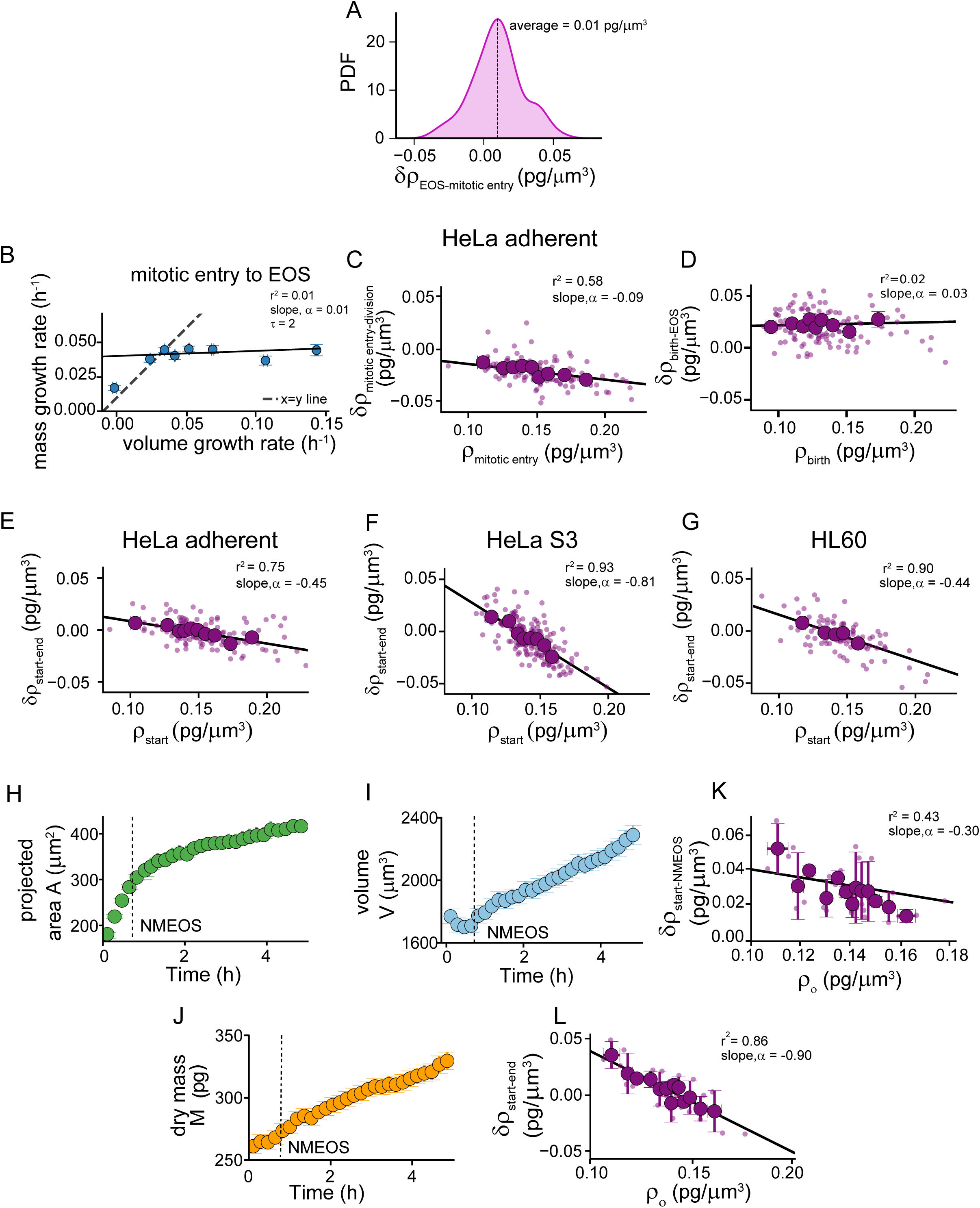
Dry mass density during the cell division cycle and during spreading experiments, in various cell lines. **(A)** Distribution of the difference between the dry mass densities of single cells at EOS compared to the value at mitotic entry for their mother cell **(B)** Mass versus volume growth rates for individual cells over 2 hours windows from mitotic entry to division. (**C-G)** Dry mass density correction (8π), **C:** in mitosis (between mitotic entry and division) against dry mass density at mitotic entry (linear regression, p = 0.06); **D:** during post-mitotic spreading (between birth and EOS) against dry mass density at birth (linear regression, p = 0.69); **E-G:** for a period of 6 hours during interphase (between densities at end and start times) against density at start for E: HeLa adherent; F: HeLa S3; G: HL 60 cells; **(H)** Mean spreading area, **(I)** volume and **(J)** dry mass during a spreading experiment. Dashed line indicates NMEOS. A-E: cycling Hela adherent cells (n=133, N=3); F: HeLa S3 (n=115, N=3); G: HL 60 (n=115, N=2). H-I: spreading HeLa adherent cells (, N=3 independent experiments; for all the plots in this figure, solid circles are binned mean values calculated on equally weighted bins weighted on the number of observations in each bin; C-G: small dots are single cell data and large ones are binned average ± SEM; solid line is a linear fit on the binned averages (slope and goodness of fit, r^2^ are shown).

**Supplementary Figure 3 (related to Figure 3).**
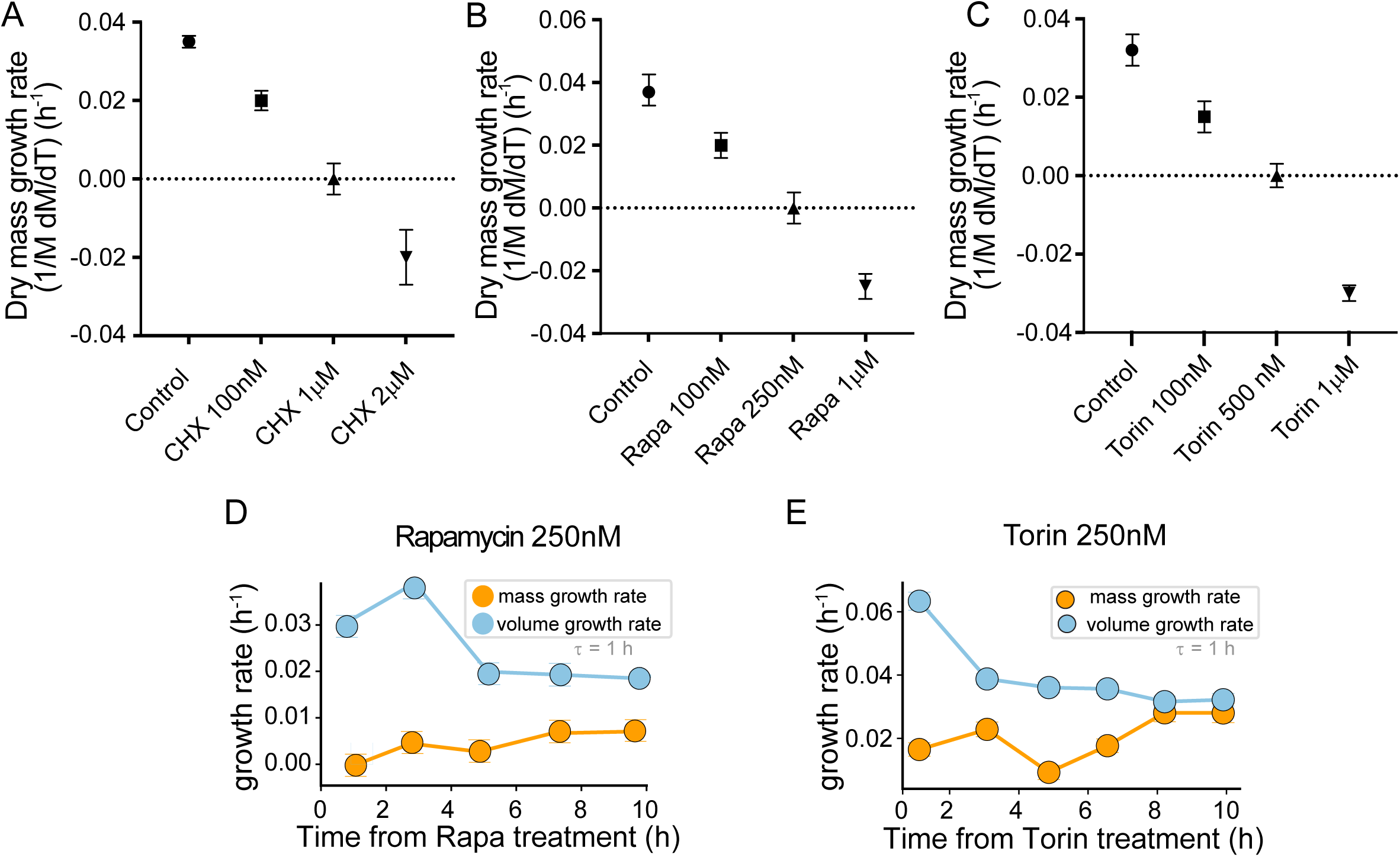
Dose responses for dry mass growth drugs and volume response to dry mass growth arrest. Dose response effect on dry mass growth rate over a period of 4 hours of treatments with **(A)** Cycloheximide (CHX) **(B)** Rapamycin **(C)** Torin (n>50 cells per replicate, N=3 replicates per concentration). Average mass and volume growth rates (measured for τ = 1 hour) plotted against time for HeLa adherent cells treated with **(D)** Rapamycin 250nM (n=72, N=2) and **(E)** Torin 250nM (n=85, N=2). Data is represented as binned mean values (blue circle) ± SEM.

**Supplementary Figure 4 (Related to Figure 4).**
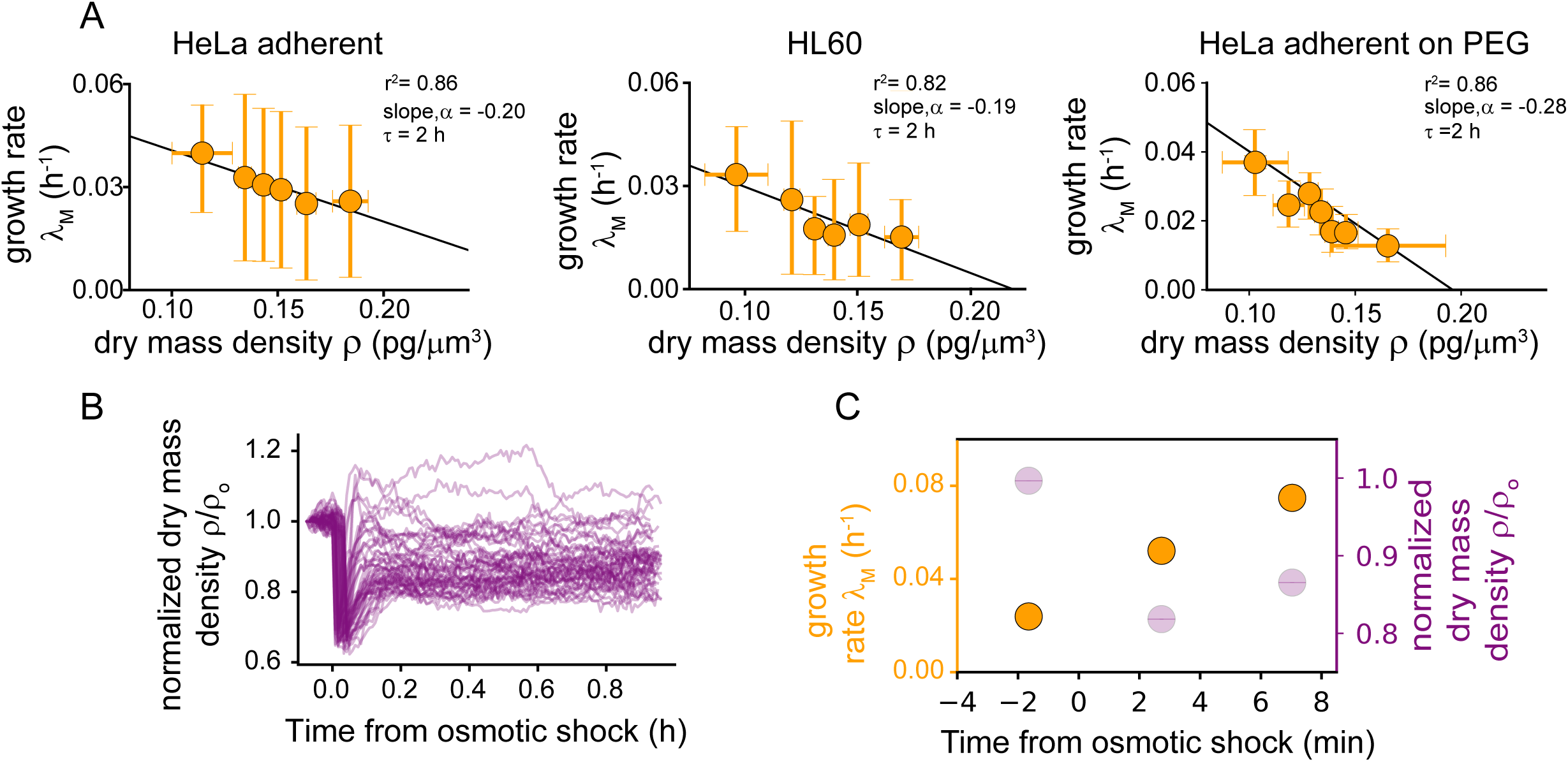
Homeostatic modulation of dry mass growth rate across cell lines. **(A)** Conditional average of mass growth rate (measured over a τ=2 hours window) at fixed dry mass cell density (linear fit, solid lines, p <0.005 for all plots), for, left to right: HeLa adherent, HL 60 and HeLa adherent cells platted on a PEG coated substrate. **(B)** Individual dry mass density trajectories, for all the cells, normalized by the value before the shock, after a hypoosmotic shock of 30% applied at time 0. **(C)** Normalized dry mass density and average mass growth rate: data showed in Figure 4D, with a zoom on minutes before and after the application of the hypoosmotic shock (applied at time 0). Data is represented as binned mean, A: ± SD; C: ± SEM. A: all cell lines and conditions as in Figure 2D; B, C: HeLa adherent cells with osmotic shock (n=52, N=3).

**Supplementary Figure 5 (related to Figure 5).**
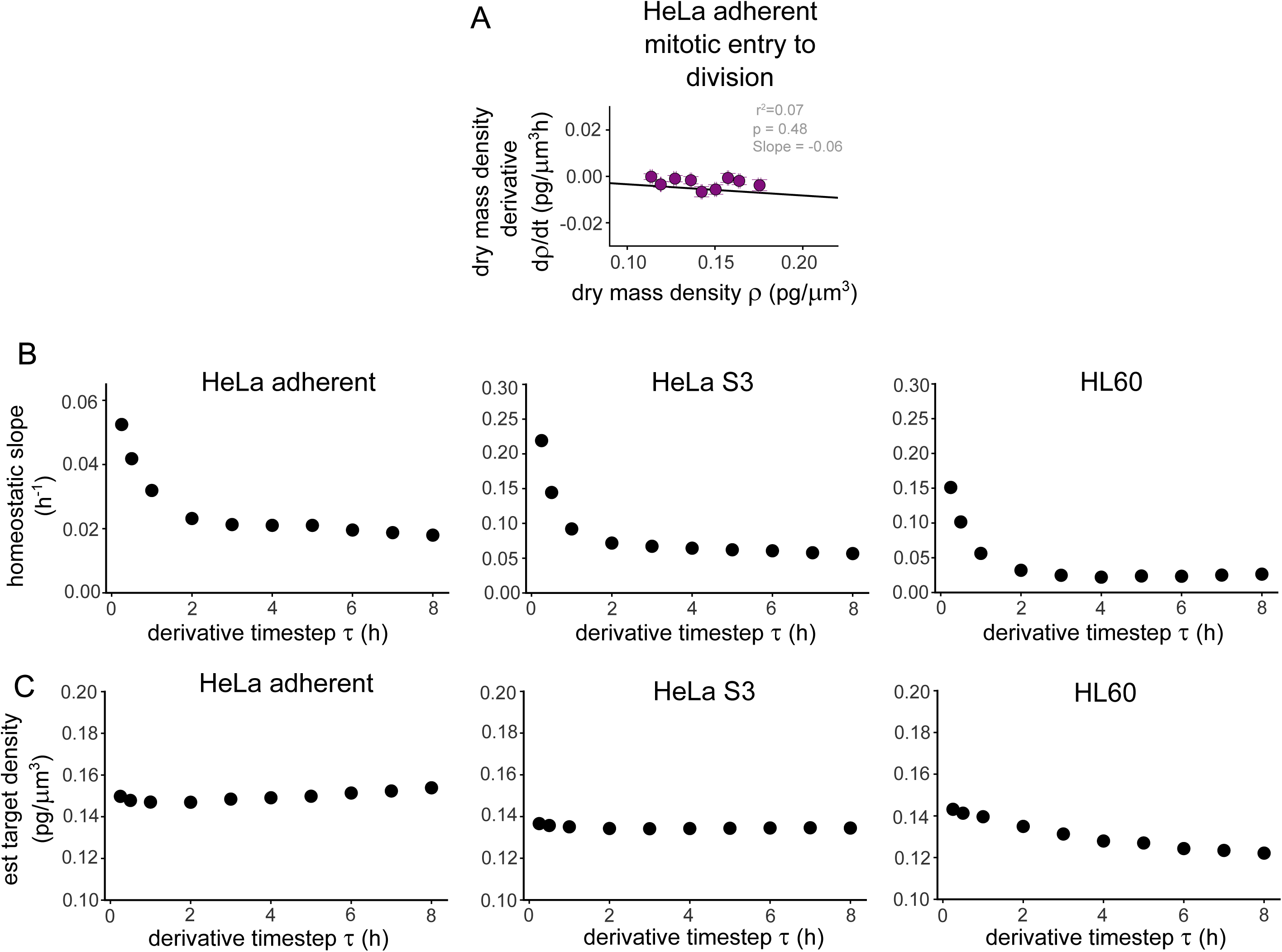
Robustness of the target density determined from time derivatives of dry mass density measured at various timescales. **(A)** Conditional average of dry mass density derivative during mitosis (between mitotic entry and division) with respect to the dry mass density at mitotic entry for HeLa adherent cells (n=140, N=3). Solid circle indicates binned mean ± SEM with equal weighted bins. Solid line indicates linear fit on the binned values (linear regression p-value = 0.46). **(B)** Slope of the dry mass density time derivative versus dry mass density plots for time derivatives measured over increasing time windows (derivative timestep τ), for (left to right): HeLa adherent, HeLa S3 and HL60. **(C)** Estimated target density for density time derivatives measured over various time windows (derivative timestep τ), for (left to right): HeLa adherent, HeLa S3 and HL60. The experimental dataset used is the complete dataset for steady growing cells, used in other figures (e.g. Fig. 2D).

**Supplementary Figure 6 (related to Figure 6).**
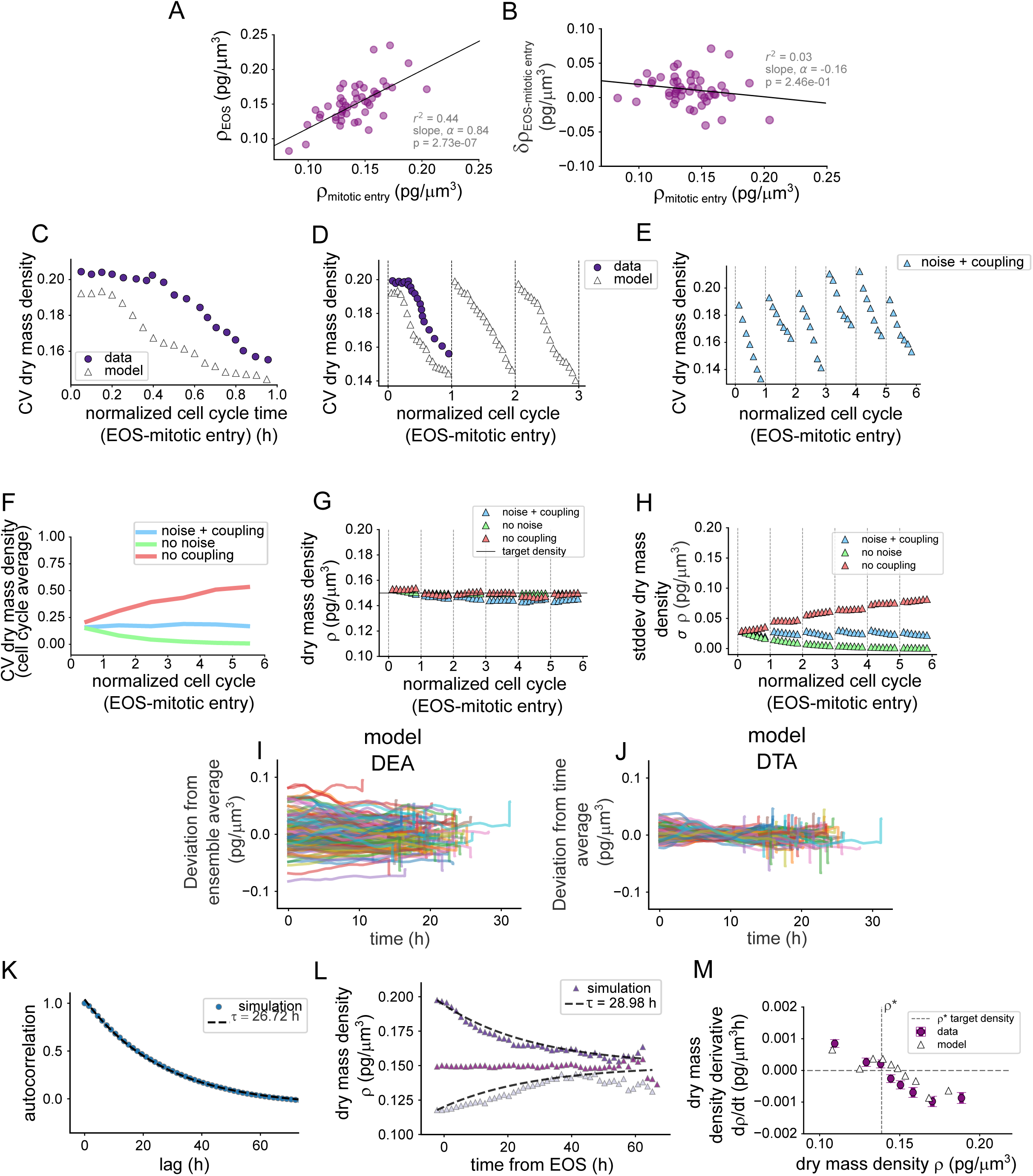
Distribution of dry mass density in a population of proliferating cultured cells, experimental data and model predictions. **(A)** Dry mass density of daughter cell at EOS versus its mother cell at mitotic entry for single HeLa adherent cells; Black line : y=x. **(B)** Dry mass density change between EOS of daughter cell and mitotic entry of its mother cell versus density at mitotic entry of its mother cell for single HeLa adherent cells. **(C)** Evolution of the CV of dry mass density across a single cell division cycle (between EOS and mitotic entry, CVs calculated on temporal bins). Solid circles: experiments; triangles: simulated cell densities from the model. **(D)** like C, but the data from simulations are shown for 3 cycles. **(E)** CV of dry mass density from simulated single cell trajectories, in the presence of noise and with homeostatic coupling (similar to Figure 6C blue triangles), shown alone with a different Y scale, to better visualize changes across each cycle. (F) CV of dry mass density from simulated trajectories, similar to Fig. 6C, but the curves are time-averaged and only the trend across cell cycles are shown and not the variation within one cell cycle. For each cell cycle, the time average of the CV across the cycle is shown (G) Mean dry mass density evolution in time, from simulated single cell trajectories, for the different model scenarios shown in Fig. 6C. (H) Standard deviation of dry mass density evolution in time, from simulated single cell trajectories, for the different model scenarios shown in Fig. 6C. (I) Ensemble average (DEA) and (J) time average (DTA) subtracted dry mass density from simulated single cells trajectories (K) Density autocorrelation for a simulation without noise due to mitosis. Autocorrelation is computed after letting the system reach equilibrium. Blue circles: simulated data; dashed black line: exponential decay with a timescale equal to 26.72 h. (L) Trend of the mean dry mass density for three groups of cells with different starting conditions for a simulation without noise around mitosis. Triangles: simulated data; dashed black line: exponential decay to equilibrium with a timescale equal to 28.98 h. (M) Target density plot of model against data (HeLa adherent cells), similar to Fig S5A. The experimental dataset used is the complete dataset for steady growing adherent HeLa cells, used in other figures (n=133 cells, N=3 independent experiments).

**Supplementary Figure 7 (related to Figure 7).**
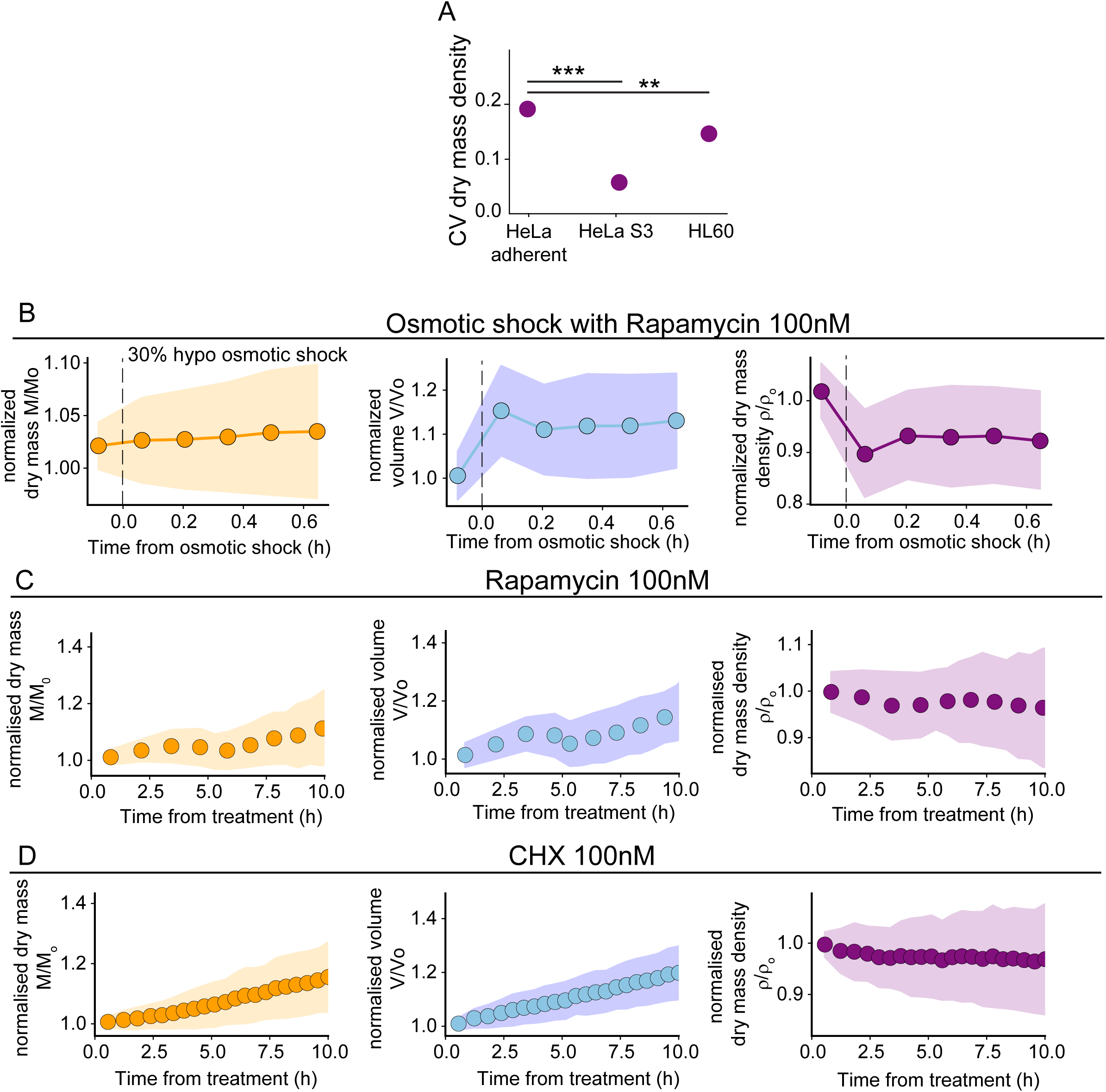
Perturbations of dry mass density homeostasis. **(A)** CV of dry mass density calculated for the 3 main cell lines used in the study (** p<0.05, *** p<0.005). **(B)** Normalised binned averages of volume (left, blue), dry mass (right, orange) and dry mass density (middle, purple) against time after a 30% hypoosmotic shock (applied at time 0), for HeLa adherent cells treated with 100nM Rapamycin (n=49, N=3). **(C-D)** Mean normalised dry mass (left, orange), volume (middle, blue) and dry mass density (right, purple), against time after treatment for HeLa adherent cells growing in (C): 100 nM rapamycin (n=65, N=3) and (D) 100 nM CHX (n=70, N=3). B, C: Data is represented as binned mean ± SEM with equal weighted bins.

**Supplementary Figure 8 (related to Figure 7).**
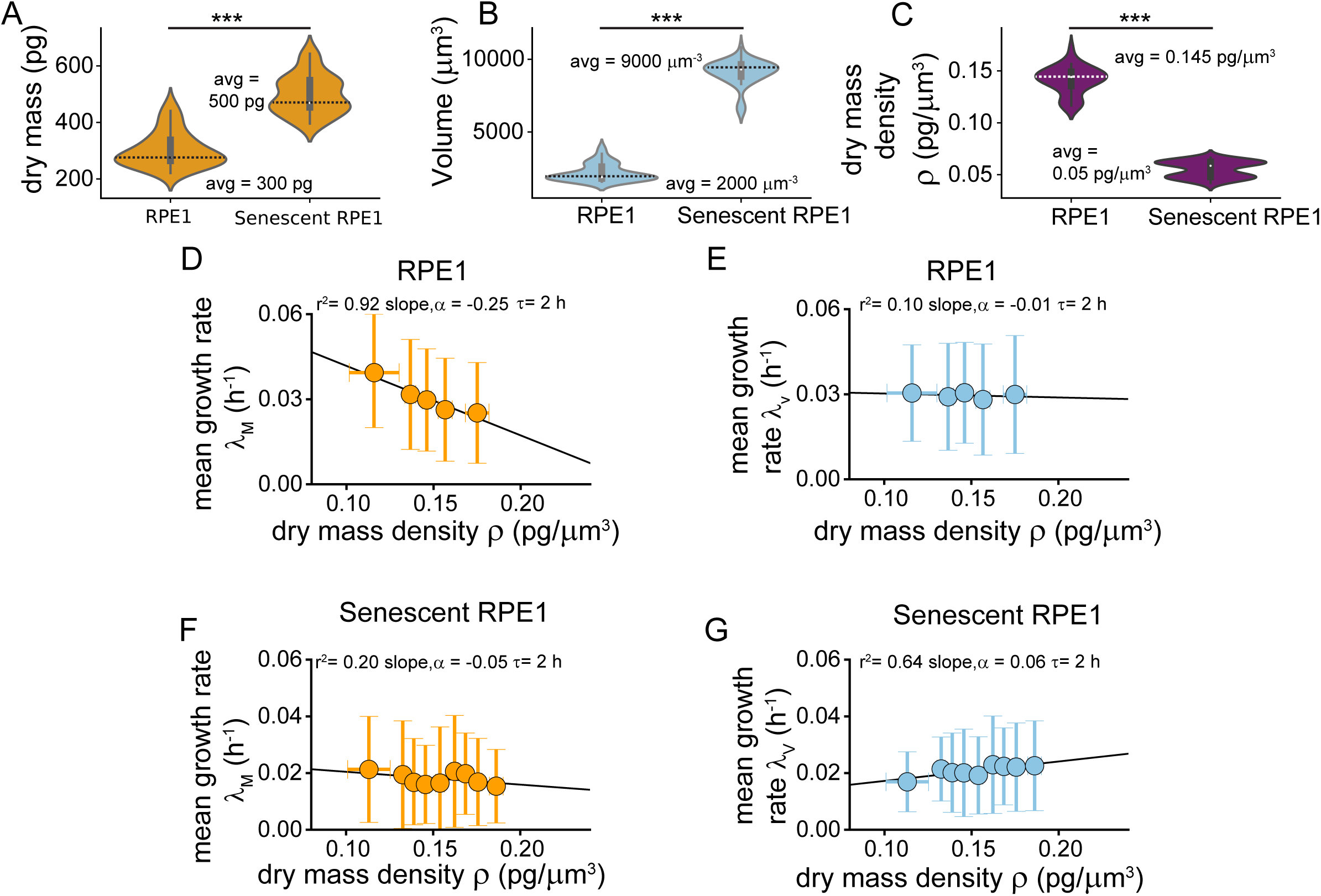
Dry mass density in RPE1 cells and RPE1 senescent cells. **(A-C)** Rainbow plot distribution of A: dry mass, B: volume, and C: dry mass density, for RPE1 control and senescent cells; average values on the plots (*** p<0.005). **(D-G)** Conditional averages of D, F: mass growth rate; E, G: volume growth rate at fixed dry mass cell density for D, E: control cells; F, G: senescent cells. Discrete derivatives are calculated for time windows of τ = 2 hours. D-G: binned mean values ± SEM with equal weighted bins, solid line: linear fit (slope and r^2^ values shown on the plots); All data in this figure: RPE1 cells (n=68, N=2) and senescent RPE1 cells (n=70, N=2)

