## Supplementary Model file for "A Dual Homeostatic Regulation of Dry Mass and Volume Defines a Target Density in Proliferating Mammalian Cells"

### Supplementary Notes on Mathematical Models for Srivastava, Calabrese *et al.*

These Supplementary Notes present in further detail all the theoretical aspects of the study, the different mathematical models that were employed, and how they were applied to analyze our data.

#### INTRODUCTION

In our study, we employed three distinct mathematical models to investigate the processes of joint dry mass and volume growth and guide our data analysis. The first model, introduced in the main text (see Fig. 3-5 of the main text) and described in sec. 1 and 2 is a physical "osmo-metabolic" model describing the average-cell behavior. This first model integrates a coarse grained model of biosynthesis, flux-balance and resource allocation [1, 2] with a physical pump-leak framework designed to describe osmotic equilibrium and ion transport [3–5]. This approach is tailored to understand the growth dynamics of the average cell within a population, providing an understanding of the typical behavior of cells in response to physiological and environmental cues. The second model, described in sec. 3 and 4 (see Fig. 5 of the main text) is a phenomenological mean-field approach used for the detection of dry-mass density homeostasis and regulation, based on two conditions: the population dry-mass density reaches and maintains a target density in the absence of perturbations, and the dry-mass density adjusts itself towards this set point when perturbed. The model also explores the relationship between mass and volume growth rates and cell dry-mass density to ensure the existence of a homeostatic value. To complement and extend our analysis, particularly in examining single-cell fluctuations, we also employed a stochastic phenomenological model of joint volume and macromolecular dry-mass growth in single cell trajectories (see Fig. 6 of the main text). This second model, described in sec. 5 and 6 offers a different perspective, focusing on individual cell behavior to quantify dynamic variations form typical behavior and cell-specific features that are masked in population-level analyses.

#### 1. OSMO-METABOLIC MODEL OF JOINT DRY MASS AND VOLUME GROWTH

This section describes the osmo-metabolic model of joint macromolecular mass and volume growth used in the main text (in particular see Fig. 3). This model includes a description of flux-balance and biosynthesis [1, 2] jointly with a physical pump-leak model describing osmotic equilibrium and ion transport [5]. We note that this model applies to average quantities and we refer the reader to section 5 for a phenomenological model that describes single-cell behavior, which we applied to analyze single-cell fluctuations in our data.

We briefly describe the main model ingredients and assumptions. In our simplified treatment, we approximate the macromolecular dry mass measured by quantitative phase microscopy as the protein (dry) mass (i.e., $M \approx M_p$). This can be justified by the fact that proteins take from 50% to 70% of the overall dry mass [6]. Therefore, in our model macromolecular dry mass growth is assumed to be a consequence of protein translation.

Cell volume is divided into its osmotically inactive and active (accessible) components, i.e.

$$V = V_{\mathrm{i}} + V_{\mathrm{e}} \, . \tag{S1}$$

For a simple system, the osmotically inactive volume can be interpreted as the total volume of all nonsolvent molecules in the compartment [5, 7]. For a complex system like a cell, this volume is difficult to interpret specifically, but is clearly due to entire compartments that contain water and are very complicated to describe in terms of entropy. Hence, $V_{\mathrm{i}}$ contains water and all the macromolecular dry mass that does not contribute to osmotic pressure (e.g. immobilized compartments or any cell components whose entropy is reduced for various reasons), $M_{\mathrm{i}}$. By osmotic balance, our theory assumes that the water content of the cell setting $V_{\mathrm{e}}$ is proportional to the molar amount of impermeant osmolytes, and that the dominant non-ionic osmolytes of this kind are small osmolytes (e.g. free amino acids and/or their metabolic precursors) [5].

Gicen these arguments, the model assumes that volume is set by $V_{\mathrm{e}} \propto A$, where $A$ is the total content of these osmotically dominant free species in the cell, likely of metabolic origin. For simplicity, we will refer to these species as “amino acids” in the following, but it is important to clarify that their definition might be more general. Indeed, even if a strong dominance of amino acids as non-ionic small osmolytes can be debated, the model would not change much, as other small osmolytes such as metabolite precursors are all produced/accumulated under by metabolic enzymes or transporters. Under these conditions, they will be subject to the same general relations between their accumulation rate and the amount of proteins in the cell specified by our model. We will derive this relationship in the next sections. To make the notation uniform, we write $V_{\mathrm{e}} \propto M_a/m_a$, where $M_a$ is the total free amino acid mass and $m_a$ is mass of a single amino acid. The following paragraphs define the mass dynamics of our model.

**Macromolecular dry mass dynamics in the osmo-metabolic model of cell growth.** Dry mass comprises various components such as proteins, lipids, carbohydrates and various other metabolites, DNA, and RNA. Out of these, total protein content is the most abundant, constituting more than 50% of the dry mass across diverse species [6]. The second most abundant component varies depending on the organism. For example, RNA comprises 20% of the dry mass in *E. coli*, while carbohydrates make up 30% in *S. cerevisiae*, and lipids account for 15% of mammalian cells' dry mass [6].

Given the complexities inherent in accounting for the synthesis of all cellular components, we focus on the production of two fundamental constituents, proteins, which dominate macromolecular dry mass, and free amino acids (in the wide sense specified above), which we assume to be the most abundant non-ionic small osmolyte. As we will see below, under these assumptions amino acid dynamics sets cell volume in our model, which provides a link between mass and volume production [5].

We define $J_P$ as the total protein production flux (number of proteins produced per unit time), and $J_A$ as the total amino acid production flux (number of amino acids produced per unit time). Biologically, these fluxes can depend on several biosynthesis parameters inside the cell, including ribosomes concentration, concentrations of amino acid transporters and enzymes, translation rates, etc [1, 2]. For simplicity and economy of parameters, we did not describe in detail all these dependencies, but these details can be incorporated in our model as more data become available [2, 8–10]. Let us define $P$ and $A$ as the number of proteins and amino acids. By the definition of the fluxes $J_P$ and $J_A$, we can write

$$\frac{dP}{dt} = J_P \tag{S2}$$

and

$$\frac{dA}{dt} = J_A - L\,\frac{dP}{dt} = J_A - L\,J_P\,, \tag{S3}$$

where the second equation takes into account the fact that for each protein produced, $L$ amino acid are incorporate inside a protein where $L$ is the average protein length measured in amino acids.

To convert these equations to dry masses $M_P$ and $M_A$ we can use the dry mass of a single amino acid $m_A$, to obtain $M_P = P\,L\,m_a$ and $M_A = m_a A$. We can also re-define the fluxes as mass fluxes, $J_P \to m_P J_P$ and $J_A \to m_A J_A$, obtaining

$$\frac{dM_P}{dt} = J_P \tag{S4}$$

and

$$\frac{dM_A}{dt} = J_A - \frac{dM_P}{dt} = J_A - J_P\,. \tag{S5}$$

Conveniently, both the protein length parameter $L$ and the masses disappear in this equation.

Next, we assume that since they rely on growing amounts of catabolic and translation machinery, the total protein and amino acid fluxes are proportional to the total protein mass through some mass-specific fluxes $j_P$ and $j_A$

$$J_P = j_P M_P\,; \qquad J_A = j_A M_P \tag{S6}$$

$j_P$ and $j_A$ are the specific protein and amino acid fluxes (fluxes per total protein mass). Unless specified, we shall refer in the following to $j_P$ and $j_A$ as "the fluxes". From a mechanistic point of view, such fluxes are likely to be complicated functions of several parameters, but for our purposes we will not need to make this dependence explicit.

Consequently, we obtain the following equations for production of protein mass and amino acid mass,

$$\frac{dM_P}{dt} = j_P M_P \tag{S7}$$

and

$$\frac{dM_A}{dt} = (j_A - j_P)M_P\,. \tag{S8}$$

Next, we link the equations to the total dry mass $M$. By definition, the total dry mass is the sum of the protein dry mass, the free amino acid dry mass and all other components, which we term $M_O$. We make the further assumptions that $\frac{dM_O}{dt} = j_O\,M_P$, and that the amino acid pool has a smaller overall mass compared to the other two contributions. This assumption appears justified [5]: assuming that the concentration of free amino acids is approximately 100 mM and that of protein, with an average length of 400 amino acids, is 3 mM, there can be roughly ten times as many amino acids present in proteins as in free form. Hence,

$$M = M_P + M_A + M_O \approx M_P + M_O\,, \tag{S9}$$

and

$$\frac{dM}{dt} = (j_P + j_O)\,\frac{M_P}{M}\,M\,. \tag{S10}$$

With some algebra, and considering Eq. (S6), these equations imply that assuming steady-state growth $\frac{M_P}{M} = \frac{j_P}{j_O + j_P}$. Hence, we can write the equations that regulate protein and amino-acid mass growth as

$$\frac{dM}{dt} = j_P \, M \tag{S11}$$

and

$$\frac{dM_A}{dt} = (j_A - j_P) \, f_P \, M \,, \tag{S12}$$

where we have defined $f_p := \frac{M_P}{M}$ (protein mass fraction).

**Volume dynamics in the osmo-metabolic model of cell growth.** As mentioned above, the model divides cell volume into osmotically active and passive volume components. We assume that osmotically inactive volume production is enslaved to dry mass production, hence we use the equations derived in the previous section to describe the main processes that drive changes in this part of cell volume. Conversely, the "osmotically active" volume is related to amount of water flowing in and out of the cell, hence to total cell volume. Note that since most of the volume is active, researchers have noted before that the most basic problem of cell volume regulation is to understand how water flows in and out of cells [4, 11]. We adopt a minimal description of the basic physical principles underling the dynamics of these water fluxes. Our approach in the the following directly follows from previous work contained in refs. [4, 5, 11].

Note that what we call osmotically active and inactive volumes do not coincide with the "dry" (water free) and "wet" volume of the cell [12, 13], although this terminology is sometimes adopted [5]. For example, the osmotically inactive volume can contain water and a lot of the dry volume (e.g. proteins) is actually osmotically active.

The mobility of water in and out of a cell depends on the permeation through the plasma membrane, a process regulated by aquaporin channels [11]. These channels permit the passage of water, glycerol, and some small molecules, but not ions. Given the assumption that the channels operate without ATP consumption, the influx of water correlates with the difference between two chemo-mechanical driving forces: differences in osmotic pressure ($\Pi$) and hydrostatic pressure ($P$) between the interior and the exterior of the cell [11]. The filtration coefficient of the membrane ($L_p$), influenced by the density of aquaporins, is the proportionality factor between the flow rate and the pressure difference, which varies across different cell types.

Osmotic pressure can be calculated based on the concentration of osmotically active molecular species in and out of the cell. Determining the difference in hydrostatic pressure needs a mechanical model of the cell, focusing on the balance of mechanical forces on its contour. This balance stems from a complicated interaction that may involve tension at the cell wall, membrane, and cortex, depending on the cell type. The hydrostatic pressure difference thus associates the intracellular volume with its mechanical properties through a force balance at the cell contour [11]. Here, we are concerned with mammalian cells, and we can safely assume that the small contribution due to membrane tension can be neglected [7, 14, 15]. Indeed, the cytoskeletal forces that can be exerted at the cell cortex are known from the literature and can be one to the order of magnitude smaller than those required to press water out of the cell. Hence, we can summarize the basic physics illustrated above with the following equation

$$\frac{dV}{dt} = \frac{dV_\mathrm{i}}{dt} + L_p \left( \Delta p - \Delta \Pi \right) \,. \tag{S13}$$

Importantly, the active volume term can change fast (seconds for water flows through the membrane and minutes for ion fluxes [16]) compared to the inactive volume (which doubles in a cell cycle of about 20-24 hours). At the same time, the basic physics of osmosis dictates the imbalance is compensated within the volume-relaxation timescale. As we are interested in volume expansion at the timescale of minutes/hours, we employ a quasi-static approximation [5, 11] whereby

$$\Delta p \approx \Delta \Pi \,. \tag{S14}$$

We stress that this assumption does not mean that active volume stays constant, but rather than the mechanical-equilibrium (active) volume $V_\mathrm{e}^*$, will vary on minutes-hours timescales according to the variation of the relevant variables $(x_1, x_2, ..)$ inside the cell, mainly the osmolytes that set the the osmotic pressure. As $(x_1, x_2, ..)$ change slowly compared to the equilibration timescale, we are merely assuming that the volume then grows by progressively taking a series of equilibrium values. Next, we derive an expression of the quasi-static equilibrium value of the volume as a function of these variables.

The imbalance of the osmotic pressure is proportional to the difference of the osmolyte concentration between the interior and the exterior of the cell. We separate the osmolytes in permeant $n$ and impermeant ones $X$. Impermeant osmolytes cannot diffuse through the cell membrane, while permeant ones may cross the cell membrane and at steady state the incoming flux into the cell must equate the outgoing flux. Most of the permeant osmolytes are ions and the model assumes that all of them are. We divide them into negatively $n_-$ and positively $n_+$ charged ones. Impermeant osmolytes include proteins, DNA, RNA amino acids and metabolites of various kinds. These last species can be actively imported for nutrient uptake. Following the van't Hoof equation [17], the osmotic pressure of a species follows an ideal-gas like law $\Pi V = N k_b T$, and the osmotic pressure difference can be expressed as

$$\Delta\Pi = k_b T \left[ \left( \frac{n_+ + n_- + X}{V_\mathrm{e}} \right) - c_\mathrm{osm}^\mathrm{ext} \right] . \tag{S15}$$

To achieve mechanical equilibrium (Eq. (S14)), water flows in or out of the cell until the equation $\left(\frac{n_+ + n_- + X}{V_\mathrm{e}}\right) = c_\mathrm{osm}^\mathrm{ext}$ holds. Therefore, $V_\mathrm{e} \propto n_+ + n_- + X$.

The next crucial step is to realize that $n_+$, $n_-$ and $X$ are not independent variables. Following the classic Pump-Leak model [3, 5, 18] we can obtain two conditions that connect the two variables, electroneutrality of the cell and ionic flux balance. Mathematically, these conditions can be expressed as

$$n_+ - n_- - zX = 0 \tag{S16}$$

and

$$\frac{n_+}{V_\mathrm{e}} \frac{n_-}{V_\mathrm{e}} = \alpha_0 \, \left(c_\mathrm{osm}^\mathrm{ext}\right)^2 . \tag{S17}$$

The first equation simply sums all the charges inside the cell, with $z$ being the average charge of the impermeant osmolytes. The second equation is a consequence of the electrochemical gradient to due diffusion of charged particles and active pumping of ions on the cell part. $\alpha_0$ represent the pumping efficiency, which can take values between 0 (infinite pumping of anions outside of the cell) and 1 (no pumping)[1]. We will consider the limit of infinite pumping, which can be argued to be a good approximation of physiological situation where the cell interior is dominated by cations acting as counterions of charged macromolecules (see the appendix of ref. [5] for further details). By using these relationships to express the variables $n_+$ and $n_-$ in terms of $X$, we obtain

$$V_\mathrm{e} = \frac{(z+1)}{c_\mathrm{osm}^\mathrm{ext} + \frac{\Delta p}{k_b T}} X . \tag{S18}$$

Note that this equation can be interpreted in a causal manner only if the hydrostatic pressure difference $\Delta p$ satisfies certain conditions. Indeed, $\Delta p$ is generally a function of the active (excess) volume $V_\mathrm{e}$. This means that the equation above provides the volume only implicitly. For example, Laplace's law relates the pressure difference to the membrane/wall tension $\gamma$ and the radius of the cell $R$, assuming spherical geometry. More precisely, Laplace's law takes the following form: $\Delta p = \frac{\gamma}{R}$. As the tension is related to the extent the cell surface is stretched, $\gamma \propto S \propto V_\mathrm{e}^{2/3}$, while $R \propto V_\mathrm{e}^{1/3}$. Therefore, the pressure difference $\Delta p$ generally depends on the active volume. A fully general picture should carefully model $\Delta p$, which includes taking into account the specific cell shape and membrane/wall tension regulation [19, 20].

Here, by assuming $\Delta p \approx 0$ (negligible hydrostatic pressure and cell tension contributions) compared to osmotic pressure, which is the case for mammalian cells [14], we can avoid this problem. Hence, we proceed to write the dynamics of the total volume in the case of negligible hydrostatic pressure,

$$\frac{dV}{dt} = \frac{dV_\mathrm{i}}{dt} + \frac{(z+1)}{c_\mathrm{osm}^\mathrm{ext}} \frac{dX}{dt} . \tag{S19}$$

Importantly Eq. (S19) couples different volume- and dry-mass related quantities. We assume that changes in inactive volume are subdominant and linked to changes in protein levels (dry mass). In other words, we assume that the macromolecular dry mass of the osmotically inactive

[1]We note that in the case of no pumping, osmotic equilibrium cannot hold according to these equations - water flows uncontrollably and the cell eventually lyses.

part of the cell is proportional to the total macromolecular dry mass, $M_\mathrm{i} = f_\mathrm{i} M$, where $M_\mathrm{i}/\rho_\mathrm{i} = V_\mathrm{i}$, and $\rho_\mathrm{i}$ is the macromolecular dry-mass density of the inactive compartment of the cell. We note that $\rho_\mathrm{i}$ can be less than the density of water, since the osmotically inert cell compartments can contain water. Consequently, we can write

$$\frac{dV_\mathrm{i}}{dt} = \frac{1}{w_\mathrm{i}'} \frac{dM}{dt} \,, \tag{S20}$$

where $w_\mathrm{i}' = \rho_\mathrm{i}/f_\mathrm{i}$ has units of mass over volume.

On the other hand, the model assumes that changes in active volume are proportional to changes in the number of impermeant osmolytes $X$, which in principle contain proteins, RNA, DNA and metabolites (the components that make up the dry mass of the cell). We further assume that the number of impermeant osmolytes $X$ is dominated by free amino acids (counted together with their intracellular precursors) [5]. Under this assumption dry mass growth is driven mainly by increase in protein content, and volume growth is controlled by free amino-acid content. If $A$ is the total free amino-acid content (number of molecules), then we write to a first approximation,

$$\frac{dX}{dt} \approx \frac{dA}{dt} \,. \tag{S21}$$

By combining this equation with the equations for macromolecular mass production, $\frac{dM}{dt} = j_P M$ and amino acids dynamics, $\frac{dA}{dt} = \frac{1}{m_a}\frac{dM_A}{dt} = \frac{1}{m_a}(j_A - j_P)M_P$, we obtain an equation for the exponential growth rate of volume $\lambda_V$ which is the main variable of interest, and which drives dry-mass density changes through volume,

$$\frac{dV}{dt} = \frac{1}{w_\mathrm{i}'}\frac{dM}{dt} + \frac{(j_A - j_P)\, f_P}{w_\mathrm{osm}} M \,. \tag{S22}$$

Here, $w_\mathrm{osm} := \frac{m_a c_\mathrm{osm}^\mathrm{ext}}{z+1}$, which has units of mass over volume. We note that in this framework, since amino-acid metabolic flux sets volume growth, a model of amino acid metabolism regulation is in principle required to predict the volume dynamics.

#### 2. OSMO-METABOLIC MODEL: RESULTS

##### A. The osmo-metabolic model predicts a homeostatic relationship between the volume growth rate and the dry mass density

Our first result from the osmo-metabolic model is to obtain a relationship between volume growth and dry-mass density. This prediction is verified in Fig. 3FG in the main text. We find that suspension cells follow the linear trend between volume growth rate and macromolecular dry-mass density predicted by the model (Fig. 3F in the main text), while adherent cells show an average flat trend (Fig. 3G in the main text). As discussed in the main text, we interpret this discrepancy as due to confounding factors related to cell adhesion, mainly the fact that active and passive forces related to adhesion affect the osmo-mechanical fluctuations, masking the positive correlation due to small osmolytes [16].

Let us now proceed and derive this result from the model. To summarize the model definition, we have obtained the following mean-field equations for the dry mass and the volume,

$$\frac{dM}{dt} = j_P\, M \tag{S23}$$

and

$$\frac{dV}{dt} = \frac{j_P}{w_\mathrm{i}'} M + \frac{(j_A - j_P)\, f_P}{w_\mathrm{osm}} M \tag{S24}$$

Since the rate of change of the dry mass is directly proportional to the dry mass itself, the first equation describes exponential mass growth. Moreover, the rate of change of the volume is also proportional to the dry mass, which increases exponentially - consequently, the dry volume also increases exponentially. The proper kinetic parameters to describe growth are therefore the specific growth rates

$$\lambda_M := \frac{1}{M}\frac{dM}{dt} \,; \qquad \lambda_V := \frac{1}{V}\frac{dV}{dt} \,, \tag{S25}$$

which we can immediately write as

$$\lambda_M = j_P \tag{S26}$$

and

$$\lambda_V = \left( \frac{j_P}{w_{\rm i}^{'}} + \frac{(j_A - j_P)\, f_P}{w_{\rm osm}} \right) \rho \tag{S27}$$

where $\rho$ is the dry-mass density $\frac{M}{V}$. We note that the second equation suggests that the volume growth rate of a single cell could be proportional to its dry-mass density, provided the other parameters are fixed and do not depend on density.

#### B. The osmo-metabolic model predicts a homeostatic dry-mass density for steady-growing cells

Our second result is proving that the osmo-metabolic model naturally reaches a steady-state where $\lambda_M = \lambda_V$ (Fig. 1 and Fig. 5A in the main text). To see this, we write $\frac{d\rho}{dt}$ using the definition of $\rho$ and the derivative chain-rule. This gives

$$\frac{d\rho}{dt} = \rho\, (\lambda_M - \lambda_V) = \rho \left[ j_P - \left( \frac{j_P}{w_{\rm i}^{'}} + \frac{(j_A - j_P)\, f_P}{w_{\rm osm}} \right) \rho \right] . \tag{S28}$$

Since in this equation $\lambda_V$ is proportional to $\rho$, the system reaches a fixed point $\rho^*$ where the density attains a constant value,

$$\rho^* = \frac{j_P}{\frac{j_P}{w_{\rm i}^{'}} + \frac{(j_A - j_P)\, f_P}{w_{\rm osm}}} . \tag{S29}$$

At this value of the density, the dry mass and the volume growth rates are identical, $\lambda_M = \lambda_V$. We call this value, which is the predicted steady-state growth rate of the model $\lambda^*$,

$$\lambda^* = j_P = \frac{j_P}{w_{\rm i}^{'}}\, \rho^* + \frac{(j_A - j_P)\, f_P}{w_{\rm osm}}\, \rho^* \tag{S30}$$

Since $j_P$ is equal to the instantaneous growth, this is only an implicit relationship between the equilibrium growth rate and dry-mass density. With some algebra, we can make it explicit:

$$\lambda^* = j_A\, \frac{f_P \frac{\rho^*}{w_{\rm osm}}}{1 - \frac{\rho^*}{w_{\rm i}^{'}} + f_P \frac{\rho^*}{w_{\rm osm}}} \tag{S31}$$

Finally, we can give a rough estimate of the quantities involved. Let us recall that the definition $w_{\rm osm} := \frac{m_a c_{\rm osm}^{\rm ext}}{z+1}$. The concentration of external osmolytes is roughly 300 mM while $z$ is roughly equal to 0.1 [5, 21], where for the latter we used the fact the domaninant osmolites for HeLa is glutamine, which is neutral. The mass of a single amino acid is 100 Da. We expect $f_P \sim 0.5$, as roughly half of the dry mass is made of protein. Therefore, $w_{\rm osm} = 2.72\ 10^{-2}$ g/mL, which implies $f_P \frac{\rho^*}{w_{\rm osm}} \approxeq 2.75$ where we used the equilibrium value of $\rho^* \approxeq 0.15$ g/mL observed in the data. We can estimate $f_{\rm i}$ from so-called Ponder plots [16], which give the cell volume changes upon osmotic shocks. Before the shock $\Pi_0(V_0 - V_{\rm i}) = N k_b T$, and after the shock $\Pi_f(V_f - V_{\rm i}) = N k_b T$, so that

$$\frac{V_f}{V_0} = \left(1 - \frac{V_{\rm i}}{V_0}\right) \frac{\Pi_0}{\Pi_f} + \frac{V_{\rm i}}{V_0} . \tag{S32}$$

Using the fit of $\frac{V_f}{V_0}$ *vs* $\frac{\Pi_0}{\Pi_f}$ from data in ref. [16], we get $V_{\rm i}/V_0 \simeq 0.3$. Since $M_{\rm i} = \rho_{\rm i} V_{\rm i}$ and $M = \rho V$, we can use this observation to estimate $f_{\rm i} \simeq 0.3\ \rho_{\rm i}/\rho$, hence $w_{\rm i}^{'} = \rho_{\rm i}/f_{\rm i} \simeq \rho/0.3 \approx 0.5$g/mL (given our measurements for the homeostatic density of about 0.15 g/mL). Therefore $\lambda^* \sim j_A \frac{2.75}{1-0.15/0.5+2.75} \sim j_A\ 0.8$.

Consequently, we note $j_A$ should have the same order of magnitude of the observed growth rate $\lambda^*$. A simple argument assuming balanced exponential growth also argues in favor of this observation. From Eq. S12 in conjunction with balanced exponential growth

$$\frac{dM_A}{dt} = (j_A - j_P)\, f_P\, M = \lambda^* M_A\, , \tag{S33}$$

or equivalently

$$j_A = \lambda^* \left(1 + \frac{M_A}{M_P}\right) , \tag{S34}$$

where we used $j_P = \lambda^*$ and $M_P = f_P M$ at equilibrium. The key observation is that $M_P \gg M_A$, that is, the protein dry mass is much greater than the free amino acid dry mass. Therefore, the ratio disappears and $j_A \approx \lambda$.

##### C. Volume growth may persist under full inhibition of translation due to non-vanishing amino-acid flux

A subsequent prediction of this model concerns the behavior of volume growth when biosynthesis is inhibited. Under full inhibition of translation, the protein-production flux $j_P$ falls to zero. If translation inhibition is strong and rapid, as it is case for the sudden treatment with a fully inhibitory dose of translation-targeting drugs such as Cycloheximide, we can assume that (for some time) every other parameter stays constant. According to equation S27 the osmo-metabolic model predicts that the volume keeps growing under this perturbation due to the continued influx of amino-acids which are not employed in protein translation, as

$$\lambda_V = \left(\frac{j_P}{w_i'} + \frac{(j_A - j_P)\, f_P}{w_{osm}}\right) \rho|_{j_P=0} = \frac{j_A\, f_P}{w_{osm}}\, \rho > 0\,. \tag{S35}$$

We expect that $j_A$ may change rapidly in response to this stress. However, it is not obvious how quickly this could happen, e.g., depending if the response, which must be regulated, is transcriptional or post transcriptional. Our data in Fig. 3AB in the main text show that the volume keeps on growing for few hours before finally ceasing to increase. Hence, assuming that the osmo-metabolic model correctly describes the data, this response is not very rapid. Fig. 3AB in the main text show that a model where $j_A$ does not change ("osmo-metabolic model" in Fig. 3) is falsified by the data, while a model where $j_A$ can respond to the treatment ("osmo-metabolic model w/feedback" in Fig. 3D) can fit the data. See below for a detailed discussion of the regulatory responses after full inhibition of translation within the osmo-metabolic growth model.

##### D. Inhibition of translation causes a sudden increase in the volume growth rate due to flux mismatch

Another observation of Fig. 3B in the main text is the fact that volume growth rate immediately increases upon full inhibition of translation. In order to understand this result within the osmo-metabolic growth model, we proceed by quantifying the flux mismatch predicted by the osmo-metabolic model, in absence of regulatory response. Targeting translation corresponds to decreasing the protein flux $j_P$. In particular, as discussed above, our experiments lead us to consider the simple case where $j_P$ becomes zero instantaneously. Using equation S22 and S27, the model describes this perturbation with the following equations for mass and volume,

$$\frac{dM}{dt} = \begin{cases} \lambda^*\, M & t < 0 \\ 0 & t \geq 0 \end{cases} \tag{S36}$$

and

$$\frac{dV}{dt} = \begin{cases} \lambda^*\, V & t < 0 \\ \frac{j_A\, f_P}{w_{osm}}\, M & t \geq 0\,. \end{cases} \tag{S37}$$

Specifically, the model predicts that the macromolecular mass of the cell (assumed to be dominated by proteins) stops growing as translation halts, while cellular volume continues to increase. This is due to the fact that in the osmo-metabolic model volume growth is linked to amino acid flux $j_A$ rather than protein production, i.e. ($j_P$), and $j_A$ is not directly inhibited by anti-translation drugs. We also note that while volume growth does not stop, the mode of growth does change. Since the mass $M$ is now constant in equation S22 for $t \geq 0$, the volume grows linearly instead of exponentially. This qualitative change of trend is visible in Fig. 3A in the main text. A similar argument was used in ref. [5] to explain the experimental observation that long-term G1 arrest of yeast cells leads to a linear growth regime when the DNA gets diluted and gene content becomes limiting for growth [22].

Using Eq. S30 to express the term $\frac{j_A f_P}{w_{\rm osm}}$ as a function of the equilibrium growth rate $\lambda^*$, we obtain $\frac{j_A f_P}{w_{\rm osm}} = \frac{\lambda^*}{\rho^*}\left(1 - \frac{\rho^*}{w_{\rm i}'} + f_P \frac{\rho^*}{w_{\rm osm}}\right)$. Consequently, Eq. S37 gives

$$\frac{dV}{dt} = \begin{cases} \lambda^* \, V(t) & t < 0 \\ \lambda^* \, \left(1 - \frac{\rho^*}{w_{\rm i}'} + \frac{f_P \, \rho^*}{w_{\rm osm}}\right) \, \frac{M}{\rho^*} & t \geq 0 \, . \end{cases} \tag{S38}$$

We can further simplify this equation by noting that $\frac{M}{\rho^*} = V(t=0)$ for $t \geq 0$, as the mass $M$ remains constant at the value it had at the time of treatment, $t = 0$, and does not increase over time.

$$\frac{dV}{dt} = \begin{cases} \lambda^* \, V(t) & t < 0 \\ \lambda^* \, \left(1 - \frac{\rho^*}{w_{\rm i}'} + \frac{f_P \, \rho^*}{w_{\rm osm}}\right) \, V(t=0) & t \geq 0 \, . \end{cases} \tag{S39}$$

Eq. (S39) shows clearly that the model predicts a discontinuity in the volume growth rate at $t = 0$ (from $\lambda^*$ just before treatment to $\lambda^* \left(1 - \frac{\rho^*}{w_{\rm i}'} + \frac{f_P \rho^*}{w_{\rm osm}}\right)$ right after). The jump $\Delta\lambda$ is then

$$\Delta\lambda := \lambda_V^{\rm post} - \lambda_V^{\rm pre} = \lambda^* \, \rho^* \left(\frac{f_P}{w_{\rm osm}} - \frac{1}{w_{\rm i}'}\right) \, . \tag{S40}$$

Note that the sign of the jump can be positive or negative, and depends on the factor $\left(\frac{f_P}{w_{\rm osm}} - \frac{1}{w_{\rm i}'}\right)$. In Fig. 3, we fit this parameter from the experimental curves obtained by treating HeLa cells with CHX, a well-known translation inhibitor.

Importantly, the sign of this jump depends on the difference $\left(\frac{f_P}{w_{\rm osm}} - \frac{1}{w_{\rm i}'}\right)$. This result has a simple interpretation. As mass growth stops, there are two conflicting effects on volume growth within this model: (i) since part of the volume is osmotically inactive, the volume growth rate decreases due to the fact that the growth of this component is stopped (the term $-\frac{1}{w_{\rm i}'}$), while (ii) on the other hand, since protein synthesis stops, amino acids accumulate, which causes an increase in the growth rate (the term $\frac{f_P}{w_{\rm osm}}$). The difference between these two terms determines which effect dominates. Note that in our data $\lambda_V^{\rm post} > \lambda_V^{\rm pre}$, which is consistent with the model if $\frac{f_P}{w_{\rm osm}} > \frac{1}{w_{\rm i}'}$.

We can turn to a numerical estimate to see whether this relationship is empirically grounded. As before, we use $w_{osm} := \frac{m_a c_{\rm osm}^{\rm ext}}{z+1}$. The concentration of external osmolytes is roughly 300 mM while $z$ is roughly equal to 0.1 [5, 21]. The mass of a single amino acid is 100 Da, therefore, $w_{osm} \approx 2.72 \cdot 10^{-2}$ g/mL. We expect $f_P \approx 0.5$, as roughly half of the dry mass is made of protein, therefore $\frac{f_P}{w_{\rm osm}} \approx 18$ mL/g. Finally, if we take $w_{\rm i}' \approx 0.5$ g/mL as discussed previously, we get $\frac{1}{w_{\rm i}'} \approx$ 2 mL/g. Consequently, $\left(\frac{f_P}{w_{\rm osm}} - \frac{1}{w_{\rm i}'}\right) \approx= (18-2)$ mL/g $\approx 16$ mL/g. This implies the growth rate should increase right after the shift. The overall jump should be $\lambda^* \, \rho^* \, (16$ mL/g$) \approx 0.07$ h$^{-1}$. This number has the correct sign and is within the correct order of magnitude, although it is higher than the observed jump by a factor of 4. We believe this should be taken as a reasonable order-of-magnitude estimate, but possible sources for this discrepancy are an overestimate of $w_{\rm i}'$ or an underestimate in $w_{\rm osm}$, given the uncertainty in these parameters.

##### E. The model predicts cell-volume dynamics across the translation-inhibitory treatment

Using the above considerations, the volume dynamics across the translation-inhibitory treatment can be expressed as

$$\frac{dV}{dt} = \begin{cases} \lambda^* \, V(t) & t < 0 \\ (\lambda^* + \Delta\lambda) \, V(t=0) & t \geq 0 \, , \end{cases} \tag{S41}$$

Therefore, the quantitative volume growth after translation has stopped depends only on the value of the steady-state growth rate $\lambda^*$ and the jump $\Delta\lambda$ at the shift. The above equation can integrated in a straightforward way to obtain the time evolution of the volume $V(t)$,

$$\frac{V(t)}{V(t=0)} = \begin{cases} e^{t\lambda^*} & t < 0 \\ [1 + t\ (\lambda^* + \Delta\lambda)] & t \geq 0\,. \end{cases} \tag{S42}$$

By using the definition of the growth rate $\lambda_V = \frac{1}{V}\frac{dV}{dt}$, the osmo-metabolic model also predicts its time evolution across the translation inhibitory treatment,

$$\lambda_V = \begin{cases} \lambda^* & t < 0 \\ \frac{\lambda^* + \Delta\lambda}{[1 + t\ (\lambda^* + \Delta\lambda)]} & t \geq 0\,, \end{cases} \tag{S43}$$

as well as the evolution of the dry-mass density $\rho(t) = \frac{M(t)}{V(t)}$,

$$\rho(t) = \begin{cases} \rho^* & t < 0 \\ \frac{\rho^*}{1 + t\ (\lambda^* + \Delta\lambda)} & t \geq 0\,. \end{cases} \tag{S44}$$

Therefore, upon translation arrest the volume grows as $V(t) \propto t$, while the volume growth rate and density decrease as $\propto \frac{1}{t}$, following factors that depend on $\Delta\lambda$, which explains the trend of the theoretical plots in Fig. 3ABD of the main text.

#### F. Model prediction on the relationship between dry-mass density and volume growth rate

By definition, the volume growth rate $\lambda_V$ is equal the logarithmic derivative, that is, $\lambda_V := \frac{1}{V}\frac{dV}{dt}$. Therefore, we obtain that the model predicts the following relationship:

$$\lambda_V = \left[\frac{\lambda_M}{w_{\mathrm{i}}'} + \frac{f_p(j_a - \lambda_M)}{w_{\mathrm{osm}}}\right]\rho \tag{S45}$$

This equation suggests that the volume growth is proportional to the dry-mass density if the factor $\left[\frac{\lambda_M}{w_{\mathrm{i}}'} + \frac{f_p(j_a - \lambda_M)}{w_{\mathrm{osm}}}\right]$ is kept constant. This is not obvious a priori. In our work, we plotted the volume growth rate against the density for different cells at different times during the bulk of interphase. Even if the model is correct, whether the proportionality appears in the data will depend on the cell-to-cell and time variability of the factor $\left[\frac{\lambda_M}{w_{\mathrm{i}}'} + \frac{f_p(j_a - \lambda_M)}{w_{\mathrm{osm}}}\right]$. Our observation of the proportionality for suspended cells suggests that this factor does not change enough to mask the linear relationship. In addition, we may reduce some of the variability by fixing the value of $\lambda_M$. As we expect $w_{\mathrm{i}}'$ and $w_{\mathrm{osm}}$ to be relatively constant (since they depend mainly on physical constants), the main source of variability should come from the parameter $j_a$.

Finally, we also write the expression of the volume growth to emphasize its dependence on the mass growth rate:

$$\lambda_V = -\left(\frac{f_p}{w_{\mathrm{osm}}} - \frac{1}{w_p}\right)\rho\,\lambda_M + \frac{f_P j_a}{w_{\mathrm{osm}}}\rho\,. \tag{S46}$$

In particular, this equation shows that, at fixed $\rho$, the sign of the slope relating the volume and mass growth rate is equal to the sign of the jump in the volume growth rate under inhibition of mass growth, see Eq. (S40). Unfortunately, this consistency relationship is not visible in our data, when considering the conditional average of $\lambda_M$ given $\lambda_V$ (see Fig 2E in the main text). This inconsistency is possibly due to technical limitations in the data or in the data analysis, or to the existence of yet uncharacterized correlated fluctuations in the two quantities whose origin is not included in our simplified model, pointing to the need for further investigation.

#### G. Different physiological regulatory responses restore the volume growth rate after translation inhibition

The previous considerations assumed that all parameters of the model describing the physiological state of the cell are kept constant under translation inhibition, with the obvious exception of the protein production flux $j_P$, which becomes zero. In particular, amino acid production remains unchanged and active indefinitely, which is only reasonable for short times. However, we have already discussed that Fig. 3BD of the main text falsify this scenario. Hence, the predictions of the osmo-metabolic model under these assumptions are expected to hold only until the cell has time to react. We now explore the consequences of relaxing these strict assumptions and describing the physiological response of the cell.

Let us consider again Eq. S37. Previously, the amino acid production flux $j_A$ was fixed as a constant parameter in time. We need to investigate what happens if $j_A$ becomes a function of time through some time-varying biologically relevant parameters and under few possible biologically relevant scenarios, which are not necessarily mutually exclusive.

Generally speaking, we can anticipate a reduction in $j_A$ when protein synthesis stops, for several reasons [1, 2, 8, 9, 23]. From a functional perspective, accumulating amino acids that are not being used for protein production may be considered unnecessary. Another reason might relate more directly to maintaining dry-mass density homeostasis. A rapid decrease in $j_A$ could help maintain the balance between mass growth and volume expansion, especially when protein synthesis is inhibited, allowing volume growth to slow in tandem with mass. On a mechanistic level, the reduction in $j_A$ following the cessation of protein production could occur due to feedback inhibition, where an accumulation of free amino acids down-regulates the activity of transporters and metabolic enzymes [23]. More broadly, sophisticated biological regulation of amino acid production and transport might play a role in responding to translation inhibition. Next, we will discuss three different mechanisms and their impact on how volume changes over time after a full inhibition of translation, which is an observable quantity in our data (Fig. 3BD in the main text).

***Scenario 1. Enzyme dilution decreases amino acid production under translation inhibition.***

As a first minimal hypothesis, we assume that $j_A$ could a be a function of the concentration of a class of protein enzymes that is responsible for amino acid transport and metabolism, that is $j_A = c[E]$ where $c$ is a constant factor. In this case, a simple mechanism that causes amino acid production to drop is the dilution of the enzyme concentration $[E]$. Since under full inhibition of translation proteins are not being produced anymore, the concentration $[E]$ becomes simply proportional to the inverse of the volume $V$.

Writing the equation for the volume after translation inhibition in light of these assumptions, we get

$$\frac{dV}{dt} = \frac{j_A\, f_P}{w_{\text{osm}}}\, M(t=0) = c[E]\, \frac{f_P}{w_{\text{osm}}}\, M(t=0) = \frac{c^{'}}{V}\,, \tag{S47}$$

where $c^{'} = cE(t=0)\, \frac{f_P}{w_{\text{osm}}}\, M(t=0)$. By solving this equation, we obtain

$$V(t) = V_0\, \sqrt{1 + 2[E_0]\frac{f_P\, \rho^*}{w_{\text{osm}}}\, t}\,. \tag{S48}$$

Interestingly, the volume does not increase linearly anymore as in the previous section, but as the square root of time after treatment $\propto \sqrt{t}$. However, the volume still grows indefinitely in this picture.

***Scenario 2. Enzyme degradation decreases amino acid production and leads to volume saturation***

If amino acid production depends on the concentration of metabolic enzymes $[E]$, a second rationale for a decreasing amino acid production is degradation of such enzymes in response to translation inhibition. The equation for the concentration $[E]$ can be generically expressed as

$$\frac{d[E]}{dt} = k - (\lambda_V + \eta)\, [E]\,, \tag{S49}$$

where $k$ is the production rate, while $\eta$ is the protein degradation rate. As translation halts, $k = 0$. In addition, we assume that degradation occurs faster than dilution when it is activated, that is, $\eta \gg \lambda$. In this case $\frac{d[E]}{dt} = -\eta\, [E]$, consequently

$$[E] = [E_0]\, e^{-\eta\, t}\,. \tag{S50}$$

As $j_A$ follows this concentration, this is equivalent to stating that

$$j_A = j_A^0\, e^{-\eta\, t}\,. \tag{S51}$$

In this case, the volume equation becomes

$$\frac{dV}{dt} = \frac{j_A\, f_P}{w_{\text{osm}}}\, M(t=0) = \frac{j_A^0\, f_P}{w_{\text{osm}}}\, M(t=0)\, e^{-\eta\, t}\,. \tag{S52}$$

Following the same steps as the previous section, we express the equation as

$$\frac{dV}{dt} = (\lambda^* + \Delta\lambda)\; V(t=0)\; e^{-\eta\, t}\,. \tag{S53}$$

Consequently, the prediction of the osmo-metabolic model for the time evolution of volume in this scenario becomes

$$V(t) = V_0 \left[1 + (\lambda^* + \Delta\lambda)\; \frac{(1 - e^{-\eta\, t})}{\eta}\right]\,. \tag{S54}$$

Hence in this scenario, the volume behaves linearly for sufficiently short timescales ($t \ll 1/\eta$). On longer timescales, protein degradation becomes relevant and the volume saturates to the constant value $V_0\left(1 + \frac{(\lambda^*+\Delta\lambda)}{\eta}\right)$. Therefore, volume does not grow indefinitely in this case. In this scenario, the level of dry mass dilution at the end of an ideal long experiment is also set by the dilution factor $\left(1 + \frac{(\lambda^*+\Delta\lambda)}{\eta}\right)$ which represents how much a cell would be diluted if protein production stops.

Eq. (S54) was assumed in the fits of the osmo-metabolic model with feedback used in Fig 3ABD of the main text. See the "Data Analysis" section of this document below for further details on the fitting procedure.

***Scenario 3. Metabolic enzyme down-regulation leads to sublinear volume growth.***

While in the previous section we have considered the case $j_A$ is proportional to the total concentration of enzymes $[E]$, here we generalize the previous case by considering the possibility that $j_A$ is proportional to the active concentration of enzymes $[E_{ac}]$. In this scenario, $j_A$ may go down due to progressive inactivation of such enzymes rather than dilution or degradation.

Rather than a specific scenario this is a set of many scenarios, since multiple mechanisms could be responsible for inactivation, including allosteric regulation, competitive binding, etc. However, generically we can claim that $j_A(t)$ needs to decrease to nearly zero due to regulation of the metabolic sector, e.g. by enzyme downregulation, sequestration or inactivation, in order to match the blocked translation flux. Hence, we generically expect that

$$\frac{dV}{dt} = \frac{j_A}{w_{\text{osm}}}\; M_P(t=0) \tag{S55}$$

or in other words that the volume will grow sublinearly with a slowing-down kinetics that depends on the details of the regulatory response.

An important consideration in this scenario is the timescale of enzyme inactivation, which determines how strictly volume and dry mass are coupled. If inactivation occurs rapidly, volume changes closely track dry mass alterations, resulting in minimal dilution effects during translation arrest or overgrowth. Consequently, fluctuations in protein translation rates would have little impact on dry mass density, as the volume adjusts to maintain coupling. This model, alongside experimental data, suggests that the coupling timescale between dry mass and volume is relatively long. One possible explanation could be that small density fluctuations are tolerable and can be corrected by the homeostatic mechanisms described in the model, leading to a broader range of acceptable dry mass densities rather than requiring a narrowly fixed density. This indicates that both physical and biological coupling mechanisms may operate on longer timescales, allowing for flexibility in maintaining cellular homeostasis without the need for adjustments on faster time scales.

#### 3. PHENOMENOLOGICAL MEAN-FIELD MODEL OF DRY-MASS DENSITY HOMEOSTASIS

This section describes the phenomenological model of dry-mass density homeostasis for the average-cell behavior used in Fig. 5 of the main text for the analysis of homeostatic behavior of cellular density. This model does not directly directly describe the evolution of the total mass and volume, but does so using the specific rates. Hence, mass and volume may be obtained by integration of the specific rates, and, roughly speaking, they depend exponentially on the rates $M \propto e^{t\,\lambda_M}$ and $V \propto e^{t\,\lambda_V}$. We verified that the basal average growth mode in mass and volume is exponential in our data (see Supplementary Figure S1JK), in line with previous studies [11, 24]

**The phenomenological mean-field model describes the dry-mass density dynamics by explicitly taking into account mass and volume dynamics** We begin by describing the behavior of the model in the bulk of the cell cycle. We recall (see Fig. 1BD of the main text and Methods) that that we define the "bulk" of the cell cycle as the time period where mass and volume growth rates are on average equal, thus excluding the period from mitotic entry to the end of the post-mitotic spreading (see Supplementary Figure 1B). The model is general because it describes the evolution of macromolecular dry-mass density $\rho$ using an equation derived solely from the definition of that density as the ratio of macromolecular dry mass to volume, readily obtained using the chain rule. Indeed, following the chain rule,

$$\frac{d\rho}{dt} = (\lambda_M - \lambda_V)\,\rho\,, \tag{S56}$$

where $\lambda_M$ and $\lambda_V$ can generally also vary with time through other state variables. Finally, the rates may be function of the dry-mass density, as shown by our data. In the simplest scenario, we may assume an instantaneous dependency $\lambda_M = \mu_M(\rho)$ and $\lambda_V = \mu_V(\rho)$ based on our results (see Fig. 3F and 4A of the main text. In this case, all three variables $\rho$, $\lambda_M$ and $\lambda_V$ directly depend on density. More generally, a change in the macromolecular dry-mass density may lead to a change in the growth rates over a relaxation time scale. In such case, the dependencies require a further time kernel. The equations under the assumption of a direct dependency are

$$\frac{d\lambda_M}{dt} = [\mu_M(\rho) - \lambda_M]\,\theta_M\,, \tag{S57}$$

and

$$\frac{d\lambda_V}{dt} = [\mu_V(\rho) - \lambda_V]\,\theta_V\,. \tag{S58}$$

**Differences with standard approaches.** Note that this approach differs from assuming an autonomous exponential volume growth, as done for example when describing gene expression dynamics. That approach provides a simplified perspective on cellular macromolecule dry-mass density dynamics by assuming constant production and dilution rates. This leads to a straightforward steady-state solution but overlooks the regulatory links between mass and volume observed in cells.

In contrast, the proposed model explicitly considers the dynamics of mass growth rate ($\lambda_M$) and volume growth rate ($\lambda_V$), which are treated as dynamic variables that can depend on dry-mass density and potentially on other factors. In this view density changes are a direct consequence of the difference between mass and volume growth rates, ($\lambda_M - \lambda_V$), allowing for a representation of cellular behavior through the phenomenological functions $\mu_M(\rho)$ and $\mu_V(\rho)$, describing density-dependent target mass and volume growth rates, respectively. This feedback loop provides a more accurate description of cellular density regulation, acknowledging that growth rates are not static but can be influenced by other factors. In particular, they can be functions of dry-mass density, reflecting our experimental observations.

#### 4. MEAN-FIELD MODEL OF DRY-MASS DENSITY HOMEOSTASIS: RESULTS

##### A. Conditions for dry-mass density homeostasis

This generic mean-field model provides some general conditions for dry-mass density homeostasis to be enforced (see Fig. 5 in the main text for the data inspiring these assumptions). Specifically, the conditions are the following:

1. there is a target value of the macromolecular dry-mass density $\rho^*$ such that a cell with such density maintains indefinitely this value in the absence of external perturbations ($\frac{d\rho}{dt}|_{\rho=\rho^*}$);
2. if the density $\rho$ of the cell is greater than $\rho^*$ then it decreases back to $\rho^*$ ($\frac{d\rho}{dt}|_\rho < 0$), while if the density $\rho$ of the cell is less than $\rho^*$ then the density increases back to $\rho^*$ ($\frac{d\rho}{dt}|_\rho > 0$). This must be true at least if $\rho$ is sufficiently close to $\rho^*$. More formally, our statement is equivalent to the following mathematical condition $\left(\frac{\partial}{\partial\rho}\frac{d\rho}{dt}\right)|_{\rho=\rho^*} < 0$.

In the language of dynamical systems theory, this is equivalent to stating that the dry-mass density dynamics has a stable fixed point $\rho^*$. Following Eq. S56 we shall define the conditions for the couplings between the growth rates and the density that guarantee the existence of a homeostatic value $\rho^*$ that satisfies these two conditions.

We first answer this question assuming that $\lambda_M = \mu_M(\rho)$ and $\lambda_V = \mu_V(\rho)$, or equivalently, that a change in the density is reflected instantaneously in a change in the growth rates. The two conditions above translate mathematically in the following two conditions for the growth rates,

1. $\mu_M(\rho^*) = \mu_V(\rho^*)$
2. $\left(\frac{\partial}{\partial\rho}\frac{d\rho}{dt}\right)|_{\rho=\rho^*} = \frac{\partial}{\partial\rho}\mu_M(\rho) - \frac{\partial}{\partial\rho}\mu_V(\rho)|_{\rho=\rho^*} < 0$ .

However, although quite general, these expressions are not very transparent. To provide an intuition on the meaning of these conditions, we consider the concrete case of linear couplings,

$$\mu_M(\rho) = a_M + b_M\,\rho\,, \tag{S59}$$

and

$$\mu_V(\rho) = a_V + b_V\,\rho\,, \tag{S60}$$

which is also the one supported by the data (see Fig. 3F, Fig. 4A and Fig. 5B in the main text). More specifically, the slope of $\mu_M(\rho)$ is always negative in our data (Fig. 4A), while as we have already commented above, the slope of $\mu_V(\rho)$ is positive for suspension cells and flat for adherent cells (Fig. 3FG), leading to slightly different model predictions for these two situations.

In case of linear dependencies, the second condition above translates into saying that $b_M < b_V$. In particular, homeostasis is guaranteed if

1. $b_M < 0$ and $b_V = 0$
2. $b_M = 0$ and $b_V > 0$
3. $b_M < 0$ and $b_V > 0$
4. $b_M > 0$ and $b_V > 0$ with $b_M < b_V$ ,

which is always the case in our data. Any other case does not lead to the existence of a homeostatic dry-mass density.

The homeostatic target density $\rho^*$ is predicted by the model to be exactly at the intersection between $\mu_M(\rho)$ and $\mu_V(\rho)$. Fig. 5DEF of the main text shows that this prediction is followed by the data of all three tested cell lines, regardless of whether they grow in suspension or adhesion. Intriguingly, the specific value of the target density $\rho^*$ is similar across all cell lines considered, that is, $\rho^* \approxeq 0.15$ pg $\mu$m$^{-3}$.

We also note that even in absence of strictly linear coupling, we may expand the growth rate around $\rho^*$ and obtain the same condition on the linear expansion $\mu_M(\rho) \simeq \lambda^* + b_M\,(\rho^* - \rho)$ and $\mu_V(\rho) \simeq \lambda^* + b_V\,(\rho^* - \rho)$.

Finally we also note that the slope parameters $b_V$ and $b_M$ may be re-interpreted as the range of growth rates that can be obtained by modulating the density, as $b_V = \Delta\lambda_V/\Delta\rho$ and $b_M = \Delta\lambda_M/\Delta\rho$ given the linear assumption. In particular, in our data the density ranges roughly from 0.1 to 0.2 pg ¯m$^3$, that is the density has a 2-fold range. Then, we may ask what is the corresponding range of growth rates $\Delta\lambda$. In our data, this partly depends on the cell type, but overall in range between 0 and 0.03 h$^{-1}$ depending on cell type and whether the mean the mass or the volume growth rate. Therefore, growth rates can change at most with a factor of 2 from the equilibrium value (which is 0.03 h$^{-1}$ roughly in hour data).

#### B. The dependency of the rate of change of dry-mass density from dry-mass density itself reflects the strength of density homeostasis implemented by growth-rate changes

Another generic prediction of this mean-field phenomenological model is that the slope of the relationship $\frac{d\rho}{dt}$ vs $\rho$ is a direct reflection of the strength of the homeostatic coupling between growth and density. This prediction is tested in Fig. 5GH of the main text, and shows very good agreement with the data.

Let us now describe how this prediction is obtained. From Eq. (S56), if $\rho$ is sufficiently close to the target density $\rho^*$, $\lambda_M = \mu_M(\rho)$ and $\lambda_V = \mu_V(\rho)$, we obtain by expanding to first order around $\rho^*$

$$\frac{d\rho}{dt} = -\left(\frac{\partial}{\partial\rho}\mu_V(\rho)|_{\rho=\rho^*} - \frac{\partial}{\partial\rho}\mu_M(\rho)|_{\rho=\rho^*}\right)\rho^*\,. \tag{S61}$$

Therefore, the slope of $\frac{d\rho}{dt}$ vs $\rho$ reflects the rate of change of the growth rates with respect to the dry-mass density near the homeostatic value. The more sensitive the growth rates are to changes in density, the steeper the slope and the faster the restoring response leading to the homeostatic density. For the particular case of linear coupling, the equation reduces to

$$\frac{d\rho}{dt} = -\left(b_V - b_M\right)\rho^* \left(\rho - \rho^*\right) . \tag{S62}$$

The slope of $\frac{d\rho}{dt}$ vs $\rho$ around $\rho^*$ is therefore $(b_V - b_M)\,\rho^*$. If $b_M < 0$ this equation is particularly transparent, since the slope is just the sum of the absolute slope of the growth rates with the dry-mass density, multiplied by the steady state value of the density.

In our data, the value of the slope of the relationship $\frac{d\rho}{dt}$ correlates very well with $(b_V - b_M)\,\rho^*$ as shown by Fig. 5H in the main text. The overall values of the slope range from 0.03 to 0.06 $h^{-1}$ depending on the particular cell lines. It is worth noting that this is roughly the value of the steady-state growth rate. Therefore, the relaxation timescale of the dry-mass density is set by the value of the steady state growth rate. *A priori*, this did not need to be the case. According to the model, the growth rates vary more strongly with density, $b_V$ and $b_M$ would be greater and the relaxation timescale would be faster. Instead, our data shows that growth rates vary 2-3 fold at most with density and this fact implies that the relaxation timescale of the density is roughly equal to the inverse growth rate.

#### 5. STOCHASTIC SINGLE-CELL MODEL OF DRY-MASS DENSITY HOMEOSTASIS

This section illustrates the single-cell phenomenological model of dry-mass density homeostasis under noisy growth employed in Fig. 6 of the main text. The model expands the phenomenological model of the previous sections, and describes the dynamics of three fluctuating quantities, the mass growth rate $\lambda_M$, the volume growth rate $\lambda_V$ and the dry-mass density $\rho$, assuming that the former two quantities are coupled to the latter following our empirical observations in Fig. 3 and 4, and that the mean behavior is described by Eq. (S56). As previously, the model does not directly describe the evolution of the total mass and volume, but these quantities may be obtained by (stochastic) integration of the fluctuating rates. However, we note that our primary goal when using this model is to describe the behavior of density fluctuations, not fluctuations in absolute mass and volume.

**The stochastic model describes the interplay of noise with mass- and volume-based homeostasis in single cells.** The model is illustrated in Fig. 6A of the main text. As in Eq. (S56), the dry-mass density evolves with an equation that depends only on the definition of dry-mass density as the ratio of dry mass over volume. However, in this stochastic model $\lambda_M$ and $\lambda_V$ are fluctuating variable due to noisy growth. Specifically, we assume they both evolve according to a modified Ornstein–Uhlenbeck process [25]. In a stochastic process of this kind, a fluctuating variable tends to relax toward its average value $\mu$ with a timescale $\theta$, but it is also subjected to noisy variation at each timestep with noise parameter $D$. In our modified version, both growth rates obey this dynamics but the average value depends on the instantaneous density value, that is, $\mu(\rho)$. This reflects the presence of a coupling between the growth rates and the density. Mathematically, we write the following equations

$$\frac{d\lambda_M}{dt} = \left[\mu_M(\rho) - \lambda_M\right]\theta_M + D_M\,\xi(t) , \tag{S63}$$

and

$$\frac{d\lambda_V}{dt} = \left[\mu_V(\rho) - \lambda_V\right]\theta_V + D_V\,\nu(t) . \tag{S64}$$

where the white noise tems $\xi(t)$ and $\nu(t)$ are independent realizations of the derivative of a Wiener process.

We note that according to these equations, mass and volume growth rates fluctuate around a mean value, respectively ($\mu_M(\rho)$ and $\mu_V(\rho)$) that depends on the dry-mass density, which in turn obeys Eq. S56. To describe growth in the bulk of the cell cycle (when mass and growth rates are equal, see Fig. 6 in the main text, the following conditions must be verified,

$$\langle\lambda_M\rangle = \langle\mu_M(\rho)\rangle\,; \qquad \langle\lambda_V\rangle = \langle\mu_V(\rho)\rangle\,; \qquad \langle\lambda_M\rangle = \langle\lambda_V\rangle\,, \tag{S65}$$

where $\langle\cdot\rangle$ represents expected values. If $\mu_M$ and $\mu_V$ are linear functions of the dry-mass density $\rho$, which is a good approximation in our data, we may also express these conditions as

$$\langle\lambda_M\rangle = \mu_M(\langle\rho\rangle) \ ; \qquad \langle\lambda_V\rangle = \mu_V(\langle\rho\rangle) \ ; \qquad \mu_M(\langle\rho\rangle) = \mu_V(\langle\rho\rangle) \ , \tag{S66}$$

where the final equation defines the average steady-state density, as its solution is a value of $\langle\rho\rangle$ such that $\frac{d\langle\rho\rangle}{dt} = 0$.

Therefore, the model is consistent with a steady state for the average dry-mass density, in accordance with the mean-field model presented previously. In addition, the model can generate single-cell tracks and therefore formulate predictions regarding the cell-to-cell variability and the dynamics of growth fluctuations. We note that in order to simulate the model we need to specify some initial conditions $\rho_0$, $\lambda_{M0}$ and $\lambda_{V0}$. In the main text, we fix the initial conditions directly from the data (see Methods). These initial conditions are useful to describe the effect of the perturbations occurring around division (see below), but we initially focus on the bulk phase of the cell cycle.

**The period across a cell division is treated as a perturbation.** We tailored this part of the model to key observations regarding our data (see main text Fig. 1). We note that outside of the bulk phase of the cell cycle (i.e. during mitosis and post-mitotic spreading), the homeostatistic relationship between the growth rates and the dry-mass density are broken (see main text). In addition, no apparent homeostatic mechanism is present for the density (Supplementary Fig. S6AB). Specifically, before cell division cells are diluted because of the arrest of macromolecular dry-mass growth and of the mitotic swelling, ans subsequently concentrated due to the post-mitotic spreading (Fig. 1 of the main text). As a consequence of these processes, after spreading, the cells start the bulk phase of the cell cycle with a dry-mass density that is weakly correlated with the one they had in the previous generation before mitosis.

Hence, in order to focus on the effect of these changes to the bulk phase of the cell cycle (see Supplementary Fig. 1BD), we treat the period from onset of mitosis to spreading in the subsequent generation as a perturbation on density. Specifically, if $t_f$ is the time where the bulk cell-cycle period ends and $t_0$ is beginning of the new post-spreading bulk period in the following generation, we assume that the dry-mass density at time $t_0$ obeys the following equation,

$$\rho(t_0^{n+1}) = \rho(t_f^n) + \Delta_\rho \ , \tag{S67}$$

where $\Delta_\rho$ is an independent random variable, assumed Gaussian, with zero mean and whose variance may be fixed from data, which models the fact that the period outside of bulk growth is not homeostatic. Panel A and B in Supplementary Fig. S6 shows the densiy indeed obeys this equation, as the increment $\rho(t_0^{n+1}) - \rho(t_f^n)$ behaves as an uncorrelated random variable. For the growth rates, we write, along the same lines,

$$\lambda_M(t_0^{n+1}) = \lambda_M(t_f^n) + \Delta_{\lambda_M}, \tag{S68}$$

and

$$\lambda_V(t_0^{n+1}) = \lambda_V(t_f^n) + \Delta_{\lambda_V} \ . \tag{S69}$$

**Behavior of the model across multiple cell cycles.** Putting together the previous two model components, we come to the full picture of the stochastic single-cell growth model. Starting from the beginning of a bulk period, the model follows Eqs. (S63, (S64) and (S56), which predict average-density homeostasis under a suitable form of the function $\mu_M(\rho)$ and $\mu_V(\rho)$. The period across division is not explicitly described, and at the beginning of the subsequent bulk period the density and growth rates are partially randomized from mother to daughter. The process can be iterated along arbitrarily long lineages in order to predict the long-term behavior of the fluctuations, specified by the growth-rate and dry-mass density distributions.

In summary, if we call $n$ the cell-cycle index along a lineage, the process is described by the following equations,

$$\frac{d\lambda_M}{dt} = [\mu_M(\rho) - \lambda_M\,]\,\theta_M + D_M\,\xi(t) \ , \tag{S70}$$

$$\frac{d\lambda_V}{dt} = [\mu_V(\rho) - \lambda_V\,]\,\theta_V + D_V\,\nu(t) \ , \tag{S71}$$

$$\frac{d\rho}{dt} = (\lambda_M - \lambda_V)\,\rho \ , \tag{S72}$$

for the bulk phases, and

$$\rho(t_0^{n+1}) = \rho(t_f^n) + \Delta \,, \tag{S73}$$

$$\lambda_M(t_0^{n+1}) = \lambda_M(t_f^n) + \Delta_{\lambda_M} \,, \tag{S74}$$

$$\lambda_V(t_0^{n+1}) = \lambda_V(t_f^n) + \Delta_{\lambda_V} \,, \tag{S75}$$

describing the transition between bulk cell-cycle phases of subsequent generations.

#### 6. STOCHASTIC SINGLE-CELL MODEL OF DRY-MASS DENSITY HOMEOSTASIS: RESULTS

This section describes in more detail the main predictions provided by the model. The reader is referred to Fig. 6 of the main text for a summary of these results.

##### A. The model predicts controlled density fluctuations at each cell cycle and along lineages.

Using the stochastic model, we established under which conditions the system achieves a stable dry-mass density distribution (see Fig. 6C in the main text).

We note that the given the quasi-periodic perturbations across generations, the model does not attain a stationary probability distribution. However, we can still ask whether the time-dependent cell-to-cell variability observed during each cell cycle does not spread. To this end, given the density distribution $P(\rho, t)$ across cells during the bulk period, we quantified the long-term fluctuations as proxied by the standard deviation of this distribution. In general, this is an oscillatory function, but its range remains bounded, and its time average as defined in the following equation is finite,

$$(\sigma_\rho^\infty)^2 = \lim_{t\to\infty} \sigma_\rho^2(t) = \lim_{t\to\infty} \frac{1}{T} \int_0^t dt \int (\rho - \langle \rho(t) \rangle)^2 \, P(\rho, t) \, d\rho < \infty \tag{S76}$$

An analytical calculation of this quantity for the general model is difficult. The results in Fig. 6C derive from stochastic computer simulations of the trajectories $\rho(t)$ given by the model from which we derive $\sigma_\rho(t)$ for different combinations of parameters with a focus on three qualitatively different scenarios, as follows:

- no growth noise, $D_M = D_V = 0$, and homeostatic coupling in the functions $\mu_M(\rho)$ and $\mu_V(\rho)$;
- no homeostatic coupling in the function in $\mu_M(\rho)$ and $\mu_V(\rho)$, for instance a costant function independent of density, and some growth noise, that is, $D_M > 0$ and $D_V > 0$;
- both growth noise and homeostatistic coupling are present.

**A simplified analytical model gives key insights on the existence of controlled dry-mass density fluctuations.** Additionally, provide insight on the different scenarios, we have worked with analytical calculations of a simplified version of the stochastic single-cell model. In order to simplify the picture, we asusme an effective model for only the dry-mass density variable $\rho$ (which is ultimately our observable of interest), as follows,

$$\frac{d\rho}{dt} = -k(\rho - \rho^*) + D_\rho \xi(t) \,, \tag{S77}$$

where we suppose that the parameter $k$ and $D_\rho$ are function of the original parameters governing the growth dynamics,

$$k = f(\mu_M, \mu_V) \qquad D = g(D_V, D_M). \tag{S78}$$

Additionally, across cell divisions, density is perturbed by the following prescription,

$$\rho(t_0^{n+1}) = \rho(t_f^n) + \Delta \qquad t_f^n - t_0^n = \tau \,, \tag{S79}$$

where $Delta$ is an independent Gaussian variable with mean 0 and variance $\sigma_0^2$.

Using this simplified model, we can estimate the asymptotic fluctuation $\sigma_\rho$ analytically. Assuming that we start after cell spreading in a certain cell cycle and the dry-mass density is initially distributed with a standard deviation $\sigma_0$, we can solve Eq. (S77) to obtain an equation for the fluctuation at the end of the bulk cycle. As we aim to understand the behavior after many cycles, we denote the standard deviation of the density after the first cycle using the subscript 1.

$$(\sigma_1^f)^2 = \frac{D^2}{2k} + \left(\sigma_0^2 - \frac{D^2}{2k}\right) \exp(-2k\tau) \,. \tag{S80}$$

We can rewrite this equation using the notation $(\sigma_1^i)^2 := \sigma_0^2$, recognizing that the variance at time 0 is equal to the variance at the beginning of the first cycle. This notation will help us generalize the equation.

After a bulk cycle, the model follows Eq. S79. Since the perturbation is completely independent from the state of the cell at the end of the bulk cycle, the standard deviation at the beginning of the second cycle becomes

$$(\sigma_2^i)^2 = (\sigma_1^f)^2 + \sigma_\Delta^2 \,. \tag{S81}$$

We can substitute the expression for $(\sigma_1^f)^2$ and obtain

$$(\sigma_2^i)^2 = \frac{D^2}{2k} + \left[(\sigma_1^i)^2 - \frac{D^2}{2k}\right] \exp(-2k\tau) + \sigma_\Delta^2 \,. \tag{S82}$$

Next, we recognize that a similar equation can be derived for any cycle, not just the first and second cycles. This leads to a general recursive equation

$$(\sigma_{n+1}^i)^2 = \frac{D^2}{2k} + \left[(\sigma_n^i)^2 - \frac{D^2}{2k}\right] \exp(-2k\tau) + \sigma_\Delta^2 \,. \tag{S83}$$

To better understand the behavior as $n \to \infty$, we can rewrite this equation as a finite difference equation:

$$\Delta(\sigma_{n+1}^i)^2 = -(\sigma_n^i)^2 \left[1 - \exp(-2k\tau)\right] + \frac{D^2}{2k} \left[1 - \exp(-2k\tau)\right] + \sigma_\Delta^2 \,. \tag{S84}$$

Setting the left-hand side to zero gives us the solution for the asymptotic value of the variance at the beginning of the bulk cycle,

$$\sigma_\infty^i = \frac{D^2}{2k} + \frac{\sigma_\Delta^2}{1 - \exp(-2k\tau)} \,. \tag{S85}$$

This equation describes the long-term behavior of the fluctuations in terms of the biological parameters $D$, $\sigma_\Delta$, and $k$, which represent noise due to growth, noise due to processes across divisions (mitosis and spreading), and the strength of homeostatic control, respectively. From this equation, we can draw the following conclusions:

1. Any positive value of $k$ is sufficient to stabilize the fluctuations in the long term, meaning that even weak homeostatic control ensures the existence of finite fluctuations.

2. Without a homeostatic mechanism (i.e., $k = 0$), the fluctuations diverge.

3. In the absence of noise ($D = \sigma_\Delta = 0$), the fluctuations converge to zero.

4. The contribution of the mitotic perturbation to the long-term fluctuations depends on the length of the bulk cycle $\tau$.

Our simulation results in Fig. 6C, SFGH indicate that these findings hold true in a more complex model where growth rates also fluctuate, at least for realistic value of their fluctuations, with respect to our data.

##### B. The lifetime of dry-mass density fluctuations is set by the amount of growth variation with dry-mass density

Having discussed the factors that determine the size of fluctuations in our model, we now turn to understanding the lifetime of dry-mass density fluctuations, as illustrated in Fig. 6D of the main text. We go back to the full model and we begin by examining the zero noise limit where the growth rates are deterministic functions of the dry-mass density, i.e., $\lambda_M = \mu_M(\rho)$ and $\lambda_V = \mu_V(\rho)$ (which corresponds to the mean-field model),

$$\frac{d\rho}{dt} = (\mu_M(\rho) - \mu_V(\rho))\rho \,. \tag{S86}$$

Assuming that the coupling between growth rates and dry-mass density leads to a homeostatic density $\rho^*$ where $\mu_M(\rho^*) = \mu_V(\rho^*)$, we can analyze the behavior near this homeostatic density,

$$\frac{d}{dt}(\rho - \rho^*) = \left(\frac{\partial}{\partial \rho}\mu_M - \frac{\partial}{\partial \rho}\mu_V\right)\Big|_{\rho=\rho^*} (\rho - \rho^*)\rho^* \,. \tag{S87}$$

Let us define $k := \left(\frac{\partial}{\partial \rho}\mu_M - \frac{\partial}{\partial \rho}\mu_V\right)\big|_{\rho=\rho^*}$, then we have:

$$(\rho(t) - \rho^*) = (\rho(0) - \rho^*)\ \exp(-k\,\rho^* t)\,. \tag{S88}$$

The exponential decay of the dry-mass density fluctuation $(\rho(t) - \rho^*)$ shows that such fluctuations are damped, with a single characteristic timescale associated with the relaxation process. The parameter $k$ plays a crucial role in determining the rate of decay, as it quantifies the sensitivity of the growth rates to changes in density near the homeostatic point. The characteristic timescale $\tau_c$ is the inverse of the product of $k$ and $\rho^*$

$$\tau_c = \frac{1}{k\,\rho^*} \tag{S89}$$

This timescale represents the time it takes for the system to return to its homeostatic density after a perturbation. A smaller value of $\tau_c$ indicates faster relaxation, while a larger value implies slower relaxation. This relationship highlights the importance of the homeostatic density $\rho^*$ and the sensitivity parameter $k$ in determining the lifetime of density fluctuations in the model.

Thus, an initial fluctuation $(\rho(0) - \rho^*)$ from the homeostatic value decays back to zero with a characteristic timescale $\tau_{life} := \frac{1}{k\,\rho^*}$. In the case of linear coupling between growth and death rates, this timescale can be interpreted more intuitively. If the growth rates follow linear relationships with the dry-mass density, as observed on average in our data,

$$\mu_M(\rho) = a_M + b_M\,\rho \tag{S90}$$

$$\mu_V(\rho) = a_V + b_V\,\rho, \tag{S91}$$

we can express the characteristic decay timescale of density fluctuations as

$$\tau_c := \frac{1}{(b_M - b_V)\,\rho^*}\,. \tag{S92}$$

In this scenario, the slopes $b_M$ and $b_V$ represent the sensitivity of the growth and death rates to changes in dry-mass density. These slopes can be approximated roughly by the ratio of the growth rate range to the density range, e.g., $b_M \sim \Delta\lambda_M/\Delta\rho$. Consequently, $\tau_c$ measures approximately the timescale required for the cell to coordinate changes in growth rates in response to variations in density (hence, a higher sensitivity of growth and death rates to density changes leads to faster decay of density fluctuations). We show that this relationship holds in the full model in Supplementary Fig. 6L, where the dashed black line represents the prediction of Eq. (S92) and the symbols represent direct simulation of the model.

In presence of noise, this response time is measured by the autocorrelation function of the dry-mass density in absence of perturbations. If $\mu_V(\rho)$ and $\mu_M(\rho)$ fluctuate, the situation is more complex to approach analytically, as they provide a multiplicative noise. However we can provide a simple argument based on the simplified analytical model used above. Since Eq. (S77) defines an Ornstein-Uhlenbeck process [25], for which the decay of the mean upon a perturbation and the one of the autocorrelation coincide, hence we expect that $\tau_c$ is also the autocorrelation time.

We have tested this prediction and Eq. (S92) with simulations of the full model, and they are verified for parameters derived from our data (See Supplementary Fig. S6K and L).

Fig. 6D shows results from simulations of the full model with parameters taken from our data, and shows that the measured relaxation times correspond to the expectation from Eq. (S92). Ideally we would need to measure the autocorrelation time in data and compare it to these relaxation times. In practice however, the measured relaxation times (2-3 cell cycles) are longer than the characteristic times at which perturbations (from mitosis, division and spreading) determine new perturbed initial conditions. Hence, the system is never stationary, and determining autocorrelations for density fluctuations from data is very challenging.

##### C. Cells maintain a strong identity over one cell cycle

As we have seen, the relaxation times in our data (Fig. 6D of the main text and Supplementary Fig. S6L), both measured directly and estimated from Eq. (S92), are longer than a bulk phase of the cell cycle (from end of spreading to mitosis). The former is estimated to be ≊ 29 h, while the latter takes ≈ 15 h. Additionally, the density perturbations felt across mitosis, cell division and spreading, give rise to uncorrelated initial densities in subsequent generations, as shown in Supplementary Fig. S6AB. Hence, we expect that over a cell cycle the cells maintain considerable diversity in their individual densities, but that along lineages these cell-specific densities are "shuffled" by the perturbations around cell division, which is indeed what we observe in our data. This subsection provides a more detailed description of how single-cells preserve their individuality over cell-cycle time scale, despite the presence of a target density, as illustrated in Fig. 6D of the main text.

We start by mathematically defining the time average and the ensemble average as follows

$$\bar{\rho}_i := \frac{1}{N_{\text{track}}} \sum_{i=1}^{N_{\text{track}}} \rho_i(t_j), \quad j = 1, \ldots, N_{\text{track}}; \quad i = 1, \ldots, N_{\text{cell}} \,. \tag{S93}$$

$$\langle \rho(t_n) \rangle := \frac{1}{N_{\text{cell}}} \sum_{i=1}^{N_{\text{cell}}} \rho_i(t_j), \quad j = 1, \ldots, N_{\text{track}}; \quad i = 1, \ldots, N_{\text{cell}} \,. \tag{S94}$$

It is important to note that $N_{\text{cell}}$ represents the number of cell tracks in the dataset, whereas $N_{\text{track}}$ denotes the number of timepoints within a track.

In addition to the time and ensemble averages, we define the respective deviations from these averages

$$\sigma_{\bar{\rho}} = \sum_{i=1}^{N_{\text{cell}}} \left( \rho_i(t_j) - \bar{\rho}_i \right)^2 \,, \tag{S95}$$

and

$$\sigma_{\langle \rho \rangle} = \sum_{i=1}^{N_{\text{cell}}} \left( \rho_i(t_j) - \langle \rho(t_j) \rangle \right)^2 \,. \tag{S96}$$

We evaluated these quantities computationally by simulating the trajectories of the model, and compared them to the data. Fixing the parameters by the considerations presented above we were able to make quantitative predictions as well as qualitative ones. For HeLa adherent cells, we use equilibrium growth rate and dry-mass density $\lambda^* = 0.03\ \text{h}^{-1}$, $\rho^* = 0.15\ \frac{\text{pg}}{\mu\text{m}^3}$, coupling constant $b_M = -0.23\ \frac{\mu m^3}{\text{pg}}\text{h}^{-1}$ and $b_V = 0$, while $a_M = -b_M\rho^* + \lambda^*$ and $a_V = \lambda^*$. Finally, we have set $\theta_M = \theta_V = 1\ \text{h}^{-1}$, $D_M = D_V = 0.02\ \text{h}^{-3/2}$, $\Delta = 0.022\ \frac{\text{pg}}{\mu\text{m}^3}$ and $\Delta_{\lambda_M} = \Delta_{\lambda_V} = 0$ (see Data analysis section below). The results, shown in Fig. 6EF of the main text, indicate that the deviations from the ensemble average are 3-folder higher than the fluctuations of each density track around theoretical trends, as predicted by the model.

##### D. The model predicts fluctuations across lineages

In addition to the trends within the cell cycle, the model can also predict trends across cell cycles, i.e., along long lineages. To explain this behavior, we remind that in the model mitosis and spreading at the end and start of the cell cycle act together as a consecutive period where dry-mass density is perturbed from its homeostatic value, while at the same time no homeostatic mechanism is present in this period (see Supplementary Fig. S6A). Therefore, the CV of the density is predicted to rise quickly from mother to daughter. Conversely, after this perturbation the homeostatic mechanisms become active and therefore the CV of the density should decrease. As discussed in the previous section, the time scale of this decrease is set in the model by the parameters $b_M$ and $b_V$, and is necessary for the dry-mass density to be homeostatic at all. Putting all these considerations together, the CV of the density should show sawtooth-like oscillations, as it increases quickly between mother and daughter, then decreases until the next mitosis etc. Our data only comprises short lineages of one generation. For such short lineages, the model can still predict how density fluctuations behaves from mother to daughter, as shown in Fig. 6B in the main text. Such short lineages provide evidence of cycling of the CV of density, which increases around mitosis and then decreases in the bulk.

While our data lacks longer lineages comprising of several cell cycles, we can still use it to predict the long-term trends based on the observed parameters. Specifically, the model can predict the long-term trend of the fluctuations as a function of the noise level and coupling strength, as shown in Fig. 6C in the main text (CV) and Fig. S6H (standard deviation). Note that Fig. S6G shows a sanity check the long-term trend of the average density, which is constant. Supplementary Fig. 6F shows the cell cycle average of the noise level, without the cell cycle fluctuation due to noise at division and mitosis. Conversely, Fig. S6E focuses on the cell cycle fluctuations in the case where both noise and coupling are present. The long-term trend of the fluctuations may be tested empirically in future long-term experiments. Conceptually, this result highlights that even a weak coupling between growth and dry-mass density is sufficient to stabilize a density range over the long term, despite of the periodic perturbations provided by cell division. At the same time, the density fluctuations would diverge without any enforcement of dry-mass density homeostasis, as shown by the red curve in Fig. 6C in the main text, which supports the idea that the restore mechanisms discovered here are necessary. In other words, the model, with the parameters extracted from the data, addresses an issue that is difficult to access directly from our data: the importance of the homeostatic coupling to stabilize the long-term distribution of density in the cycling population, given the levels of noise measured from the data.

#### 7. MODEL-GUIDED DATA ANALYSIS AND MODEL-DATA COMPARISONS.

This section describes in more detail our data analysis procedures.

##### A. Fit of the osmo-metabolic model

Here, we describe the fitting procedure used in Fig 3ABD of the main text. The basic osmo-metabolic model predicts the dynamics of the volume under inhibition of translation from two parameters, the equilibrium growth rate $\lambda^*$ and the jump in the growth rate $\Delta\lambda$ immedialy after inhibition. We note that the latter is a composite effective parameter that is itself a function of microscopic parameters.

To extract $\lambda^*$, we use the steady-state growth rate of the untreated HeLa cells along the cell cycle (the 'bulk' phase, see Supplementary Fig. 1). To extract $\Delta\lambda$, we fit the predicted curve of the normalized volume $V/V_0$ using equation S42, which is a simple linear function whose slope is $\lambda^* + \Delta\lambda$:

$$\frac{V(t)}{V(t=0)} = \begin{cases} e^{t\lambda^*} & t < 0 \\ [1 + t\ (\lambda^* + \Delta\lambda)] & t \geq 0\ . \end{cases} \tag{S97}$$

We used the "curvefit" function from the scipy.optimize library in python to extract the slope and then subtract the found value of $\lambda^*$.

In the more advanced osmo-metabolic models, the cellular response under inhibition of translation includes a physiological regulatory response (see Section 2E). The simplest version of these models that is compatible with the data includes an extra parameter $\eta$ such the normalized volume $V/V_0$ behaves according to equation S54:

$$\frac{V(t)}{V(t=0)} = \begin{cases} e^{t\lambda^*} & t < 0 \\ \left[1 + (\lambda^* + \Delta\lambda)\ \frac{(1-e^{-\eta\, t})}{\eta}\right] & t \geq 0\ . \end{cases} \tag{S98}$$

As described above, we extract $\lambda^*$ from the steady-state growth rate of the untreated HeLa cells along the cell cycle. We extract $\Delta\lambda$ and $\eta$ from the time series of $\frac{V(t)}{V(t=0)}$ using the curvefit function of scipy.optimize from python with the form indicated by the previous equation.

##### B. Tests of the mean-field model

Here, we describe how we tested the mean-field model in Fig. 6. We tested the mean field model by comparing the average of the growth rates conditioned to dry-mass density with the average of the density derivative conditioned to density. Let us define the symbol $\langle Y(X)\rangle$ for any conditional average of $Y$ as a function of fixed $X$. In our data analysis, we obtain such conditional average by plotting a scatter plot $Y$ vs $X$, taking a bin with a fixed number of points around $X$ and averaging $Y$ over that bin.

To a first approximation, the conditional average of the growth rates at a given density can be approximated by a simple linear relationship. We use the following curve to fit them

$$\langle \lambda_M(\rho)\rangle = \lambda^* + b_M\ (\rho^* - \rho) \tag{S99}$$

$$\langle \lambda_V(\rho) \rangle = \lambda^* + b_V \ (\rho^* - \rho) \tag{S100}$$

Under this approximation, the model predict that the the conditional average of the dry-mass density derivative at fixed density itself should take the form

$$\left\langle \frac{d\rho}{dt}(\rho) \right\rangle = -\left[ (b_V - b_M)\,\rho^* \right] (\rho^* - \rho)\,. \tag{S101}$$

Therefore, the slope of relationship between $\frac{d\rho}{dt}$ and $\rho$ should be predicted by the slopes of the conditional average of the growth rates at fixed dry-mass density.

##### C. Simulation of the stochastic single-cell mode

We used the Euler–Maruyama method to simulate the stochastic differential equations S63 and S63, while we simulate the dry-mass density equation S72 with the Euler algorithm.

##### D. Fitting of the stochastic single-cell model

The model presents nine external outputs, the functions $\mu_M(\rho)$ and $\mu_V(\rho)$, the parameters $\theta_M$, $\theta_V$, $D_M$, $D_V$ describe the noise response and level during the bulk phase of growth and finally the parameters $\Delta$, $\Delta_{\lambda_M}$, $\Delta_{\lambda_V}$ that describe the noisy effect of mitosis.

First, we fit the function $\mu_M(\rho)$ and $\mu_V(\rho)$ with a linear function from the curves in Fig. 5).

Sunsequently, we fit $\theta_M$ and $\theta_V$ using the conditional means $\langle \lambda_M(t+\tau) | \rho(t), \lambda_M(t) \rangle$ and $\langle \lambda_V(t+\tau) | \rho(t), \lambda_V(t) \rangle$ for short enough $\tau$ as the equations of the model imply that:

$$\theta_M \approx \frac{\langle \lambda_M(t+\tau) | \rho(t), \lambda_M(t) \rangle}{\mu_M(\rho(t)) - \lambda_M(t)}\, \tau \tag{S102}$$

$$\theta_V \approx \frac{\langle \lambda_V(t+\tau) | \rho(t), \lambda_V(t) \rangle}{\mu_V(\rho(t)) - \lambda_V(t)}\, \tau \tag{S103}$$

We also note that in an Ornstein–Uhlenbeck model the standard deviation of the mass and volume growth rate obey the following relationship at stationary state, $\sigma_M = \sqrt{\frac{D_M}{\theta_M}}$ and $\sigma_V = \sqrt{\frac{D_V}{\theta_V}}$. We measure in the data these variables from the empirical distribution of the growth rate and infer $D_M$ and $D_V$ given the value of $\theta_M$ and $\theta_V$ found before.

Finally, we fit the parameters $\Delta$, $\Delta_{\lambda_M}$, $\Delta_{\lambda_V}$ by comparing the distribution of the dry-mass density and growth rate before and after mitosis. The square root of these parameters is equal to the difference between the variance of these distributions.
